# Particle-based representation of phytoplankton modulates emergent coexistence and biogeography in a model ocean

**DOI:** 10.64898/2026.10.30.763182

**Authors:** Paul Frémont, Maurizio Ribera d’Alcalà, Bruno Hay Mele, Olivier Jaillon, Joshua S. Weitz, Daniele Iudicone

**Affiliations:** Department of Biology, University of Maryland, College Park, MD, USA; Department of Integrative Marine Ecology, Stazione Zoologica Anton Dohrn, Villa Comunale, 80121, Naples, Italy; Department of Biology, University of Naples “Federico II”, 80126, Naples, Italy; Génomique Métabolique, Genoscope, Institut François Jacob, CEA, CNRS, Université d’Évry, Université Paris-Saclay, Évry-Courcouronnes, France; University of Maryland Institute for Health Computing, North Bethesda, MD, USA; Department of Physics, University of Maryland, College Park, MD, USA

## Abstract

Phytoplankton communities sustain the marine food web through primary productivity. These communities sustain a high diversity of species, all of which must draw from the same finite nutrient supply. The mechanisms driving phytoplankton biogeography and succession, as well as those enabling high diversity, remain incompletely understood. Here, we compare emerging patterns of biogeography, richness, and diversity in a size-structured phytoplankton community as it is advected and diffused within an idealized North Atlantic double-gyre circulation, using both Eulerian and Lagrangian (particle-based) modeling (PBM) frameworks. When a sufficient number of particles is modeled, the PBM, which resolves mortality at the particle level, produces stronger seasonal variability in biomass in the northern gyre and less stable and higher biomass in the southern gyre, while reproducing large-scale chlorophyll patterns observed by satellite. These contrasting dynamics are accompanied by distinct regional diversity patterns: local richness and diversity are generally higher in the PBM than in the Eulerian model in the northern gyre, with differences depending on the mortality threshold, whereas in the oligotrophic southern gyre, PBM local richness increases with particle number, while Eulerian richness is more sensitive to the mortality threshold. The PBM also produces a dynamic and emergent biogeographical structure that is more robust to the mortality threshold than the Eulerian model. Finally, Lagrangian variability in environmental conditions experienced by particles shows a stronger impact compared to the local Eulerian variability in the Northern gyre and stresses variable sensitivity to environmental histories. Together, these results suggest that the discrete representation of biomass fundamentally alters mortality, biogeography, and emergent coexistence patterns in model plankton ecosystems.

## Introduction

Marine phytoplankton is a diverse group of photosynthetic microorganisms that include prokaryotic cyanobacteria, diatoms, and other numerous eukaryotic groups (Falkowski et al. 1998; De Vargas et al. 2015; Dutkiewicz et al. 2020). They are at the base of the ocean food web (Field et al. 1998), play a crucial role in global biogeo-chemical cycles (Eppley and Peterson 1979) and are key contributors to the biological carbon pump (Henson et al. 2012; Guidi et al. 2016). In the sunlit ocean, molecular techniques have characterized a tremendous diversity of phytoplankton groups at the global scale (De Vargas et al. 2015; Sunagawa et al. 2015). Explaining the diversity and the biogeography *i*.*e*. the spatio-temporal repartition of different species across the oceans is a fundamental and long-standing issue in marine ecology.

At the global scale, phytoplankton abundance, community composition, and diversity are spatially hetero-geneous. From genomic data, it was shown that plankton communities exhibit a pronounced biogeographical structure (*β* diversity) that depends on size classes and phylogenetic groups (Sommeria-Klein et al. 2021; Richter et al. 2022; Frémont et al. 2022). For phytoplankton, in large scale ecosystem models, cell size emerges as a dominant organizing trait, with small cells prevalent in nutrient-poor, stratified regions and larger cells dominant in productive environments (Dutkiewicz et al. 2020). Size-dependent physiological constraints therefore appears as the main explainer of phytoplankton biogeographical structuring with small cells having high nutrient affinity but larger cells higher maximum growth rates (Ward et al. 2012). Nevertheless, these trait-based explanations alone do not fully account for the observed levels of local richness (*α* diversity) and persistence.

At the local scale, *α* diversity also exhibits pronounced biogeographical patterns. Across plankton groups, diversity generally decreases from equatorial regions toward the poles, with relative minima in the cores of tropical gyres (Hillebrand 2004; Fuhrman 2008; Yasuhara et al. 2012; Ibarbalz et al. 2019; Busseni et al. 2020; Dutkiewicz et al. 2020). Large-scale environmental patterns, including temperature (regulating metabolic rates), gradients in nutrient supply and light availability, and the influence of seasonal and mesoscale variability, which together shape coexistence, productivity and the range of viable ecological niches across latitudes, have been proposed to explain this gradient (Rohde 1992; Allen et al. 2007; Brown et al. 2004; Hillebrand 2004; Dutkiewicz et al. 2020). Diversity maxima generally occur in biogeographical transition zones rather than in the strongly seasonal or oligotrophic regions.

Despite these well-established large-scale patterns, explaining how high phytoplankton diversity is maintained locally remains a long-standing challenge. At the local scale, cells depend for their growth on a limited set of inorganic nutrients such as nitrogen (ammonium, nitrite, and nitrate), phosphate, iron (Sakka et al. 1999) and silica for diatoms (Flynn and Martin-Jézéquel 2000). When the dynamics of the population and nutrients follow a simple resource–consumer model, the competitive exclusion principle implies that, for N limiting resources, no more than N species can coexist in equilibrium (Armstrong and McGehee 1980). This discrepancy between observed diversity and theoretical competitive exclusion has been called the “paradox of the plankton” (Hutchinson 1961).

Theoretical and field studies have proposed potential mechanisms of community emergence, dynamics and local coexistence between species competing for the same resources (Huisman and Weissing 1999; Thingstad 2000; Follows et al. 2007; Barton et al. 2010; Dunne et al. 2013; Ottesen et al. 2014; Aylward et al. 2015; Dutkiewicz et al. 2020; Muratore et al. 2022). In resource competition models, increasing the complexity of the model by adding new resources can lead to chaotic behavior, where turnover and coexistence can emerge from such dynamics (Huisman and Weissing 1999). Physical stirring can also promote coexistence as mesoscale eddies that trap and isolate populations have been shown to sustain competitors that would otherwise be excluded (Bracco et al. 2000). Trait-based and size-structured ecosystem models have shown that large-scale biogeographical and diversity patterns can emerge from trade-offs among physiological traits interacting with environmental gradients, even under strong resource competition (Follows et al. 2007; Barton et al. 2010; Dutkiewicz et al. 2020). In these models, dispersal by ocean currents and eddies can locally increase species coexistence well above what environmental selection allows, particularly at the boundaries between biogeo-graphical provinces (Clayton et al. 2013; Lévy et al. 2015). Selective top-down control, such as differential grazing and specific viral predation, also strongly affects community dynamics and composition, and can promote microbial coexistence by preferentially suppressing the most competitive and abundant taxa, generating negative frequency-dependent selection (Thingstad 2000; Dunne et al. 2013; Dutkiewicz et al. 2020).

A promising alternative to trait-based Eulerian approaches (treating plankton tracers as homogeneous in fixed grid boxes) is particle-based modeling, in which plankton are represented as individual cells or super-individual particles (each representing a cohort of cells), with each particle carrying a unique set of properties, behaviors, and interactions that evolve through time (Hellweger, Clegg, et al. 2016). By coupling a neutrally evolving genome to each particle in a Lagrangian particle-based model (PBM) embedded within a large-scale circulation, the interaction between ocean currents and neutral genetic drift has been shown to be sufficient to generate large-scale biogeographical patterns, with total genetic diversity theoretically scaling with the number of modeled particles (Hellweger, Van Sebille, et al. 2014). More generally, PBMs resolve the Lagrangian environmental histories experienced by plankton along advective pathways that can differ markedly from Eulerian variability at fixed locations—with important consequences for community growth rates (Zaiss et al. 2021), and temperature-driven selection (Doblin and Sebille 2016; Hellweger, Sebille, et al. 2016). Oceanic transport barriers like eddies have also been shown to generate and maintain spatially distinct phytoplankton patches, associated with different fluid-dynamical niches and water masses histories (d’Ovidio et al. 2010). Finally, previous theoretical work has shown that representing populations as discrete individuals rather than continuous densities can qualitatively alter the predicted outcomes of spatial competition and coexistence (Durrett and Levin 1994). Whether similar effects emerge in plankton ecosystem models remains poorly understood and better understanding how the environmental history of plankton communities along currents affects their biogeography, diversity, and community dynamics is an important question.

In this study, we combined a trait-based formulation (Follows et al. 2007) with a custom PBM framework. Each super-individual particle, representing a large number of cells, is assigned to a species, with multiple particles representing the same species across the spatial domain. Species are characterized by nutrient affinity, maximum growth rate, and a thermal niche, following size-structured allometric relationships (Dutkiewicz et al. 2020). Particles are advected and diffused within an idealized North Atlantic double-gyre circulation and experience forced seasonal Eulerian fields of a single nutrient (nitrate), light, and temperature (Figure 1a). We aimed to compare emergent large-scale biogeographical patterns, local richness and diversity, and environmental variability between the purely Eulerian and PBM approaches (Figure 1b). Our simplified approach was motivated by three main questions. First, we asked whether particle-level representation of biomass, as opposed to Eulerian population averages, could more effectively track local environmental variability and thereby enhance species persistence. Second, we investigated which biogeographical and successional patterns could emerge from these particle-level dynamics. Finally, we compared Lagrangian environmental histories experienced by particles with local Eulerian conditions and further examined whether these histories could be linked to particle lifespan hypothesizing that it could explain differences between the two types of model. We hypothesized that PBM outputs would differ from those of the Eulerian model depending on the mortality threshold and the number of modeled particles. In the PBM, mortality was deterministic and resolved at the level of super-individual particles, whereas in the Eulerian model it was applied at the population level within each grid cell, using the same concentration threshold in both cases (Figure 1c,d).

**Figure 1:**
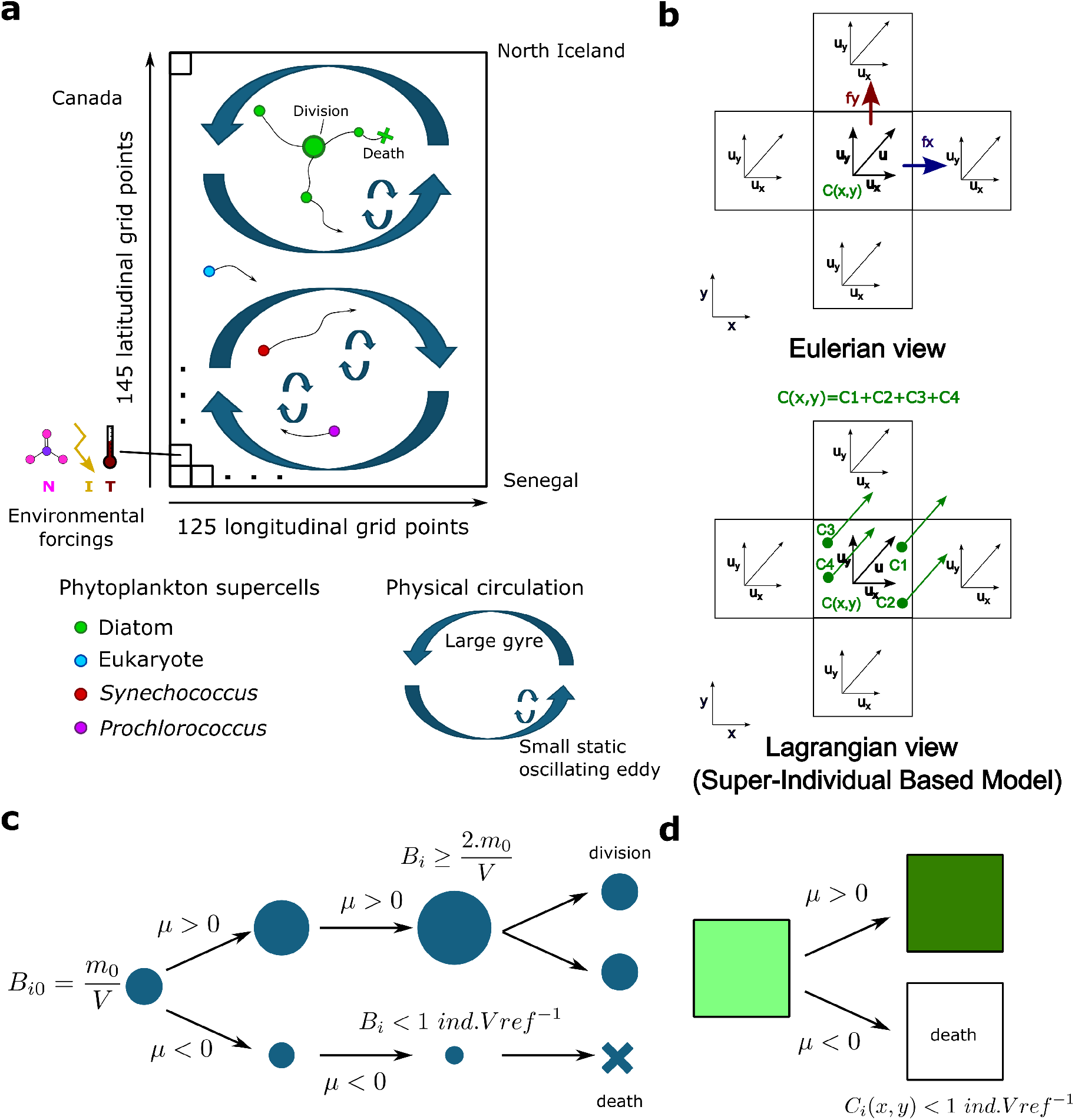
Principles of the ecology and advection of the Particle-Based Model and the Eulerian framework. (**a**) Scheme of the circulation model and Particle-Based Model (PBM) dynamics. The grid is divided in 125 longitudinal points and 145 latitudinal points. All super-individual particles within one grid point compete for the nutrients in this grid point as their total representative concentration is controlled by the nutrient concentration in the grid point. (**b**) Comparison of the Eulerian and Lagrangian framework for the advection of the phytoplankton, (**c**) Schematic of the growth and death processes of super-individual particles in the PBM. The size of the particle represents the biomass concentration it represents in a grid cell (*B*_*i*_), *µ* is the effective growth rate, *m*_0_ is the initial biomass of the particle, *V* is the volume of a grid cell considering a depth of 1 m, and *V ref* is the reference volume for mortality. (**d**) Schematic of the growth and death processes of super-individual particles in the Eulerian framework. The darkness of the color represents the biomass concentration (*C*_*i*_(*x, y*)) of a given species in a grid point.

## Materials and methods

### Model concept

We built a PBM in which each particle is advected (see Particles transport) in a dynamical two-dimensional double gyre flow (see Idealized North Atlantic double gyre flow), and grows/shrinks in biomass following deterministic dynamical equations (see Particle growth model) and dynamical one year long geochemical forcings (see Growth equilibrium: constraining phytoplankton biomass and Physical and geochemical forcing). The same forcings are repeated each year of simulations (10 year long simulations, see Simulations). Each particle is assigned to a phytoplankton species characterized by its taxonomy (*Prochlorococcus, Synechococcus*, non-diatom Eukaryote, or Diatom), size and temperature niche (Figure S1). Each particle is a super-individual: it represents a large number of cells from its assigned species and dies and duplicates following fixed conditions (see Particle biomass concentration, duplication and death). At the initiation of each simulation, an initial number of particles is randomly assigned to a given species from a fixed set of 400 species in total (all 400 species are always represented) and spread evenly across the spatial domain. Finally, we built an equivalent Eulerian model in which all 400 species are advected and grow/shrink in biomass following the same dynamical equations and biogeochemical forcings as the PBM (see Eulerian model). Both models represent the same physical advection–diffusion processes, but differ in how they are numerically represented. In the PBM, transport is represented by the movement of individual particles, including a Brownian component corresponding to the diffusion term in the Eulerian equations, whereas in the Eulerian model, transport and diffusion are resolved on a spatial grid. Mortality is also applied differently in the two models: at the particle level in the PBM and at the grid-point level in the Eulerian model.

### Particle-based ecological model

#### Particle growth model

For each particle, we use a Monod growth equation generalized to n species that resembles the Generalized Lotka-Volterra (GLV) model (Case 1999; Pontarp and Petchey 2018). All interaction terms are equal in the GLV formulated in Equation 2.1 of Pontarp and Petchey 2018, representing equal and complete competition for nutrients. Concretely, this is translated into a carrying capacity term dependent on the nutrient field to constrain total phytoplankton biomass (see Growth equilibrium: constraining phytoplankton biomass). Finally, a grazing term of a single generalist zooplankton type (i.e. it predates all phytoplankton species non specifically) is included and zooplankton biomass dynamics is modeled using the same particles as for the phytoplankton. Thus, the equation of phytoplankton biomass dynamics for a particle of biomass concentration *B*_*i*_ in control grid point *K* is the following:

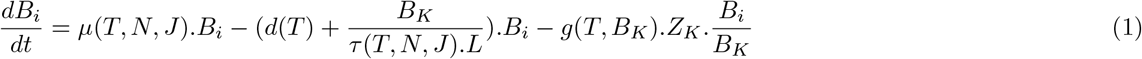

*B*_*i*_ is the biomass concentration represented by particle *i* expressed in *mmolN*.*m*^*−*3^. *µ*(*T, N, J*) is the effective growth rate of the particle *i* in *d*^*−*1^ that depends on temperature *T*, light *J*, and nutrient *N* limitation terms (unitless, described in Model of phytoplankton growth and full equations in Supplementary information). *d*(*T*) is the linear death rate of phytoplankton in *d*^*−*1^ dependent on temperature. *g*(*T, B*_*K*_) is the effective grazing rate in *d*^*−*1^ that depends on temperature and a Holling type III function for grazing (see below and extended formulation in Supplementary information). *B*_*K*_ = ∑_*Kj*_ *B*_*Kj*_ and *Z*_*K*_ = ∑_*Kj*_ *Z*_*Kj*_ are, respectively, the total phytoplankton and zooplankton biomass concentration in a grid cell *K. L* is a constant in *mmolN*.*m*^*−*3^. The equation for the zooplankton biomass *Z*_*i*_ (transported in the same particle as the phytoplankton) is:

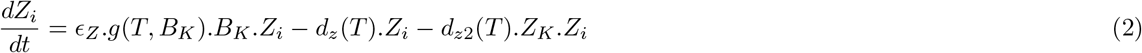

where *d*_*Z*_(*T*) in *d*^*−*1^ and *d*_*z*2_(*T*) in (*mmolN*.*m*^*−*3^)^*−*1^.*d*^*−*1^ are, respectively, the linear and quadratic mortality term associated to the zooplankton that depend on temperature and *ϵ*_*Z*_ is the grazing efficiency (unitless). The quadratic mortality term represents higher order losses due to predation of the zooplankton by higher trophic levels (not explicitly represented). To conserve zooplankton biomass, when a super-individual particle dies (see Particle biomass concentration, duplication and death), the zooplankton biomass that was in the super-individual particle is redistributed homogeneously to nearby super-individual particles from the same control grid point. When no other super-individual particle is present at this grid point, the zooplankton biomass disappears. These equations do not account for the transport of biomass that is done at the level of the super-individual particle in the PBM (see Particles transport).

#### Particle biomass concentration, duplication and death

The representative initial biomass concentration in a control grid cell *K* of a particle *i* is:

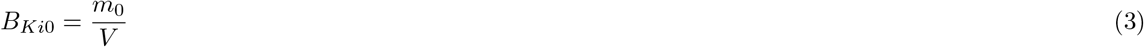

where *m*_0_ is a constant biomass and *V* is the volume of a grid cell *K* in *m*^3^ considering a depth of 1 meter (not the mixed layer depth, see additional remarks on particle biomass representation in the Supplementary information). The choice of *m*_0_ ultimately set the number of particles in a simulation, as it sets the threshold at which a particle duplicates. Particles duplicated when they reached twice their initial biomass concentration following the duplication condition:

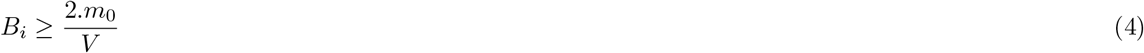

A particle died and was erased from the set of particles upon reaching a critical concentration following the death condition:

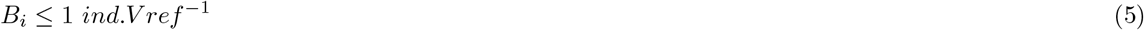

where *V ref* is a chosen volume of reference (1, 10 or 100). Concentration is converted to *ind*.*m*^3^ using cell quotas (see Supplementary information).

### Eulerian model

We developed an Eulerian version of the PBM model where the concentration of each phytoplankton species is followed at each grid point of the 145×125 grid domain.

#### Species growth model

From equations 1 and 2, the dynamical equations of total biomass, *B*_*Ks*_, of a species *s* of phytoplankton and the zooplankton biomass *Z*_*K*_ in a control grid cell *K* are, by summing over the particles’ equations in this control grid cell (of a given species for the phytoplankton and over all particles for the zooplankton):

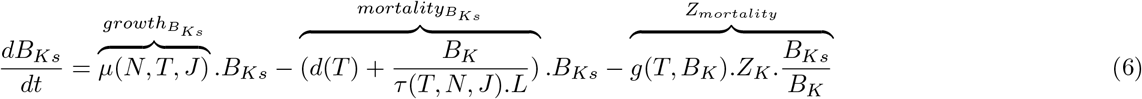

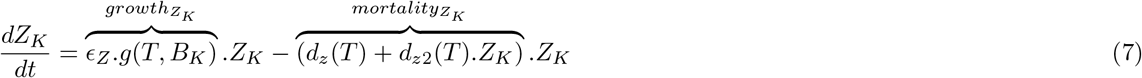

See Supplementary information for the extended version of these equations.

#### Eulerian advection

To advect phytoplankton species in the Eulerian model, we add the diffusion and advection terms to the species growth model (equations 6 and 7):

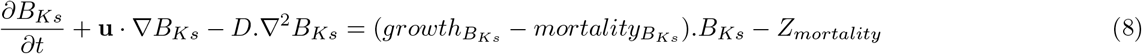

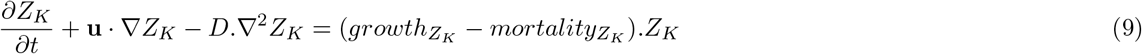

where **u** is the flow field, ∇ is the nabla operator, · is the dot product and *D* is the diffusion coefficient.

#### Eulerian mortality

In the case of the Eulerian model, a death threshold was applied at each grid point: if the concentration of a species is below 1 *ind*.*V ref* ^*−*1^, its concentration is set to 0. This criterion is fundamentally different from the one of the PBM version as it is applied at the level of the grid point and not the particle. Such thresholds are needed as continuous-density models can otherwise sustain populations at unrealistically low concentrations, the so-called “atto-fox” problem (Mollison 1991).

### Chlorophyll model

To compare our model outputs with large scale satellite product data (Sauzéde et al. 2025), we modeled chlorophyll dynamics following Geider et al. (1997) and the Redfield ratio (Redfield 1934). Briefly, the chlorophyll model allows the phytoplankton to have a dynamical chlorophyll-to-carbon ratio reflecting its physiological and photoacclimation state (Geider et al. 1997) (see equations in Supplementary information).

### Model of phytoplankton growth

#### Growth equilibrium: constraining phytoplankton biomass

To constrain maximum biomass reached by the phytoplankton in our model, we used the nutrient field (see Temperature and nitrate fields). At equilibrium, without the zooplankton, we set *B*_*K*_ = *N*, where *N* is the nutrient concentration in the control grid cell *K*. This assumes that all available nutrients could be consumed. Equating equation 1 to 0 without the zooplankton mortality term yields:

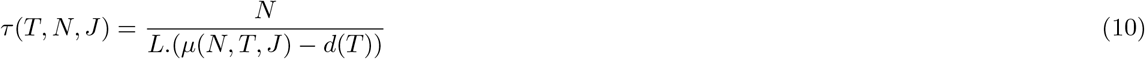

We fixed *L* to 1 *mmolN*.*m*^*−*3^ (either *τ* or *L* could be fixed). Equation 10 is used to limit the total concentration of phytoplankton biomass at equilibrium as the concentration of nutrients in the control grid cell when zooplankton is absent. Practically, there are several species in a grid cell with different effective growth rates, so we used the maximum effective growth rate of all particles in a grid point, *i*.*e*. the one of the fastest growing species to determine *τ*. Therefore, *τ* is unique in a grid cell so the competitive exclusion principle applies within a grid cell: without transport and seasonality forcings, only the fastest growing species survives at equilibrium.

#### Maximum growth rate: species definition

Each particle is assigned to a taxon and a size from an initial 100 randomly drawn sizes for each taxa (totaling 400 species): *Prochlorococcus, Synechococcus*, non-diatom Eukaryote or Diatom. The maximum growth rate of the given species then follows an empiric allometric relationship with cell volume following Dutkiewicz et al. 2020: 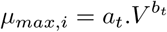, where *a*_*t*_ and *b*_*t*_ depend on the phytoplankton type (Table S2, Figure S1a,b).

#### Temperature dependency and niche

For the temperature dependency *ψ*(*T*) (growth rate dependency including a thermal niche term) and *η*(*T*) (mortality and grazing dependency to temperature), we followed Dutkiewicz et al. 2020 for the niche and the Eppley curve for temperature dependency (Eppley 1972) (see equations in Supplementary information).

#### Light dependency

For light dependency *γ*(*J*), we followed Evans and Parslow 1985 and Stec 2016. Briefly, light limitation depends on the average of a light limitation function between the surface and the mixed layer accounting for irradiation, day length, light attenuation with depth, and self shading effect of the phytoplankton (see equations in Supplementary information).

#### Nutrient dependency

For nutrient dependency *β*(*N*), we used the limitation term of the Monod equation of growth (Monod 1949) and allometric relationships based on cell volume for the half saturation constant (Dutkiewicz et al. 2020) (see equations in Supplementary information). The cell volume also determines the cell quota (see Supplementary information) necessary to constrain the concentration-based mortality we implemented.

#### Grazing limitation

For grazing limitation, we chose a Holling type III limiting function (Holling 1959) commonly used in biogeo-chemical models (Zahariev et al. 2008; Law et al. 2017; Dutkiewicz et al. 2020) (see equation in Supplementary information)

### Idealized North Atlantic double gyre flow

#### Main flow

We build an idealized double gyre flow consisting of two principal gyres (north and south) with the dimensions of the real North Atlantic double gyre and static eddies. The well known and extensively studied double gyre equation (Shadden et al. 2005; Froyland and Padberg 2009; Froyland and Padberg-Gehle 2015; Banisch and Koltai 2017; Reijnders et al. 2021) is used to mimic the double gyre. The speed potential is given by:

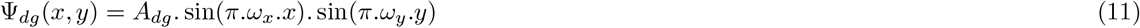

Meridional and zonal flow speeds are derived in Supplementary information. Longitudinally, the dimension of the gyre correspond approximately to the width of the North Atlantic Ocean between the french city of Bordeaux to the Canadian province of Nova Scotia (5000 *km*). Latitudinally, it corresponds approximately to the distance of a transect between the Senegal upwelling east of Dakar and the north of Iceland (54° of latitude) (5800 *km*). The maximum speed of the main flow is set to 0.15 *m*.*s*^*−*1^.

#### Mesoscale static eddies

In addition to the main double gyre, we approximately modeled mesoscale eddies using the same equations with different scale parameters. Eddies were static but oscillated in time (resulting in a periodic change in the direction of their flow). Equations are given in Supplementary information. Eddies have a spatial scale of 250 *km* and oscillate over a 2 month period. The maximum speed of the eddies field was set to 0.15 *m*.*s*^*−*1^. The addition of the main and eddies field allowed to reach a maximum speed of approximately 0.3 *m*.*s*^*−*1^ which is what is observed in the real oceans except in the Gulf Stream where higher speeds are reported (Marzocchi et al. 2015).

#### Diffusion

In addition to the main velocity field, we model small scale turbulence by adding a 2 dimensional Brownian motion to the particles. Brownian motion is modeled as follows: at each time step the particle is transported in a random direction of the 2 dimensional grid according to the diffusion equation with a constant diffusion coefficient *D* in *m*^2^.*s*^*−*1^ (see equations in Supplementary information).

#### Particle speed

For a particle *i*, its total speed is the sum of the main flow, eddies and Brownian motion (see equations in Supplementary information). Details of particle transport implementation is provided in Supplementary information.

### Physical and geochemical forcings

We built space- and time-dependent physical (mixed layer depth, light and temperature) and geochemical forcings (a single nutrient, nitrate) to constrain the growth of the phytoplankton community. Each field has a 1 day time resolution over the 125×145 spatial grid. The phytoplankton dynamic has no feedback effects on the forcings.

#### Light field

The irradiation *J* received by a phytoplankton at a given day of the year and a given latitude depends on two main parameters: the solar irradiation, that depends on the day of the year, and the cosine effect, that depends on latitude and day of the year, taking into account the inclination of solar rays with the Earth. Details of the equation of irradiation (*J*) are given in the Supplementary information. In addition, note that the light limitation growth term of the phytoplankton depends on the Mixed layer depth (MLD), self shading and day length (see Light dependency).

#### Mixed Layer Depth field

We built a daily MLD climatology over our spatial domain (125×145 spatial grid) using a regular 2°C×2°C grid temperature-based global scale MLD climatology from the IFREMER (De Boyer Montégut et al. 2004). The dynamical Mixed Layer field was used to compute the temperature field, the nutrient field and the light dependency for growth. Details of the MLD field construction are given in the Supplementary information.

#### Temperature and nitrate fields

- Main fields

We built a daily temperature and nitrate field over our spatial grid (125×145 grid) using the 1°x1° monthly data from the World Ocean Atlas 2018 (Locarnini et al. 2018; Garcia et al. 2018) as forcing fields. In the dynamical ecological model, both temperature and nitrate were independent from the flow field and any feedback from the ecology. Additional details of field construction are given in the Supplementary information.

- Eddies nutrient field

In addition to the main nutrient field, we add a nutrient field proportional to the natural logarithm (*ln*) of the eddies field. Because the stream function changes sign, it is first linearly rescaled to the range between the natural logarithms of imposed minimum and maximum nutrient concentrations. We then take the exponential of the rescaled field to obtain positive nutrient concentrations. In the northern gyre, nutrient eddies field was added to the main nutrient field while it was subtracted in the southern gyre following the paradigm that cyclonic and anticyclonic eddies respectively increase and decrease the accumulation of phytoplankton biomass (McGillicuddy 2016). Additional details of field construction are given in the Supplementary information.

### Simulations

Simulations were initialized with various values of *m*_0_ for the PBM and with three mortality thresholds defined by the reference volume (*V*_ref_ = 1, 10, and 100 L). The choice of *m*_0_ determines the initial number of particles. For both the PBM and Eulerian models, identical initial total biomass concentrations were used. Unlike the Eulerian model, in which all species are initialized across the spatial grid, the finite number of particles in the PBM imposes that many species are initially absent from some grid cells, particularly at low particle numbers. The initial species-by-cell distributions are therefore not equivalent between the two models.

Even though the systems did not reach a strict equilibrium as some species were on the way to exclusion in both the PBM and Eulerian simulations (Fig. 5), all comparisons were conducted over the final tenth year of each simulation to ensure consistency across simulations. Model implementation details are provided in the Supplementary information.

### Lagrangian tracking

For the simulations with the highest particle numbers at each reference volume, we tracked a total of 2.4 million particle trajectories. Tracking began with 200,000 particles on January 1st of the ninth simulation year, with an additional 200,000 unique particles (ie without redundancy) tracked on the first day of each subsequent month through December, ensuring seasonal representativeness. Along trajectories, we recorded each day species identity, latitude, longitude, temperature, nutrient, light, biomass and particle identity (to track particle divisions). All particles were tracked until their death or the end of the simulation. When particle divided, only the newborn particle was tracked (enabling division tracking without following new particles).

### Satellite-derived chlorophyll product

We used reconstructed global three-dimensional chlorophyll products derived from Copernicus satellite observations spanning 1998–2023 (Sauzéde et al. 2025). These products rely on a machine-learning-based reconstruction of the vertical particulate backscattering coefficient (Sauzède et al. 2016), combined with vertically resolved models of downwelling irradiance and photosynthetically available radiation at the global scale (Renosh et al. 2023). When available, chlorophyll concentrations were averaged over our spatial domain over the yearly climatology accounting for MLD temporal variations.

## Results

### Distinct seasonal biomass dynamics emerge in the particle-based model

We first compared the biomass dynamics emerging from the particle-based model (PBM) and the Eulerian model of our phytoplankton community, without species distinction. The two approaches differed fundamentally in both transport representation and mortality formulation (Fig. 1b,c; see Materials and Methods).

Focusing on the dynamics during the final year of the simulations in the northern gyre, the average total phytoplankton concentration tended to exhibit higher peaks in the PBM across all mortality thresholds, compared to the Eulerian model (Figure 2a,b, Figure S2a-f for all reference volumes). In addition, minimum concentrations were lower in the PBM, consistent with a larger seasonal amplitude. The lower winter biomass in the PBM might partly result from the discrete representation: individual particles might more easily fall below the mortality threshold and be removed, even when the total species biomass in the local grid point (represented by other particle of the same species) remains above that threshold. In the southern gyre, total phytoplankton concentration in the PBM appeared to be less stable over time compared to the Eulerian model but was overall sustained at higher levels (Figure 2c,d, Figure S2g-i for all reference volumes). These results are consistent with the PBM capturing aspects of phytoplankton dynamics that are less well represented in the Eulerian framework, such as more localized bloom events in productive regions and more sustained phytoplankton biomass in nutrient-poor regions. Yearly averaged concentrations from the Eulerian and PBM simulations were nevertheless strongly correlated and correlation increased with reference volume (r = 0.93, 0.96, and 0.96 for *V*_*ref*_ = 1, 10, and 100 *L*, respectively; Figure 2e-g and S3,S4). The transient spring-summer peak bloom within the northern gyre was clearly visible in both the PBM and Eulerian animations with a clearer south west to north east increase in the PBM (Videos 1 and Videos 2 for zooplankton). Finally, the PBM model sustained higher biomass in the Senegal equatorial upwelling region while the autumn bloom was not clearly maintained.

**Figure 2:**
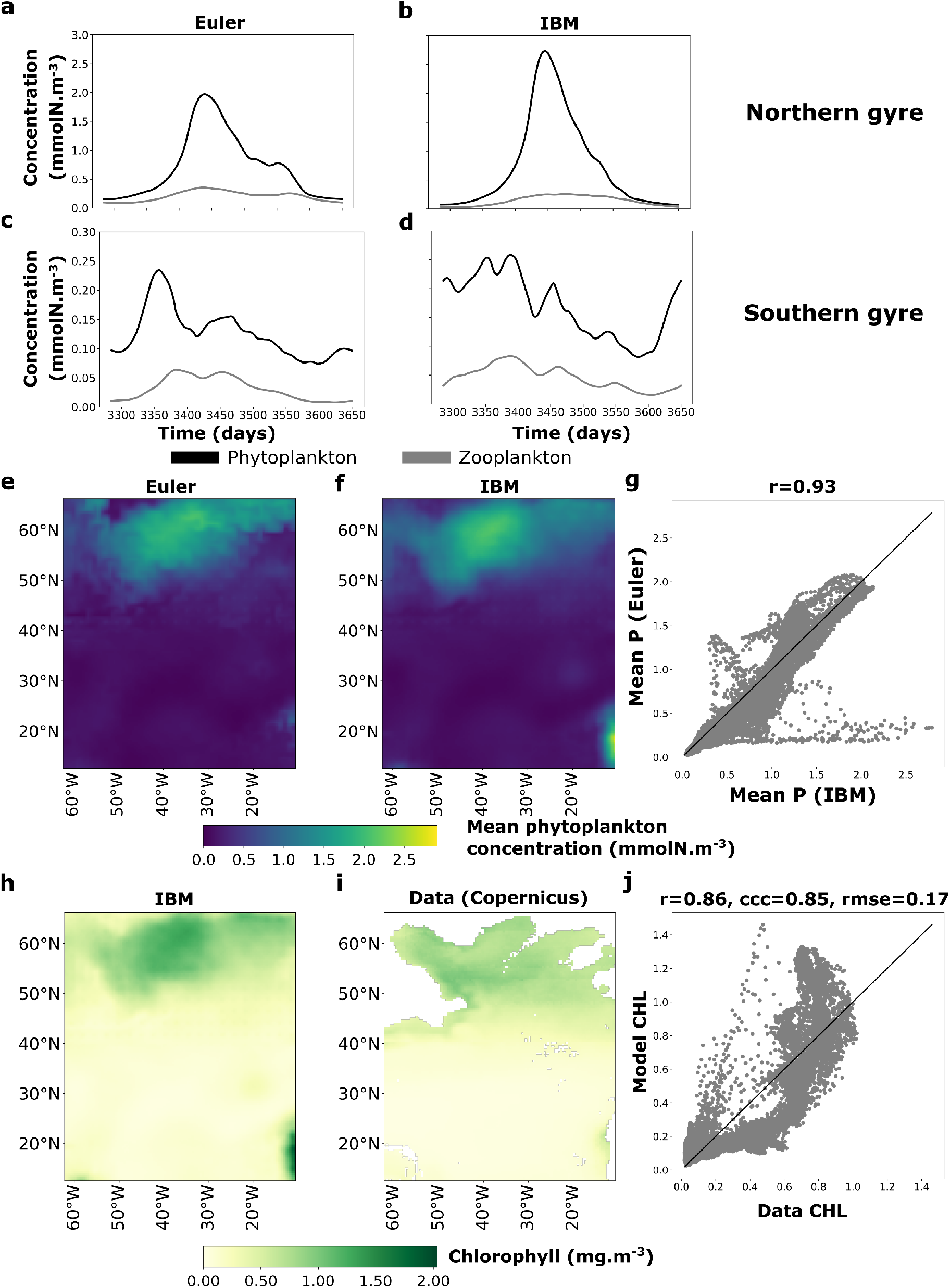
Comparison of biomass dynamics between the PBM and Eulerian model. (**a**–**d**) Time series of spatially averaged phytoplankton and zooplankton concentrations during the final year of simulation in the (**a, b**) northern and (**c, d**) southern gyres, for the Eulerian model (**a, c**) and PBM (**b, d**). (**e, f**) Yearly mean phytoplankton biomass concentration for the Eulerian model (**e**) and PBM (**f**), and (**g**) their spatial Pearson correlation. (**h, i**) Yearly mean chlorophyll concentration for the PBM (**h**) and Copernicus observations (**i**), and (**j**) their spatial Pearson correlation. All panels show the model with a reference volume of 1 L (see Figures S2–S5 for other reference volumes).

To test the robustness of our model against observable ecosystem properties, we modeled chlorophyll dynamics in both models using a photoacclimation formulation (Geider et al. 1997) (see Materials and Methods). We compared simulated chlorophyll fields with satellite-derived chlorophyll products (Sauzéde et al. 2025) (see Materials and Methods). We found strong correlations between modeled and observed chlorophyll (*r* ~ 0.85, ccc –Concordance Correlation Coefficient– ~ 0.85 testing for the y=x relationship, and rmse=0.17 *mgCHL*.*m*^*−*3^ for *V*_*ref*_ = 1 *L* in the PBM), relatively insensitive to the choice of the mortality threshold reference volume (Figure 2h-j, Figure S5 for the Eulerian model and other mortality thresholds). At the basin scale, the PBM tended to overestimate chlorophyll in the most productive northern gyre region while the Eulerian model overestimation was lower. In the southern gyre and intermediate more productive regions, both model underestimated chlorophyll although more strongly for the Eulerian model. Overall, these results demonstrate that the model reproduces large-scale chlorophyll patterns, supporting our approach of constraining nutrient concentrations using observational fields while indicating that the ecological dynamics are consistent with the observed large-scale biomass distribution.

### The particle-based model modulates overall and local richness and diversity

We then compared emerging richness and diversity patterns between the Eulerian and PBM frameworks. Richness was defined as the number of coexisting species at each location, while diversity was quantified using the Shannon index (Shannon 1948; Margalef 1958).

Spatial patterns of richness and diversity were more consistent across PBM simulations than across Eulerian simulations, as indicated by higher pairwise correlations (Figure 3g,h). We examined total richness and mean local richness and diversity across the domain and in each gyre, varying PBM particle numbers by changing the representative biomass *m*_0_ (Figure 4, Figures S7 and S8). At low particle numbers (~ 10^5^), total richness was high, despite low mean and maximum local richness. This may reflect reduced competition among resident dominant species combined with limited dispersal for invading species (see Lévy et al. 2015 for the role of dispersal in local diversity). As mean particle numbers increased toward 10^7^, total PBM richness declined, converging towards Eulerian values, but remained higher than in the Eulerian model, particularly at low reference volumes (*V*_*ref*_), with increases of +5 (+55%), +1 (+8%), and +1 (+7%) species for *V*_*ref*_ = 1, 10, and 100 L, respectively (Figure 4a). Given that each configuration represents a single realization, these small differences may fall within run-to-run variability and should be interpreted cautiously for large *V*_*ref*_. Beyond a certain number of particle modeled, mean local richness increased approximately linearly with particle number, exceeding Eulerian values only at *V*_*ref*_ = 1 *L*. Diversity followed a similar trend, although local values nearly converged between models (Figure S7a). Differences between gyres were pronounced. In the southern gyre, richness and diversity increased with particle number, driving much of the overall increase (Figure 4b,d; Figure S7a,d). In the northern gyre, both metrics were generally higher in the PBM than in the Eulerian model, except at high *V*_*ref*_ (Figure 4e; Figure S7e). In the southern gyre, PBM local richness difference was higher only at *V*_*ref*_ = 1 *L*, while at higher reference volumes it was lower than in the Eulerian model; diversity was lower overall in the PBM (Wilcoxon tests, *p <* 0.001; Figure 4f; Figure S7f). Animations of final-year richness and diversity are available in Videos 3 and 4.

**Figure 3:**
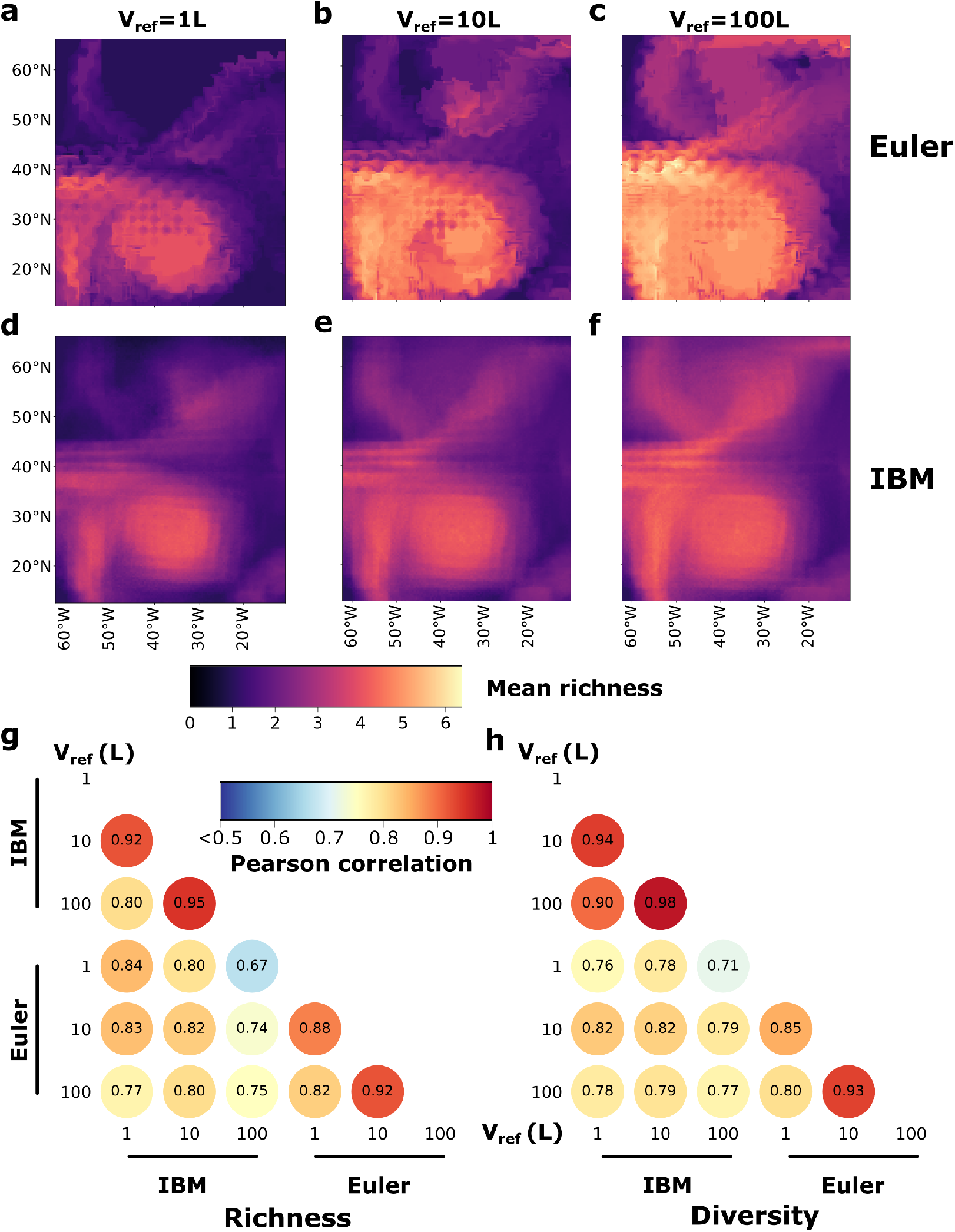
Average richness and diversity of the Eulerian and the PBM model for different mortality-threshold reference volumes. (**a**-**f**) Average richness defined as the average number of species present at each grid point over the last year of simulation for three different mortality-threshold reference volume for (**a**-**c**) the Eulerian model and (**d**-**f**) the PBM model. (**g, h**) Pairwise correlograms of mean richness (**g**) and Shannon diversity (**h**) across models and reference volumes (see Figure S6 for Shannon diversity heatmaps).

**Figure 4:**
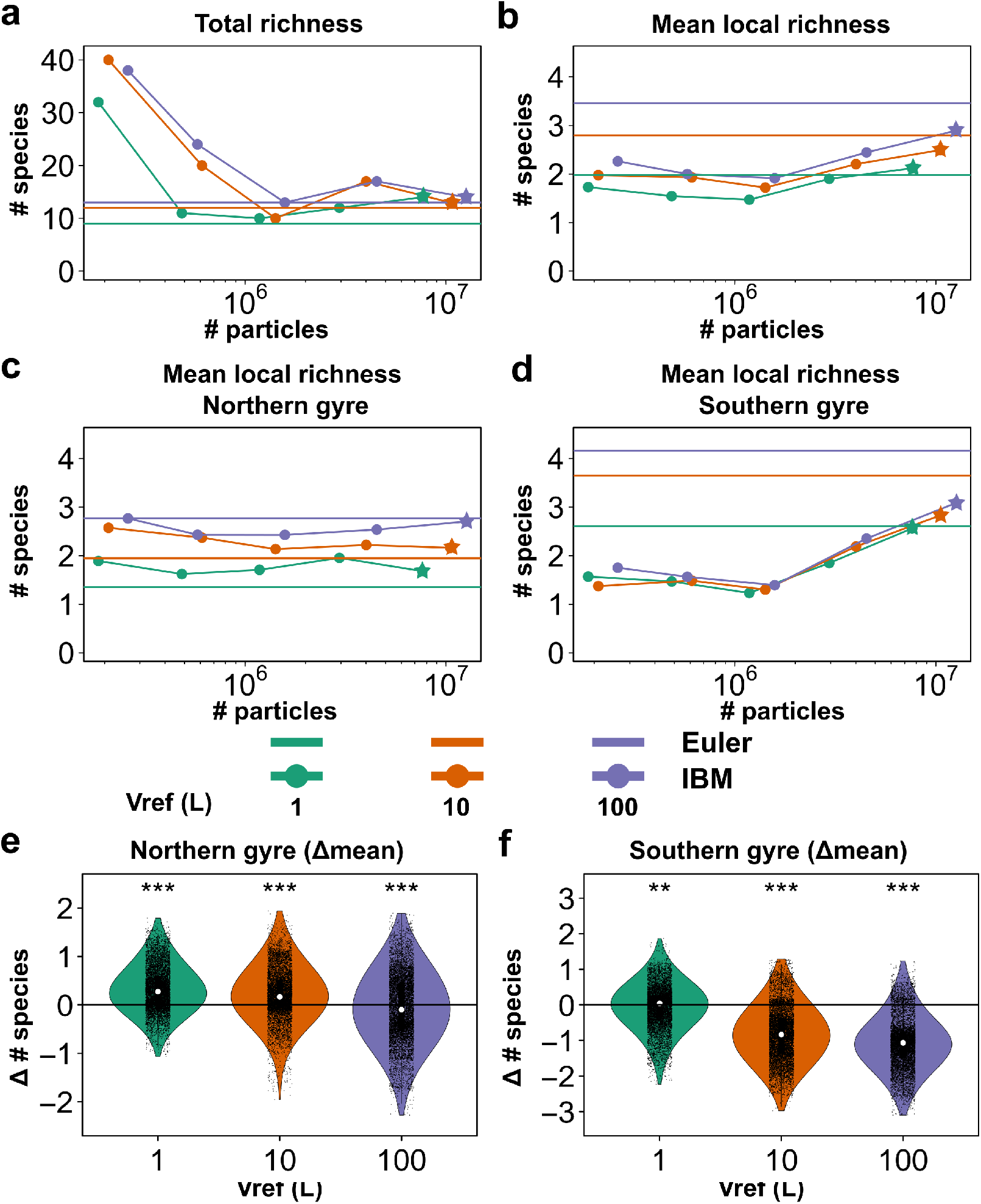
Comparison of emerging richness statistics between the Eulerian model and the PBM. (**a**) Total and time-averaged richness: (**b**) domain mean, (**c**) northern gyre mean, and (**d**) southern gyre mean, for the Eulerian and PBM models as a function of the average number of PBM particles and the mortality-threshold reference volume. Eulerian model data is plotted as horizontal lines for the different mortality-threshold reference volume. PBM simulations presented in Figure 2 and 3 and panels (**e**) and (**f**) are highlighted with stars for the different mortality-threshold reference volumes. (**e**) Violin plots of the difference in average number of present species at each grid point between the Eulerian and PBM model for different mortality-threshold reference volume in the northern gyre and (**f**) the southern gyre. Significantly different median delta mean are indicated by stars (Wilcoxon test, *p <* 0.001).

**Figure 5:**
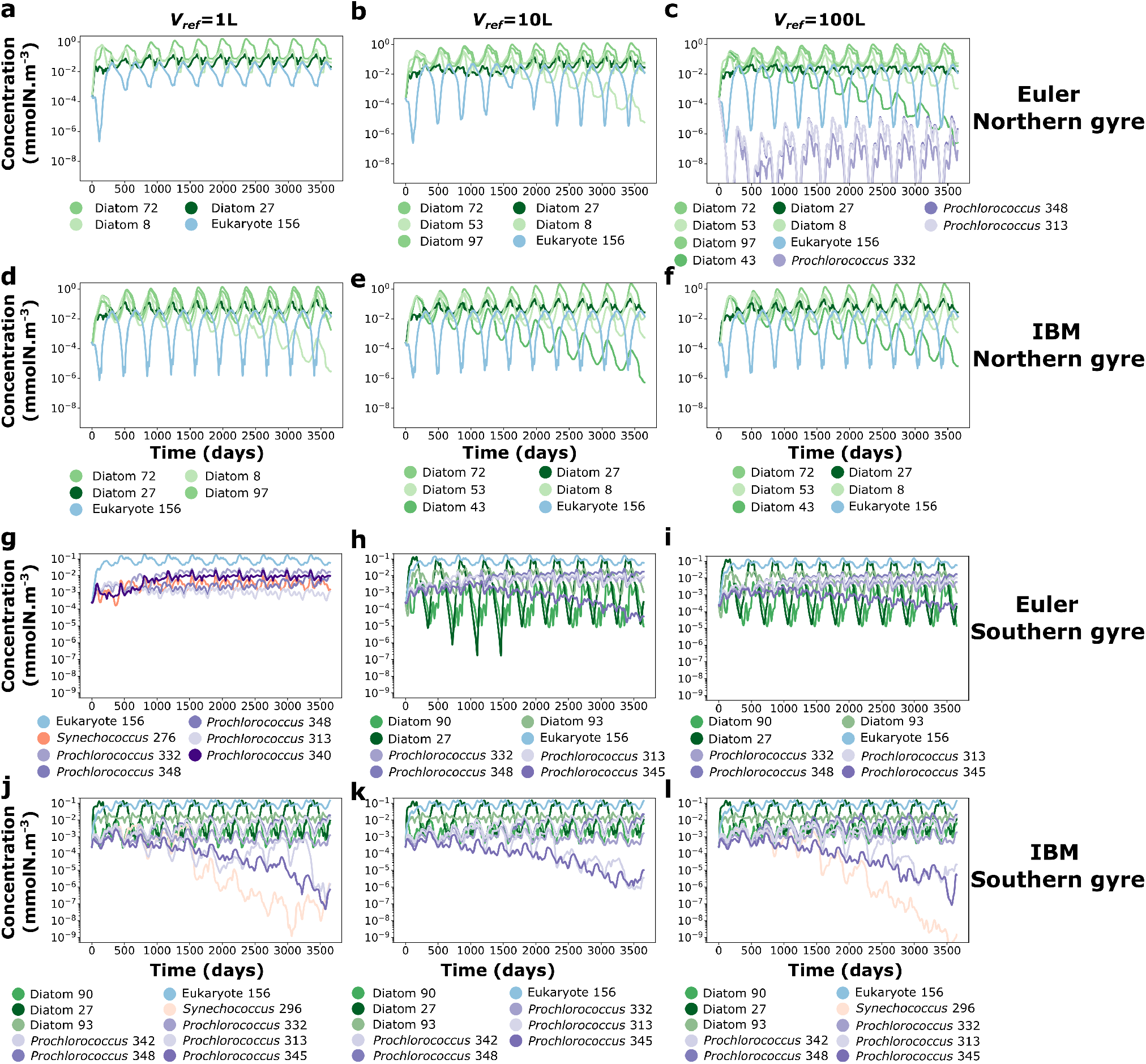
Ten years time series of average concentration of surviving species in the northern and southern gyres for the two different models and different mortality-threshold reference volumes. Eulerian model in the northern gyre for (**a**) *V*_*ref*_ = 1*L*, (**b**) *V*_*ref*_ = 10*L* and (**c**) *V*_*ref*_ = 100*L*. PBM model in the northern gyre for (**d**) for *V*_*ref*_ = 1*L*, (**e**) *V*_*ref*_ = 10*L* and (**f**) *V*_*ref*_ = 100*L*. Eulerian model in the southern gyre for (**g**) *V*_*ref*_ = 1*L*, (**h**) *V*_*ref*_ = 10*L* and (**i**) *V*_*ref*_ = 100*L*. PBM model in the southern gyre for (**j**) *V*_*ref*_ = 1*L*, (**k**) for *V*_*ref*_ = 10*L* and (**l**) *V*_*ref*_ = 100*L*. Diatoms are in shades of green, eukaryotes (non-diatom) in shades of blue, *Synechococcus* in shades of red and *Prochlorococcus* in shades of purple.

Overall, the PBM produced higher regional (*γ*) diversity than the Eulerian model, while differences in local richness and diversity depended on particle number, gyre, and mortality threshold. Local diversity was generally enhanced in the productive northern gyre, dependent on the mortality threshold, whereas differences in the less productive southern gyre were more sensitive to the particle number.

### The particle-based model captures more robust biogeographical dynamics and structure

We then compared biomass dynamics and spatial organization over 10 years of simulation by decomposing total biomass among surviving species across mortality thresholds, defined by reference volume (Materials and Methods), separately for the northern and southern North Atlantic gyres (Figure 5, Figure S9 for a zoom on the last year of simulation). In the Eulerian model, species assemblages differed across mortality thresholds especially in the northern gyre (Figure 5a–c and Figure 5g–i), whereas they were better maintained across thresholds in the PBM (Figure 5d–f and Figure 5j–l). In addition, the dynamics of individual species were more consistent across mortality thresholds in the PBM than in the Eulerian model. Overall, the PBM showed greater robustness, with higher Bray–Curtis similarity among average biomass time series in the last year (Figure S10). Emergent dynamics and biogeographies were robust to initial conditions in both models (Figures S11,12) except for the Eulerian model in the southern gyre for initial biomass conditions (Figures S12c). Most species exclusions occurred during the first two years of the simulation, although we observed a generally longer persistence of a few species in the PBM.

We then visualized patterns of yearly averaged dominant species and biomass concentrations (Figure 6). Consistent with the more robust dynamics observed in the PBM, the biogeographical structure of dominant species was more robust across mortality thresholds in the PBM than in the Eulerian model (Figure 6). For example, in the western part of the southern gyre, dominant species differed markedly across mortality thresholds in the Eulerian model, whereas they were largely conserved in the PBM, with only minor differences for *V*_ref_ = 1 L. Animated GIFs of the final simulation year for dominant species are available in Videos 5. In the Eulerian model, species temporally disappeared in eddy centers where nutrients reached near-zero concentrations. Notably, this phenomenon was absent in the PBM. The greater robustness of PBM biogeography is reflected in higher Adjusted Rand Indices (ARI) for dominant species and higher Bray–Curtis similarities for full biogeographies, which account for the spatial distributions of all species, across PBM simulations (Figure 6g,h). In addition, the PBM consistently supported a greater number of species overall across mortality thresholds, with more species persisting in the southern gyre than in the Eulerian model (Figure S13).

**Figure 6:**
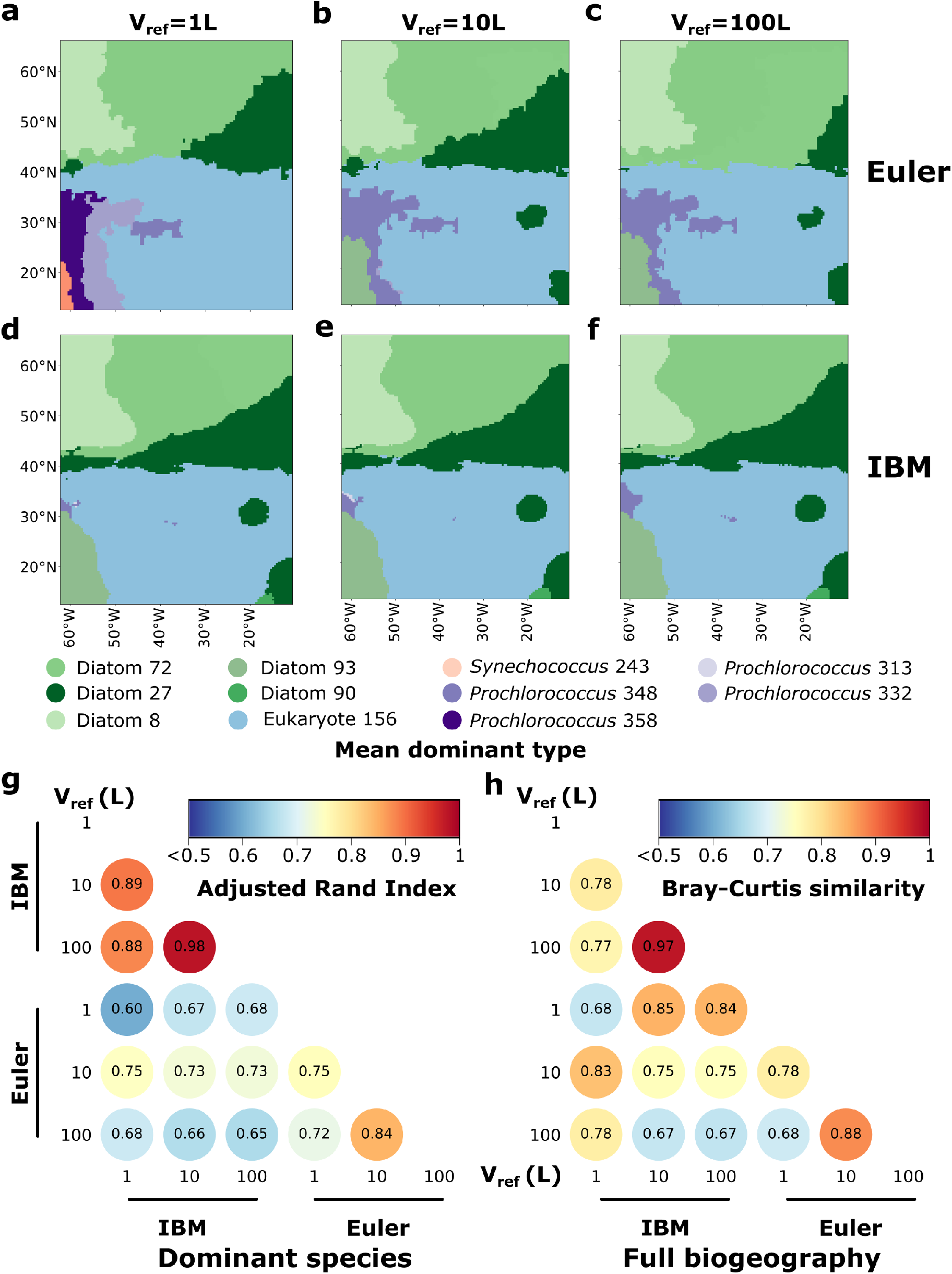
Comparison of the average biogeographies of the Eulerian and the PBM model for different mortality-threshold reference volumes. (**a**-**f**) Average most abundant species on average (last year of simulation) at each grid point for three different mortality-threshold reference volume for (**a**-**c**) the Eulerian model and (**d**-**f**) the PBM model. (**g**) Pairwise Adjusted Rand Index (ARI) between average dominant species of the last year of simulation. (**h**) Pairwise Bray-Curtis similarity indices between average full biogeographies (all species average biomass at each grid point taken into account) of the last year of simulation.

Together, these results indicate that the PBM yields more robust species-level dynamics and biogeographical patterns than the Eulerian model, largely independent of the chosen mortality threshold. This robustness was reflected in the more consistent persistence, spatial distribution, and dynamics of species across mortality thresholds.

### Lagrangian environmental histories exhibit stronger variability in the northern gyre

We then investigated how the Lagrangian PBM can inform our understanding of the role of environmental histories in population dynamics. Using 90-day trajectory segments (called paths) spanning the annual cycle, we compared the mean environmental conditions experienced by Lagrangian particles with the corresponding Eulerian histories at their arrival locations (Figure 7). For temperature, Lagrangian histories were significantly colder than fixed-location Eulerian histories in the southwestern part of the northern gyre, but warmer in its northeastern part. These patterns were generally weaker and reversed in the southern gyre, consistent with the opposite circulation directions of the two gyres. Particles experiencing colder temperature histories generally encountered higher nitrate concentrations but lower light exposure (and reversely). Nitrate-history differences were significantly weaker in the southern gyre, except in the Senegal upwelling region. The standard deviations of the differences between Lagrangian and Eulerian environmental histories (Figure 7d–f) further revealed local variability in these offsets. Temperature differences were most variable along the western portions of the fronts of both gyres, whereas nitrate differences showed greater variability in the northern gyre. Light exposure exhibited more widespread variability across the domain. We repeated the latter analysis using all particle trajectories, weighting the contributions by trajectory duration (Figure S14). Spatial patterns were qualitatively similar, but the anomalies were generally stronger, and extended farther along the direction of circulation in both gyres and across all three environmental variables. These results highlight how longer particle trajectories integrate environmental conditions over larger distances and emphasize the influence of circulation on cumulative environmental exposure. The highest variability in nutrient histories occurred in the western-to-northern region of the domain, broadly corresponding to the locations of the Gulf Stream and North Atlantic Current, highlighting the strong variability encountered along these trajectories, although these currents are not explicitly resolved by our idealized circulation (Figure S14e). Finally, we computed median differences between the encountered environment and environmental species optima for both 90 days paths and full trajectories (Figure S15). Temperature differences revealed negative anomalies along fronts in the northern gyre, predominantly positive anomalies in the southern gyre, and a positive hotspot along the eastern part of the inter-gyre jet (corresponding to the location of North Atlantic Current). Nitrate differences were mostly positive, with negative hotspots in the southern gyre. Overall, these results highlight that plankton populations experience spatially variable environmental selection pressures, sometimes far from their optima, which differ between the two gyres and across environmental variables.

**Figure 7:**
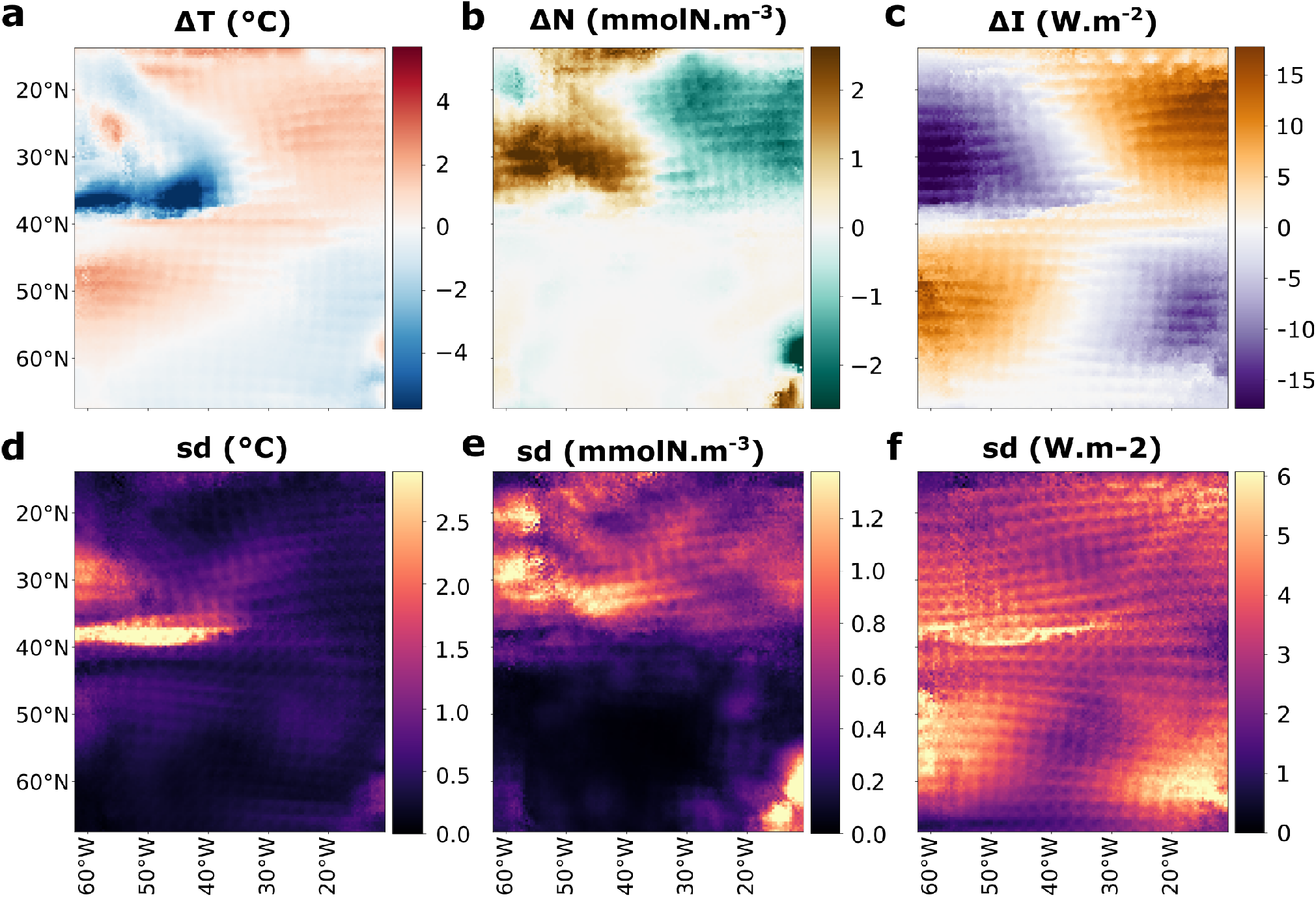
Differences between Lagrangian environmental histories and fixed-location Eulerian histories at particle arrival points. Median offsets between the Lagrangian and fixed-location Eulerian histories of environmental conditions, calculated at the grid point reached by each particle after 90 days of travel, for (**a**) temperature, (**b**) nitrate, and (**c**) light. (**d**–**f**) Standard deviations of the corresponding Lagrangian offsets at each grid point. All plots shown are for the simulation with *V*_*ref*_ = 1*L*.

### Lagrangian histories reveal environmental sensitivity and population bottlenecks

We further examined how Lagrangian trajectories and environmental histories varied with particle lifespan (Figure 8, Figure S16, and S17). Short- and intermediate-lifespan particles were biased near species’ niche edges and at higher latitudes (Figure 8a,b), whereas long-lived particles were more concentrated at intermediate latitudes and longitudes in both gyres (Figure 8c). Mortality peaked in August for short-lived particles and in June for intermediate-lived particles, consistent with the seasonal decline following the spring–summer bloom (Figure 8d–f). Intermediate-lived particles had higher mean biomass along their trajectories than the short-lived ones, whereas long-lived particles spanned a wider, bimodal range of biomass levels and were often located in the southern gyre. This suggests that persistence may be supported by relatively stable conditions in the southern gyre and consistently favorable conditions along trajectories in the northern gyre (Figure 8g–i). Environmental histories also differed with lifespan (Figure S16). Short-lived particles generally experienced low temperatures, nutrient concentrations, and light exposure, although a second peak of short-lived high-light exposure particles occurred in the Senegal upwelling region. Intermediate-lived particles experienced higher mean nutrient concentrations and light exposure than the short-lived while long-lived particles showed a bimodal pattern: the northern gyre particles experienced low-to-intermediate temperatures, high nutrient concentrations, and intermediate light exposure, whereas those in the southern gyre experienced higher temperatures and light exposure but lower nutrient concentrations. Overall, these results suggest that environmental histories along particle trajectories may contribute to population persistence in the PBM. Relative to species’ optima, short- and intermediate-lived particles generally experienced negative temperature anomalies and a skewed distribution toward negative nitrate anomalies (Figure S17). Long-lived particles instead had both positive and negative temperature and near-zero to positive nitrate anomalies, with bimodal distributions. These patterns suggest that temperature mismatches may contribute to mortality among many short-lived particles, while nutrient depletion may affect a smaller subset.

**Figure 8:**
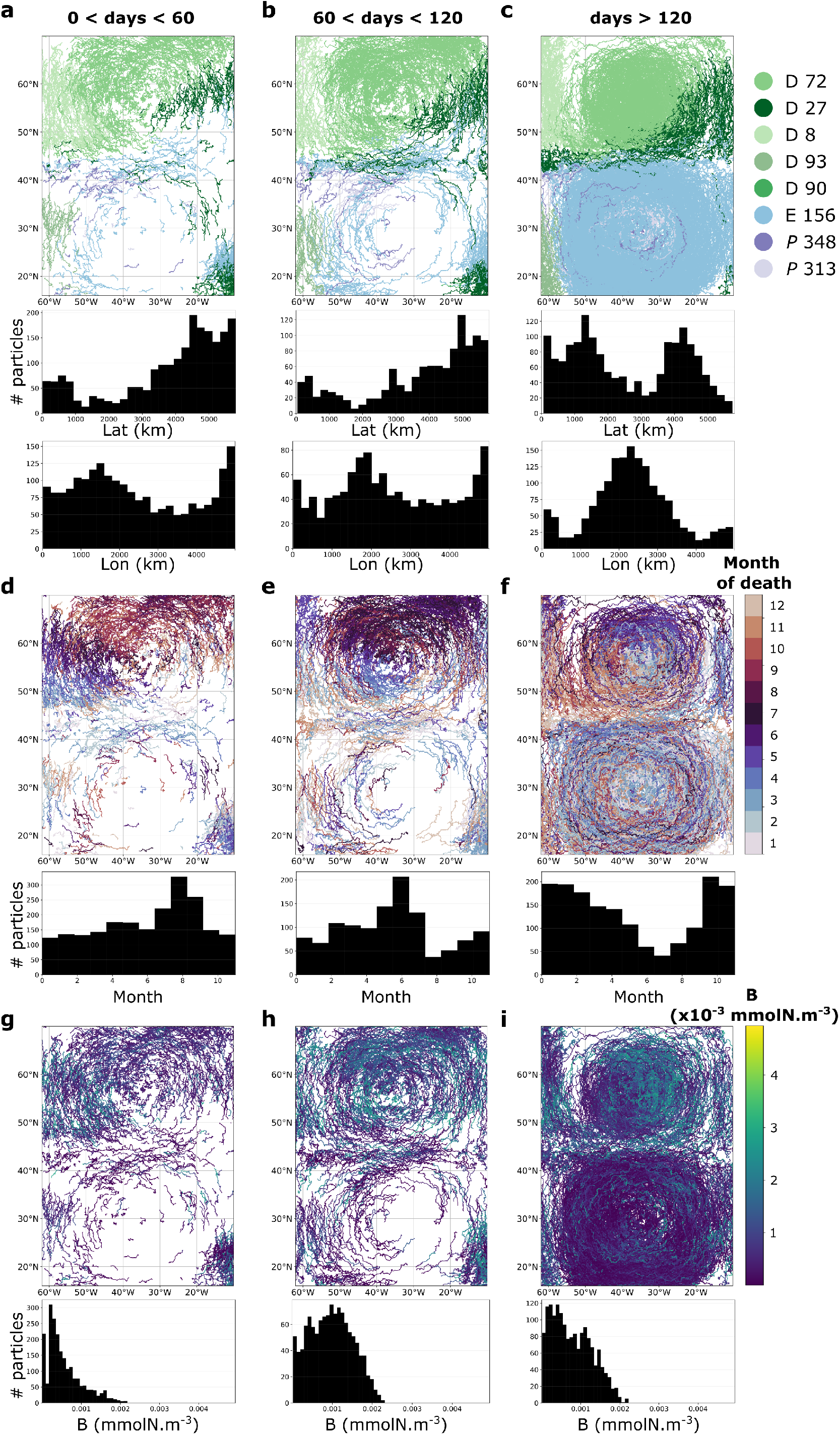
Lagrangian histories and trajectories as a function of particle lifespan. (**a**–**c**) Lagrangian trajectories colored by species, with distributions of mean latitude and longitude for particles grouped by lifespan. (**d**–**f**) Lagrangian trajectories colored by month of death, with distributions of month of death for particles grouped by lifespan. (**g**–**i**) Particle biomass trajectories, with distributions of mean biomass for particles grouped by lifespan. All plots shown are for the simulation with *V*_*ref*_ = 1*L*.

Finally, we compared spatial patterns of Lagrangian and Eulerian mortality (Figure S18). In the Lagrangian PBM, mortality hotspots in the northern gyre coincided with the annual phytoplankton bloom collapse and sharp declines in biomass and particle numbers. Eulerian mortality, by contrast, was concentrated along competition fronts, reflecting differences in how the two formulations translate population dynamics into mortality patterns. As the reference volume increased, Eulerian mortality fronts weakened slightly, while the PBM hotspot in the northern gyre became more pronounced (Figure S18g–i), suggesting a stronger diversity bottleneck in its southern part and enhanced coexistence elsewhere (Figure 3). Particle division numbers and Lagrangian trajectories distances also varied between the two gyres, further reflecting differences in particle histories across the domain (Figure S19). Overall, these results highlight differences in the mortality dynamics in the two model including its spatial distribution and suggest that particle-level dynamics shape population bottlenecks and coexistence in the PBM.

## Discussion

We compared a particle-based, size-structured phytoplankton model in an idealized North Atlantic double gyre flow with its Eulerian counterpart to assess differences in ecological dynamics, biogeography, richness, diversity, and environmental histories. The PBM represents phytoplankton as advected Lagrangian particles with particle-level deterministic mortality, in contrast to the grid-based mortality formulation of the Eulerian model. In the PBM, we found more robust dynamics and biogeographical patterns compared to the Eulerian model independently from the mortality threshold chosen. These contrasting behaviors were likely driven by the Lagrangian formulation of the PBM and its particle-based representation of mortality. In the PBM, the peak of the spring/summer bloom was higher than in the Eulerian model, suggesting that Lagrangian variability can create more intense biomass accumulation that the Eulerian approach tended to smooth out. This is consistent with previous comparisons showing that Lagrangian or particle-based formulations allow individual phytoplankton to experience heterogeneous environmental histories and respond on biological time scales that are faster than the grid-averaged dynamics captured by Eulerian models (Baudry et al. 2018). This tendency parallels high-resolution individual- and Lagrangian particle-based formulations in atmospheric and oceanographic modeling, where tracking individual traits and histories preserves fine-scale kinetic and biological variance, better capturing localized extreme events and tail-risk dynamics compared to smoothed Eulerian continuum models (Chenillat et al. 2015; Hellweger, Clegg, et al. 2016; van Sebille et al. 2018; Li et al. 2022; Morrison et al. 2024). The PBM also showed a stronger seasonality in the northern Atlantic gyre, suggesting more sensitivity to bloom termination. Comparison with satellite-derived chlorophyll concentrations (Sauzéde et al. 2025) showed strong agreement between model and observations for both approaches, although with differing biases. This agreement likely reflects both the constraint imposed by the observed nitrate field on phytoplankton carrying capacity and the ecological dynamics governing how biomass responds to environmental conditions.

In the highly seasonal northern gyre, the PBM generally produced higher time-averaged local richness and diversity than the Eulerian model, except at high mortality threshold where they converged. In the more oligotrophic southern gyre, local richness and Shannon diversity increased with particle number without reaching a plateau. Richness was higher or lower than in the Eulerian model depending on the mortality threshold while Shannon diversity reached lower levels in the PBM. These contrasting patterns may reflect differences in the mechanisms supporting coexistence in the two gyres. In the northern gyre, particle-level trajectories may better take advantage of seasonal environmental variability to promote coexistence, whereas in the southern gyre, coexistence may be more sensitive to biomass discreteness through the stochastic persistence of rare types. The distribution of long-lived particles provides a supporting perspective: they occurred primarily in the central regions of both gyres, potentially benefiting from consistently favorable environmental conditions, maintaining high biomass along their trajectories in the northern gyre and from more stable conditions in the southern gyre. Such persistence may contribute to the maintenance of local diversity. Nevertheless, the relative roles of environmental variability, competitive interactions and stochastic persistence remain unclear. The absence of richness saturation with the number of modeled particle suggests that, under weak environmental forcing in this region, local coexistence may therefore reflect a balance between the stochastic persistence of rare types and competitive exclusion, rather than deterministic niche separation alone (Chesson 2000; Pedrós-Alió 2012). Together, these results suggests that discrete biomass representation fundamentally alters emergent coexistence patterns in model plankton ecosystems. The differences arise from discreteness, which acts both through particle-level mortality and through the finite sampling of transport by particles. Furthermore, particle discreteness might capture the patchy distribution of phytoplankton at sub-grid scales (Martin 2003; d’Ovidio et al. 2010; Robinson et al. 2021), particularly near range margins where populations may be sparse and spatially heterogeneous, although the degree of patchiness relevant to population persistence remains uncertain. In our model, absolute richness differs from Dutkiewicz et al. 2020, and from studies emphasizing dispersal-driven diversity (Clayton et al. 2013; Lévy et al. 2015), reflecting differences in niche dimensions, ecological interactions, and extinction criteria. How these differences might change in the PBM with increasing ecological complexity, and whether an explicit extinction threshold is necessary to avoid the numerical “atto-fox” effect, remain open questions.

Beyond its potential role in enhancing coexistence, the PBM’s explicit representation of discrete biomass provides insights into the variability of Lagrangian environmental histories under environmental and competitive pressures. Our results reproduce previously reported differences between Lagrangian temperature histories and local Eulerian conditions (Doblin and Sebille 2016), while extending this comparison to nitrate and light. The magnitude and spatial distribution of these differences varied among environmental factors, with stronger nutrient variability in the northern gyre while temperature and light showed more widespread effects across both gyres. Discrete representation of phytoplankton was also associated with a population bottleneck in the southern part of the northern gyre, whose intensity depended on the mortality threshold. Together, these findings raise questions of phytoplankton adaptation along ocean currents. Adaptation might enable populations to better follow environmental variations they encounter while population bottlenecks might influence the adaptive potential and the maintenance of coexistence.

Several simplifications in our model should be considered when interpreting our results. First, forcing fields of temperature and nitrate, although varying in space and time, are prescribed from climatologies and are not dynamically coupled respectively to the circulation and phytoplankton uptake. Second, the idealized double-gyre circulation is not dynamically consistent with forcing fields and does not reproduce realistic current dynamics, including western intensification. Future work might therefore couple the model to dynamically consistent circulation and biogeochemical fields.

While particle-based models are computationally demanding, they allow to represent processes that Eulerian formulations cannot resolve, including environmental histories, particle level division and mortality, leading to emergent differences in population dynamics and structure. Future PBM-based studies could further incorporate additional mechanisms that are difficult to represent in Eulerian models (Hellweger, Clegg, et al. 2016). The present framework could be complemented with an evolutionary module allowing for neutral genetic drift (Hellweger, Van Sebille, et al. 2014), potentially enabling the reproduction of large-scale patterns of Lagrangian differentiation observed in the global ocean (Richter et al. 2022). While trait-based Eulerian models have demonstrated the central role of environmental variability in shaping phytoplankton resilience to environmental change (Sauterey et al. 2023; Le Gland et al. 2021), understanding the response of plankton communities to ongoing and future climate change remains a major challenge (Frémont et al. 2022). Particle-level adaptive processes could be implemented, for which PBMs are well suited to capture complex, nonlinear evolutionary responses (Clark et al. 2011). Within our framework, such extensions could reveal how adaptation affects coexistence and community assembly, and whether the population bottleneck identified in the northern gyre could constrain the adaptive potential. More broadly, by explicitly resolving Lagrangian trajectories and biomass discreteness, PBMs offer a framework to characterize how environmental histories, stochastic rare-type persistence, and adaptive and evolutionary processes jointly shape plankton communities under changing ocean conditions.

## Supporting information

Videos_1_phyto_biomass

Videos_2_zoop_biomass

Videos_3_richness

Videos_4_shannon

Videos_5_dominant_species

## Acknowledgments

The authors acknowledge the University of Maryland High Performance Computing Cluster (HPCC; http://hpcc.umd.edu), the Genoscope Inti high-performance computing cluster, and the Stazione Zoologica Anton Dohrn (SZN) High Performance Computing (HPC) cluster, which provided the computational resources used to conduct the research reported in this paper. The authors used ChatGPT to assist with coding, grammatical corrections and language clarity. The authors reviewed and edited the AI-generated content as needed and accept full responsibility for the content of the publication.

## Author contributions

Conceptualization: PF, MRA, BHM, DI; methodology: PF; investigation: PF; model development: PF; writing-original draft: PF; writing-review & editing: PF, MRA, BHM, OJ, JSW, DI; funding acquisition: OJ, JSW, DI.

## Data, code, and materials availability

All code and data necessary to generate all figures and tables will be made available at https://github.com/PaulFremont3/phytoPBM.

