## Supplementary Information for "Particle-based representation of phytoplankton modulates emergent coexistence and biogeography in a model ocean"

| Parameter | Symbol | Unit | Value | Reference |
|---|---|---|---|---|
| Basal mortality of phytoplankton | $d$ | $d^{-1}$ | 0.1 | - |
| Maximum grazing rate | $g$ | $(mmolN.m^{-3})^{-1}.d^{-1}$ | 1.0 | Dutkiewicz et al. 2020 |
| Linear zooplankton mortality | $d_z$ | $d^{-1}$ | 0.067 | Dutkiewicz et al. 2020 |
| Quadratic zooplankton mortality | $d_{z2}$ | $(mmolN.m^{-3})^{-1}.d^{-1}$ | 0.45 | Fitted |
| Water light absorption coefficient | $e_p$ | $m^{-1}$ | 0.03 | Fasham et al. 1990 |
| Phytoplankton light absorption coefficient | $e_w$ | $m^2.(mmolN)^{-1}$ | 0.04 | Fasham et al. 1990 |
| Initial slope of the photosynthesis-irradiance (P-I) curve | $\alpha$ | $d^{-1}.(W/m^2)^{-1}$ | 0.1 | Geider et al. 1997; Stawiarski et al. 2018 |
| Maximum $Chl$ to $C$ ratio | $\theta_{max}$ | $gChl.gC^{-1}$ | 0.08 | Geider et al. 1997 |
| Activation temperature | $A_T$ | K | 4000 | Dutkiewicz et al. 2020 |
| Envelope sensitivity | $T_N$ | K | 293.15 | Dutkiewicz et al. 2020 |
| Temperature growth dependency normalization | $\tau_T$ | unitless | 0.8 | Dutkiewicz et al. 2020 |
| Niche width regulator | $B_T$ | $K^{-1}$ | 0.0003 | Dutkiewicz et al. 2020 |
| Niche skewness regulator | $b_T$ | unitless | 4 | Dutkiewicz et al. 2020 |

**Table S1 | Fixed parameters regulating plankton growth independently from their size.**

| Parameter | Symbol | $a_X$ | $b_X$ | Unit | Reference |
|---|---|---|---|---|---|
| Maximum growth rate | $\mu_{max}$ | | | $d^{-1}$ | |
| Diatoms | | 3.9 | -0.08 | | Dutkiewicz et al. 2020 |
| Small eukaryotes | | 1.4 | -0.08 | | Dutkiewicz et al. 2020 |
| *Synechococcus* | | 1.4 | 0.08 | | Dutkiewicz et al. 2020 |
| *Prochlorococcus* | | 0.9 | 0.08 | | Dutkiewicz et al. 2020 |
| Cell nitrate uptake half saturation constant | K | 0.17 | 0.27 | $mmolN.m^{-3}$ | Zakem et al. 2018 |
| Minimum cell quota relative to C | $Q_{min}$ | 0.07 | -0.17 | $mmolN$ $mmolC^{-1}$ | Zakem et al. 2018 |
| Maximum nutrient uptake rate | $V_{max}$ | 0.51 | -0.27 | $mmolN$ $mmolC^{-1}.d^{-1}$ | Zakem et al. 2018 |

**Table S2 | Size dependent parameters regulating phytoplankton growth following allometric laws ($a_X V^{b_X}$).**

### Supplementary methods

#### Particle-based ecological model: supplement

##### Particle growth model: extended equations

The equation of phytoplankton biomass dynamics for a particle of biomass concentration $B_i$ in control grid point $K$ is the following:

$$\frac{dB_i}{dt} = \mu_{max,i}.\psi(T).\gamma(J).\beta(N).B_i - (d.\eta(T) + \frac{B_K}{\tau(T,N,I).L}).B_i - g.\eta(T).H_2(B_K).Z_K.\frac{B_i}{B_K} \tag{S1}$$

$B_i$ is the biomass concentration represented by particle $i$ expressed in $mmolN.m^{-3}$. $\mu_{max,i}$ is the maximum growth rate of the particle $i$ in $d^{-1}$. $\psi(T)$, $\gamma(J)$, $\beta(N)$ are, respectively, the temperature, light, and nutrient limitation terms (unitless, described below). $\psi(T)$ contains a niche temperature term while $\eta(T)$ does not (see below). $d$ is the basal death rate of phytoplankton in $d^{-1}$. $g$ is the maximum grazing rate in $d^{-1}$. $B_K = \sum_{Kj} B_{Kj}$ and $Z_K = \sum_{Kj} Z_{Kj}$ are, respectively, the total phytoplankton and zooplankton biomass concentration in a grid cell $K$. $L$ is a constant in $mmolN.m^{-3}$. Finally, $H_2(B_K)$ is a Holling type III response function (Holling 1959). The equation for the zooplankton biomass $Z_i$ (transported in the same particle as the phytoplankton) is:

$$\frac{dZ_i}{dt} = \epsilon_Z.g.\eta(T).H_2(B_K).Z_i - d_z.\eta(T).Z_i - d_{z2}.\eta(T).Z_K.Z_i \tag{S2}$$

where $d_Z$ in $d^{-1}$ and $d_{z2}$ in $(mmolN.m^{-3})^{-1}.d^{-1}$ are, respectively, the linear and quadratic mortality term associated to the zooplankton and $\epsilon_Z$ is the grazing efficiency (unitless). The quadratic mortality term represents higher order losses due to predation of the zooplankton by higher trophic levels (not explicitly represented). To conserve zooplankton biomass, when a super-individual particle dies (see Particle biomass concentration, duplication and death), the zooplankton biomass that was in the super-individual particle is redistributed homogeneously to nearby super-individual particles from the same control grid point. When no other super-individual particle is present at this grid point, the zooplankton biomass disappears.

##### Particle biomass concentration: supplement equations

In a control grid cell $K$ the total biomass concentration is defined as follows:

$$B_K = \sum_i B_{Ki} \tag{S3}$$

Where $B_{Ki}$ is the biomass concentration of particle $i$. Depending on the depth of the mixed layer, a given particle does not represent the same total biomass: it represents an effective biomass $B_i^{eff}$ in the water column of depth $z$:

$$B_i^{eff} = B_i.z.S \tag{S4}$$

Where $S$ is the area in $m^2$ of a grid cell.

#### Eulerian model: supplement

##### Species growth model: extended equations

From equations S1 and S2, the total biomass, $B_{Ks}$, of a species $s$ of phytoplankton and the zooplankton biomass $Z_K$ in a control grid cell $K$ are, by summing over the particles' equations in this control grid cell (of a given species for the phytoplankton and over all particles for the zooplankton):

$$\frac{dB_{Ks}}{dt} = \mu_{max,i}.\psi(T).\gamma(J).\beta(N).B_{Ks} - (d.\eta(T) + \frac{B_K}{\tau(T,N,I).L}).B_{Ks} - g.\eta(T).H_2(B_K).Z_K.\frac{B_{Ks}}{B_K} \tag{S5}$$

$$\frac{dZ_K}{dt} = \epsilon_Z.g.\eta(T).H_2(B_K).Z_K - d_z.\eta(T).Z_K - d_{z2}.\eta(T).Z_K^2 \tag{S6}$$

#### Phytoplankton growth model: supplement

##### Chlorophyll model: equations

We modeled chlorophyll dynamics following Geider et al. (1997) and the Redfield ratio (Redfield 1934):

$$\frac{dChl_i}{dt} = \mu_{eff,i}.C_i.\rho_{chl} - d_{chl}.Chl_i - d_{zoo}.\frac{C_i}{C_K}.\frac{Chl_i}{C_i} \tag{S7}$$

Where $Chl_i$ is the chlorophyll concentration of particle $i$ in $mgChl.m^{-3}$, $C_K$ is the total carbon concentration in grid cell $K$, $C_i = B_i.\frac{106}{16}.MMC$ is the carbon biomass of particle $i$ (in $mgC.m^{-3}$), $\mu_{eff,i} = \mu_{max,i}.\psi(T).\gamma(J).\beta(N)$ is the effective growth rate, $d_{chl} = d.\eta(T) + \frac{B_K}{\tau(T,N,I).K}$ is the effective death rate (excluding zooplankton grazing), and $d_{zoo} = g.\eta(T).H_2(B_K).Z_K^C$ is the grazing term of zooplankton with $Z_K^C = Z_K.\frac{106}{16}.MMC$ (in $mgC.m^{-3}$). $\rho_{chl}$ is the ratio of chlorophyll A synthesis to carbon fixation based on equation 4 from Geider et al. (1997):

$$\rho_{chl} = \frac{\theta_{max}^2.\mu_{eff,i}}{\theta.\alpha.J} \tag{S8}$$

Where $\theta_{max}$ is the maximum $Chl$ to $C$ ratio in $gChl.gC^{-1}$, $\alpha$ is the initial slope of the photosynthesis-irradiance (P-I) curve in $d^{-1}(W/m^2)^{-1}$, $J$ is the level of light at the surface at the zenith in $W.m^{-2}$. Note that we had to substitute $\alpha^{chl}$ by $\frac{\alpha}{\theta_{max}}$ given our definition of $\alpha$ compared to Geider et al. 1997.

#### Growth equilibrium: derivation

Equating equation S1 to 0 without the zooplankton mortality term, we get:

$$B_K = \tau(T,N,J).L.(\mu_{max,i}.\psi(T).\gamma(J).\beta(N) - d.\eta(T)) \tag{S9}$$

At equilibrium, we set $B_K = N$, where $N$ is the nutrient concentration in the control grid cell $K$. This assumes that all available nutrients could be consumed. Equation S3 yields:

$$\tau(T,N,J) = \frac{N}{L.(\mu_{max,i}.\psi(T).\gamma(J).\beta(N) - d.\eta(T))} \tag{S10}$$

#### Temperature dependency and niche: equations

For the temperature dependency $\psi(T)$ (growth rate dependency including a thermal niche term) and $\eta(T)$ (mortality and grazing dependency to temperature), we follow Dutkiewicz et al. 2020 by using the Eppley curve (Eppley 1972):

$$\psi(T) = \tau_T.\exp(-A_T.(\frac{1}{T} - \frac{1}{T_N})).\exp(-B_T.|T - T_{opt,i}|^{b_T}) \tag{S11}$$

$$\eta(T) = \tau_T.\exp(-A_T.(\frac{1}{T} - \frac{1}{T_N})) \tag{S12}$$

$T_{opt,i}$ is the optimal temperature of species $i$. $A_T$, $T_N$, $B_T$, and $b_T$ regulate the shape of the temperature dependency curve (see Table S1 for values). More precisely, $\tau_T$ and $T_N$ regulate the absolute values of $\psi(T)$, $A_T$ regulates the sensitivity to temperature, $B_T$ regulates the niche width and $b_T$ controls the smoothness of the niche borders. For simulations, each species is randomly assigned to one of ten linearly spaced thermal norms between -2 and 34°C (Figure S1c,d).

#### Light dependency: equations

For light dependency $\gamma(J)$ we followed Evans and Parslow 1985 and Stec 2016:

$$\gamma(J) = \frac{2}{H.k}.(F(\Psi.\exp(k.H),\ \frac{\tau_0}{2}) - F(\Psi,\frac{\tau_0}{2}) - F(\Psi.\exp(k.H),\ 0) + F(\Psi,\ 0)) \tag{S13}$$

Where $F$ is the following function:

$$F(y,t) = \sqrt{y^2 + t^2} - t.ln(\frac{t + \sqrt{y^2 + t^2}}{y}) \tag{S14}$$

$\Psi$ results from the transformation of vertical coordinate from depth to reciprocal light, given by $\Psi = \frac{\mu_{max,i}.\frac{\tau_0}{2}}{\alpha.J}$ and $y = \Psi.exp(kz)$ with $dz = \frac{dy}{k.y}$ (see Evans and Parslow 1985 and Stec 2016 for details of the calculation). $\frac{\tau_0}{2}$ is half the duration of the day in day (*i.e.* the time, in day, between sunrise and zenith), $\alpha$ is the initial slope of the photosynthesis-irradiance (P-I) curve in $d^{-1}.(W/m^2)^{-1}$, $J$ is the level of light at the surface at the zenith in $W.m^{-2}$. $k$ is the light attenuation coefficient in $m^{-1}$ that accounts for the water light absorption and self-shading:

$$k = e_p + e_w.\sum_{jK} Bj \tag{S15}$$

$e_p$ is the water light absorption coefficient in $m^{-1}$, $e_w$ is the phytoplankton light absorption coefficient in $m^2.(mmolN)^{-1}$ (self shading effect). Finally, in equation S11, $H$ is the Mixed Layer Depth (MLD) in $m$ (see Mixed Layer Depth field).

##### Nutrient dependency: equations

For nutrient dependency $\beta(N)$, we use the limitation term of the Monod equation of growth (Monod 1949):

$$\beta(N) = \frac{N}{N + N_{c,i}} \quad \text{(S16)}$$

Where $N$ is the nitrate concentration at the given location (control grid cell). $N_{c,i}$ the half saturation constant for growth of particle $i$, associated to a given species:

$$N_{c,i} = \frac{HS_i.\mu_{max,i}.Q_{min,i}}{V_{max,i}} \quad \text{(S17)}$$

Where $HS_i$ is the cell nutrient uptake half saturation constant, $Q_{min,i}$ is the cell minimum stoichiometric quota, $V_{max,i}$ is the maximal cell nutrient uptake rate. For all three parameters, an empirical allometric relationship with the cell volume $V$ has been observed (Dutkiewicz et al. 2020; Zakem et al. 2018):

$$HS_i = a_K.V^{b_K};\ Q_{min,i} = a_Q.V^{b_Q};\ V_{max,i} = a_V.V^{b_V} \quad \text{(S18)}$$

##### Elemental quotas

We focus on four different types of phytoplankton: *Prochlorococcus*, *Synechococcus*, non diatom eukaryotes and diatoms. The mortality threshold model we develop necessitates to fix their sizes to define their elemental quotas, *i.e.* the amount of nitrogen in one phytoplankton cell. For eukaryotes, we obtain nitrogen cell quotas following Menden-Deuer and Lessard (2000) and the Redfield ratio (Redfield 1934):

- Diatoms:

$$log_{10}(Q_{diatom}(C)) = (-0.541 + 0.811.log_{10}(Vol)), \quad \text{(S19)}$$

- Eukaryotic protists excluding diatoms:

$$log_{10}(Q_{phytoeukaryotes}(C)) = (-0.665 + 0.939.log_{10}(Vol)), \quad \text{(S20)}$$

where $Vol$ is the cell volume. We then convert to nitrogen units ($\mu molN.cell^{-1}$):

$$Q(N) = Q(C).10^{-6}.\frac{1}{MMC}.\frac{16}{106}, \quad \text{(S21)}$$

where $\frac{16}{106}$ is the Redfield ratio of nitrogen to carbon and $MMC$ is the molar mass of carbon. For cyanobacteria, *Synechococcus* and *Prochlorococcus*, we use the carbon density of *Synechococcus* from Verity et al. (1992):

$$Q_{cyanobacteria}(N) = d.Vol.10^{-9}.\frac{1}{MMC}.\frac{16}{106}, \quad \text{(S22)}$$

where $d$ is the mass density of *Synechococcus*: d=470 fgC.$\mu$m$^{-3}$ (Verity et al. 1992) and consistent with other measurements for *Prochlorococcus* (Roth-Rosenberg et al. 2021).

##### Grazing limitation: equation

For grazing limitation, we choose a Holling type III limiting function (Holling 1959) commonly used in biogeochemical models (Zahariev et al. 2008; Law et al. 2017; Dutkiewicz et al. 2020):

$$H_2(B_K) = \frac{B_K^2}{B_K^2 + S_c^2}, \quad \text{(S23)}$$

where $B_K$ is the total phytoplankton concentration in a grid cell and $S_c$ is the half saturation concentration for grazing in $mmolN.m^{-3}$.

#### Idealized North Atlantic double gyre flow: supplement

##### Main flow: equations

We build an idealized double gyre flow consisting of two principal gyres (north and south) with the dimensions of the real North Atlantic double gyre and static eddies. The well known and extensively studied double gyre equation (Shadden et al. 2005; Froyland and Padberg 2009; Froyland and Padberg-Gehle 2015; Banisch and Koltai 2017; Reijnders et al. 2021) is used to mimic the double gyre. The speed potential is given by:

$$\Psi_{dg}(x, y) = A_{dg}. \sin(\pi.\omega_x.x). \sin(\pi.\omega_y.y) \tag{S24}$$

From equation S14, respectively meridional and zonal flow speeds are derived:

$$u_{dg} = \frac{\partial \Psi_{dg}(x, y)}{\partial y} = -A_{dg}.\pi.\omega_y. \sin(\pi.\omega_x.x). \cos(\pi.\omega_y.y) \tag{S25}$$

$$v_{dg} = -\frac{\partial \Psi_{dg}(x, y)}{\partial x} = A_{dg}.\pi.\omega_x. \cos(\pi.\omega_x.x). \sin(\pi.\omega_y.y) \tag{S26}$$

$\omega_x$ is the spatial frequency of the gyre in $m^{-1}$ (*i.e.* it characterize its size in the longitudinal direction), here $\omega_x = \frac{1}{5.10^6}$. It corresponds approximately to the width of the North Atlantic Ocean between the french city of Bordeaux to the Canadian province of Nova Scotia. $\omega_y$ is the equivalent in the latitudinal direction, here $\omega_y = \frac{1}{5.8.10^6}$. It corresponds approximately to half the distance of a transect between the Senegal upwelling east of Dakar and the north of Iceland (54° of latitude). $A_{dg}$ characterizes the maximum speed of the double gyre velocity field, $A_{dg} = \frac{u_{\max}}{\pi.\omega_x}$, we set $u_{\max} = 0.15\ m.s^{-1}$.

#### Mesoscale static eddies: equations

In addition to the main double gyre, we approximately model mesoscale eddies using the same equations with different scale parameters. Eddies are here static but oscillate in time (resulting in a periodic change in the direction of their flow).

$$\Psi_e(x, y) = A_e. \sin(\pi.\omega_{ex}.x). \sin(\pi.\omega_{ey}.y).cos(\pi.\omega_t.t) \tag{S27}$$

$$u_e = \frac{\partial \Psi_e(x, y)}{\partial y} = -A_e.\pi.\omega_{ey}. \sin(\pi.\omega_{ex}.x). \cos(\pi.\omega_{ey}.y).cos(\pi.w_t.t) \tag{S28}$$

$$v_e = -\frac{\partial \Psi_e(x, y)}{\partial x} = A_e.\pi.\omega_{ex}. \cos(\pi.\omega_{ex}.x). \sin(\pi.\omega_{ey}.y).cos(\pi.w_t.t) \tag{S29}$$

$\omega_{ex}$ and $\omega_{ey}$ are the spatial frequencies of the eddies in m$^{-1}$ (*i.e.* it characterizes eddies size in the longitudinal and latitudinal direction), here $\omega_{ex} = \omega_{ey} = \frac{1}{250.10^3}$ $A_e$ characterizes the maximum speed of the eddies velocity field, $A_e = \frac{u_{e\,\max}}{\pi.\omega_{ex}}$, we set $u_{\max} = 0.15\ m.s^{-1}$. $\omega_t$ is the temporal frequency of the eddies which we set to a period of 2 *months*. The global speed field is then the addition of the double gyre and the eddies.

$$u = u_{dg} + u_e \tag{S30}$$

$$v = v_{dg} + v_e \tag{S31}$$

The addition of the two field allows to reach a maximum speed of approximately 0.3 $m.s^{-1}$ which is what is observed in the real oceans except in the Gulf Stream where higher speeds are reported (Marzocchi et al. 2015).

#### Diffusion: equations

In addition to the main velocity field, we model small scale turbulence by adding a 2 dimensional Brownian motion to the particles. Brownian motion is modeled as follows: at each time step the particle is transported in a random direction of the 2 dimensional grid according to the diffusion equation with a diffusion coefficient $D$ in $m^2.s^{-1}$. Accordingly, we have at any integration time step t:

$$x(t + 1) = x(t) \pm U(0, 1).\sqrt{\frac{2.D.dt}{r}} \tag{S32}$$

$$y(t + 1) = y(t) \pm U(0, 1).\sqrt{\frac{2.D.dt}{r}} \tag{S33}$$

Where $U(0, 1)$ is the uniform distribution between 0 and 1. $r$ is equal to $\frac{1}{3}$ to account for the bi-dimensional nature of our field (Delandmeter and Sebille 2019). This renders the global field described in equations S26 and S27 stochastic and particle dependent. By derivation over time of equation S28 and S29 we have $|u_b| = |v_b| \sim U(\frac{1}{2.dt}.\sqrt{\frac{2.D.dt}{r}}, \frac{2.D}{12.r.dt}) = U(\sqrt{\frac{3.D}{2.dt}}, \frac{D}{2.dt})$, where U is the uniform distribution. We used a diffusion coefficient of 1000 $m^2.s^{-1}$ (both in the Eulerian and PBM model). The average speed due to diffusion is around 0.64 $m.s^{-1}$ and the standard deviation is around 0.14 $m.s^{-1}$ (for the time step of 1 hour *i.e.* 3600 s used to integrate particle trajectories and Eulerian advection and diffusion).

##### Particle speed: equations

For a particle $i$ we finally have:

$$u_i(x,y) = u_{dg} + u_e + u_b \tag{S34}$$

$$v_i(x,y) = v_{dg} + v_e + v_b \tag{S35}$$

##### Particles transport

Particles were advected and diffused in a Lagrangian framework using the python library Parcels (Delandmeter and Sebille 2019) within a 125x145 2 dimensional grid (square grid point of approximately 40x40 kms). For integration over time we built a custom kernel consisting of the addition of 2 kernels: AdvectionDiffusionEM from Parcels for the advection and a custom boundary kernel. The custom boundary kernel brings back out of domain particles in the flow field by bouncing back these particles ensuring its total displacement is the same. This is achieved by setting the particle's coordinate as its mirror coordinate compared to the boundary that it crossed. A set of 142 million particle trajectories over 1 year was precomputed for the PBM simulations, allowing trajectories to be reused across simulations and every year of each simulation. A substantially larger number of trajectories than the maximum number of simulated particles was required, as additional "free" (i.e unassigned to a species) trajectories were needed to enable particle division. Daughter cells were assigned to "free" trajectories in the same grid point as the parent cell while the parent cell trajectory was kept. Advection was performed with a 1 hour time step and particles' positions were saved every day.

#### Physical and geochemical forcings: supplements

##### Light field: equations

We used a simplified radiation equation from Stec 2016:

$$J(t,lat) = I_0(t).cosineeffect(t,lat).f_{par}.(1-albedo).C_{fac} \tag{S36}$$

Where $I_0(t)$ is the solar radiation at day t of the year that depends on the earth-sun distance (Duffie and Beckman 2013):

$$I_0(t) = 0.7.I_{sc}.(1+0.034.\cos\frac{2.\pi.t}{365}) \tag{S37}$$

$I_{sc}$ is the solar constant (1367 $W/m^2$). The cosine effect, $cosineeffect(t,lat)$, takes into account the variation in inclination of irradiance rays with latitude and day of the year:

$$cosineeffect(t,lat) = \cos\frac{\alpha(t,lat).\pi}{180} \tag{S38}$$

Where $\alpha(t)$ is the solar elevation of the sun at zenith dependent on day of the year and latitude, $lat$ (Kirk 1983):

$$\alpha(t,lat) = 90 - \arcsin(\sin\frac{\delta(t).\pi}{180}.\sin\frac{lat.\pi}{180} + \cos\frac{\delta(t).\pi}{180}.\cos\frac{lat.\pi}{180}).\frac{180}{\pi} \tag{S39}$$

$\delta(t)$ is the solar declination along the year from Spencer 1971:

$$\delta(t) = 0.39637 - 22.9133\cos\psi_d + 4.02543\sin\psi_d - 0.3872\cos 2\psi_d + 0.052\sin 2\psi_d \tag{S40}$$

$\psi_d$ is the day of the year expressed as an angle: $\psi_d = \frac{360.t.\pi}{365.180}$ , $t$ being the day of the year. Finally, in equation S32, $f_{par} = 0.43$ is the fraction of solar irradiance between 400 and 700 nm (used for photosynthesis). $albedo = 0.04$ is the albedo of the sea surface. $C_{fac}$ is the effect of cloud on the atmosphere transmission (unitless).

$$C_{fac} = 1 - 0.0775W + 0.0019(90 - \alpha(t,lat)) \tag{S41}$$

$W = 6$ is the cloud fraction in oktas.
The day length (Kirk 1983) is formulated as:

$$\tau_0 = 0.133.\arccos(-\tan\frac{lat.\pi}{180}\tan\frac{\delta(t).\pi}{180}) \tag{S42}$$

##### Mixed Layer Depth field: extended

We built a daily MLD climatology over our spatial domain (125x145 spatial grid) using a regular 2°×2° grid temperature-based global scale MLD climatology from the IFREMER (De Boyer Montégut et al. 2004). The oceanic Mixed Layer is the part of the surface ocean that is considered well mixed and homogeneous either for temperature or sigma-t (density). Monthly means in the considered area of the North Atlantic were used. For the southern gyre, we use datapoints within the rectangle delimited by isolatitudes 12°N to 38°N and by isolongitudes 16°W to 68°W. For the northern gyre, we use isolatitudes 40°N to 66°N and isolongitudes 10°W to 62°W. To fill gaps where there was land in reality but not our gyre, we used the na.approx function from the R package zoo (Zeileis and Grothendieck 2005) latitudinally then longitudinally (using the transpose of the matrix). Then to increase the resolution to the 125x145 grid we linearly interpolated the whole grid using the interp.surface.grid function from the R package fields (Nychka et al. 2017). Finally, we used the linspace function from R package pracma (Borchers 2017) to interpolate linearly the MLD each day of the year and at each grid points. The dynamical Mixed Layer field is used to compute the temperature field, the nutrient field and the light dependency for growth. The biomass of a given particle in a grid square represents a fraction of the concentration of a species within this grid integrated over the Mixed Layer Depth (MLD).

##### Temperature and nitrate fields: extended

- Main fields: extended

We built a daily temperature and nitrate fied over our spatial grid (125x145 grid). To better match real *in situ* data, we use the 1°x1° monthly data from the World Ocean Atlas 2018 (Locarnini et al. 2018; Garcia et al. 2018) as forcing fields. For temperature, we used the 2005-17 climatology, while for nitrate all decades were used. The same North Atlantic boundaries as for the MLD field were used. We used exactly the same steps as for MLD for each depth of the WOA18: first missing values from land were extrapolated, then the resolution is increased to the 125x145 grid and values were extrapolated to the whole year. Finally, weighted mean (taking into account in-homogeneous depth intervals from WOA18) were calculated using the MLD field which rendered the final 2D dynamic fields of temperature and nitrate. In the dynamical ecological model, both temperature and nitrate were independent from the flow field and any feedback from the ecology.

- Eddies nutrient field: extended

In addition to the main nutrient field, we added a nutrient field proportional to the natural logarithm ($ln$) of the eddies field. In the northern gyre, nutrient eddies field was added to the main nutrient field while it was subtracted in the southern gyre following the paradigm that cyclonic and anticyclonic eddies respectively increase and decrease the accumulation of phytoplankton biomass (McGillicuddy 2016). We avoided negative nutrient concentration by setting a very low minimum ($10^{-4}$ $mmolN.m^{-3}$). Nitrate concentration supplied by the eddies in the northern gyre is set between $10^{-6}$ (between eddies) and 2 $mmolN.m^{-3}$ (eddies center). In the southern gyre, nutrient concentration depleted by the eddies in the northern gyre is set between $10^{-6}$ (between eddies) and 0.05 $mmolN.m^{-3}$ (eddies center).

#### Simulations: implementation

The particle-based model (PBM) was implemented in C++ and parallelized across eight CPU cores using multiprocessing. The longest run (corresponding to the highest particle number modeled and highest $V_{ref}$) took approximately 4.5 days to complete. Particles trajectories and division were updated with a 1 day time step (note that Lagrangian trajectories were initially integrated and recorded with a 1 hour timestep like the Eulerian model) while biological growth was updated with a 1 hour time step. The Eulerian model was implemented in Python using Numba for speedup and executed on a single CPU core. Spatial integration of advection and diffusion was computed using the midpoint technique while biomass growth was computed using the Euler method both using a 1 hour integration time step. Simulation runtimes were around 2.5 days.

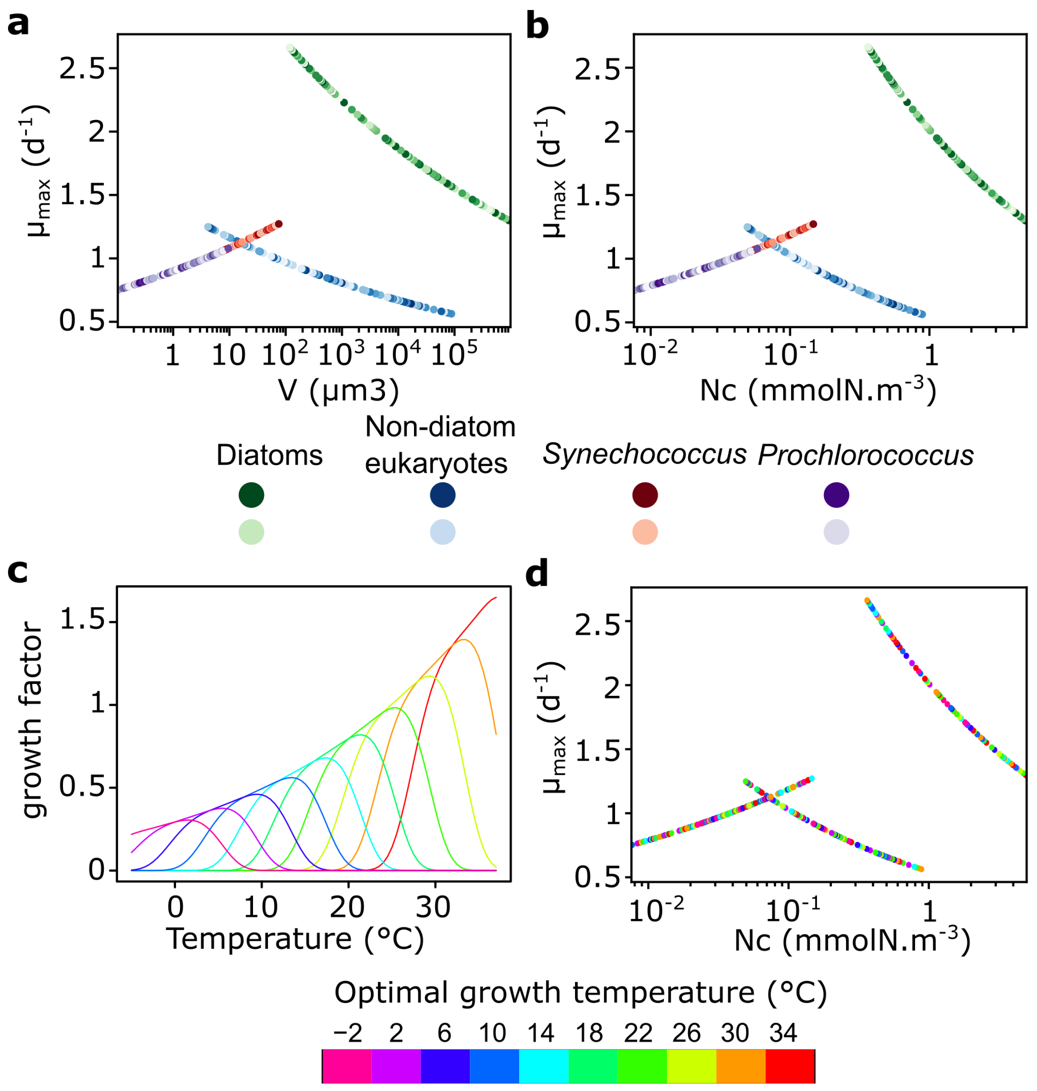


**Figure S1 | Allometric relationships and thermal envelopes defining the life-history traits of 400 randomly generated initial species.** (**a**) Allometric relation defining the maximum growth rate in function of cell volume for cyanobacteria (purples: *Prochlorococcus*, oranges: *Synechococcus*), diatoms (greens) and other eukaryotes (blues). (**b**) Allometric relation of the maximum growth rate in function of the half saturation constant for nitrate for cyanobacteria (purples: *Prochlorococcus*, oranges: *Synechococcus*), diatoms (greens) and other eukaryotes (blues). (**c**) Growth response curves of the ten modeled thermal niches. (**d**) Same as (**b**) with each species colored according to its optimal growth temperature drawn randomly from the ten optimal growth temperature.

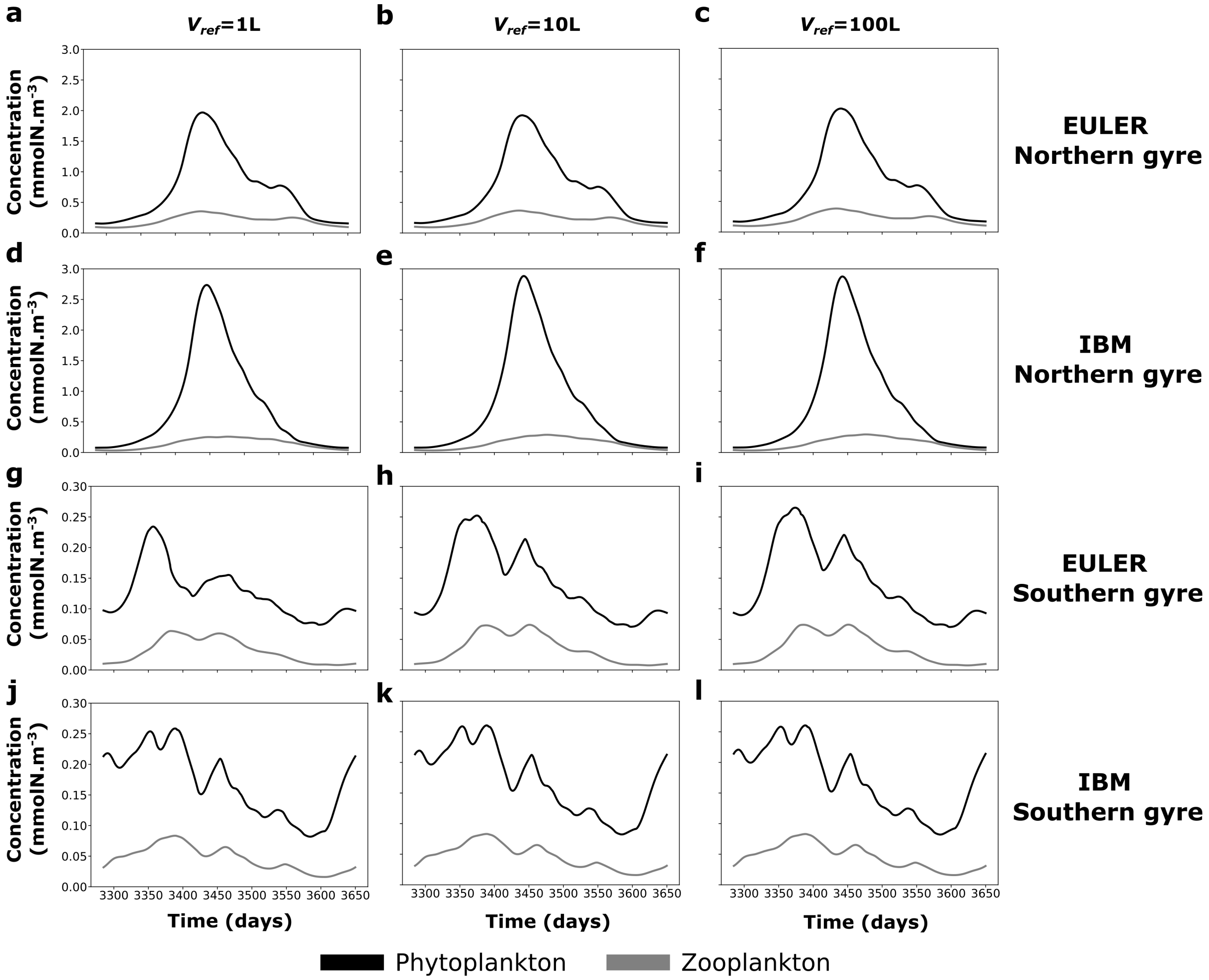


**Figure S2 | Time series of average concentration of total phytoplankton and zooplankton in the last year of simulation in the northern and southern gyres for the two different models and different mortality-threshold reference volumes.** Eulerian model in the northern gyre for (**a**) $V_{ref} = 1L$, (**b**) $V_{ref} = 10L$ and (**c**) $V_{ref} = 100L$. PBM model in the northern gyre for (**d**) for $V_{ref} = 1L$, (**e**) $V_{ref} = 10L$ and (**f**) $V_{ref} = 100L$. Eulerian model in the southern gyre for (**g**) $V_{ref} = 1L$, (**h**) $V_{ref} = 10L$ and (**i**) $V_{ref} = 100L$. PBM model in the southern gyre for (**j**) $V_{ref} = 1L$, (**k**) for $V_{ref} = 10L$ and (**l**) $V_{ref} = 100L$.

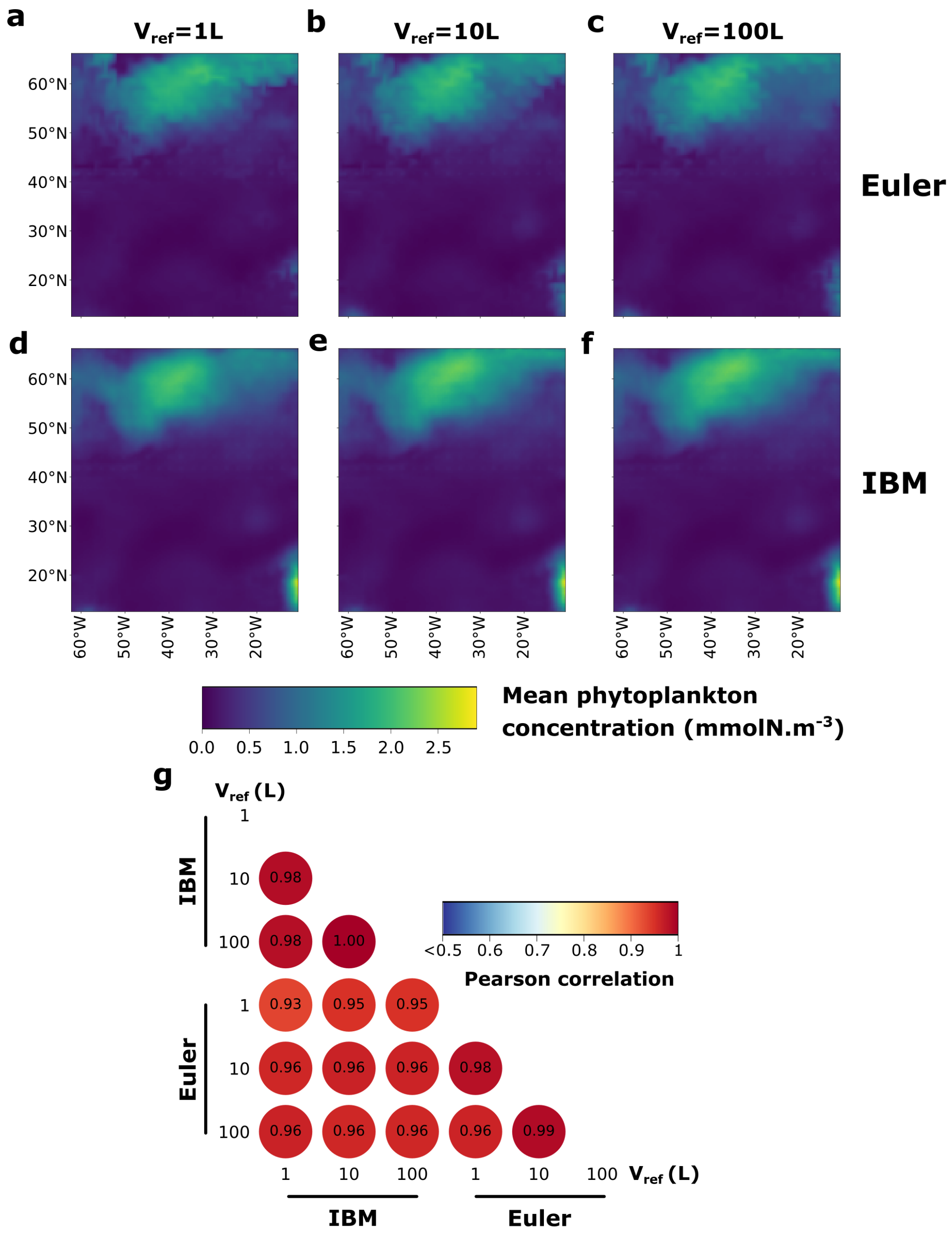


**Figure S3 | Average biomass concentration of the Eulerian and the PBM model for different mortality-threshold reference volumes.** (**a-f**) Average biomass on average at each grid point for three different mortality-threshold reference volume for (**a-c**) the Eulerian model and (**d-f**) the PBM model. (**g**) Pairwise correlogram of average biomass across models and reference volumes.

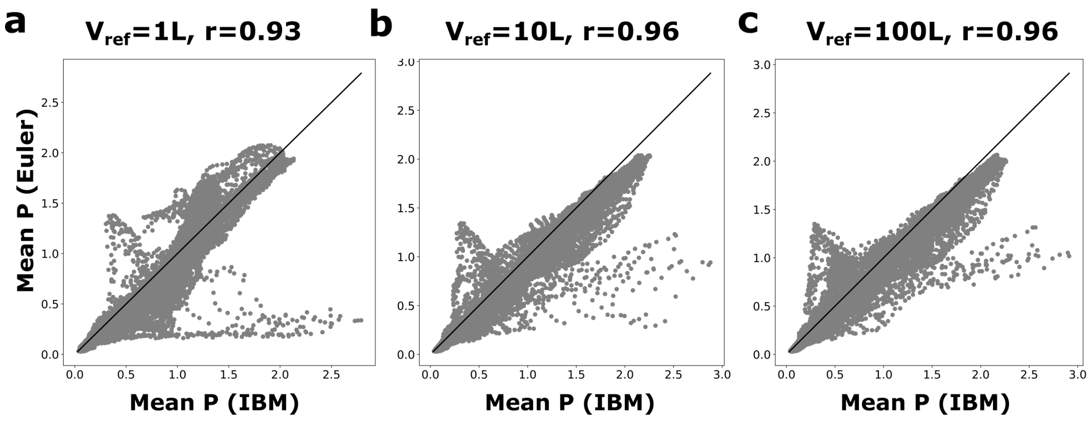


**Figure S4 | Correlation between average nitrogen phytoplankton concentration of the Eulerian model and the PBM for different mortality-threshold reference volumes.** Correlation for (**a**) $Vref = 1\ L$ (**b**) $10\ L$ and, (**c**) $100\ L$.

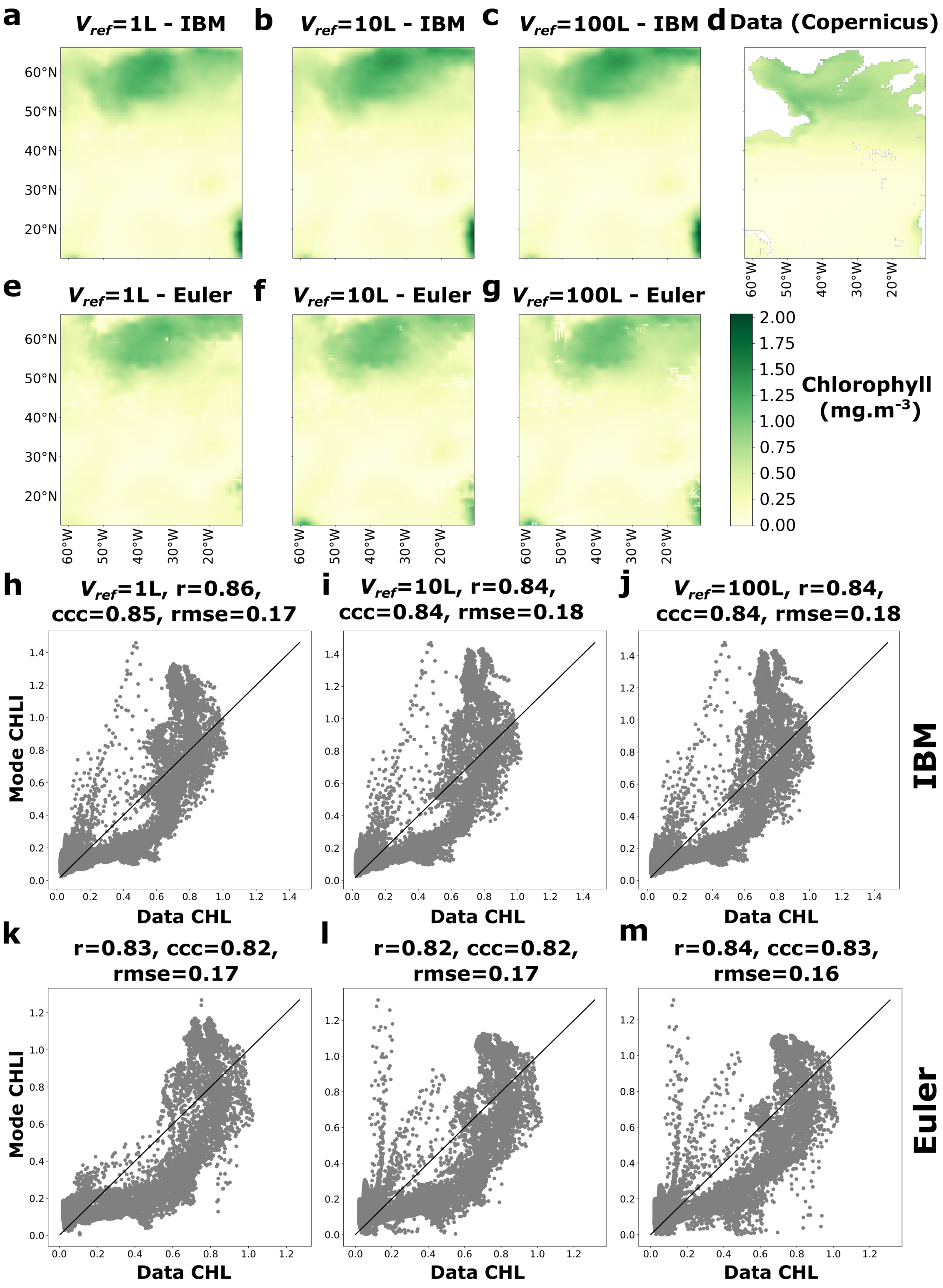


**Figure S5 | Model testing: Particle-Based Model average chlorophyll compared to satellite-derived chlorophyll data (Copernicus).** (**a-c**) Maps of yearly average chlorophyll concentration in the PBM for three different mortality-threshold reference volume. (**d**) Yearly average Copernicus chlorophyll concentrations data derived from satellite measurements (1998-2023 climatology). White area: No data or land. (**d-f**) Correlation between average chlorophyll concentrations of the PBM and Copernicus data for different mortality-threshold reference volumes.

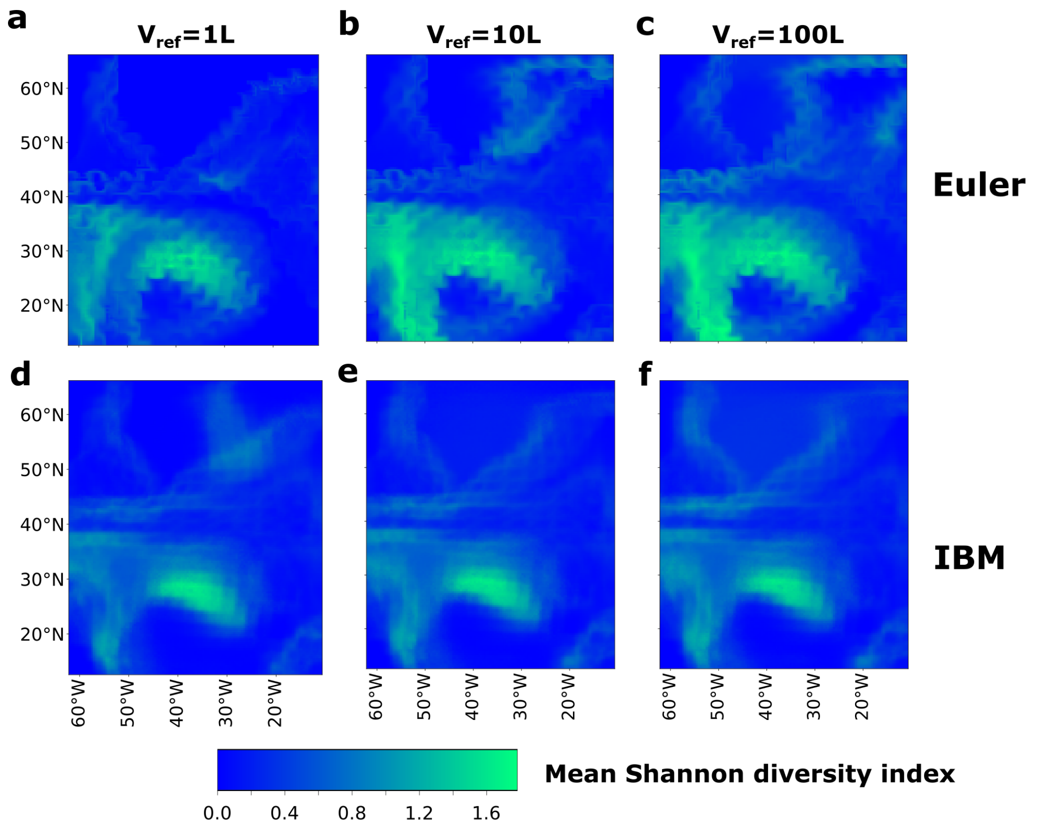


**Figure S6 | Average diversity of the Eulerian and the PBM model for different mortality-threshold reference volumes.** (**a-f**) Average diversity defined as the average Shannon index at each grid point for three different mortality-threshold reference volume for (**a-c**) the Eulerian model and (**d-f**) the PBM model.

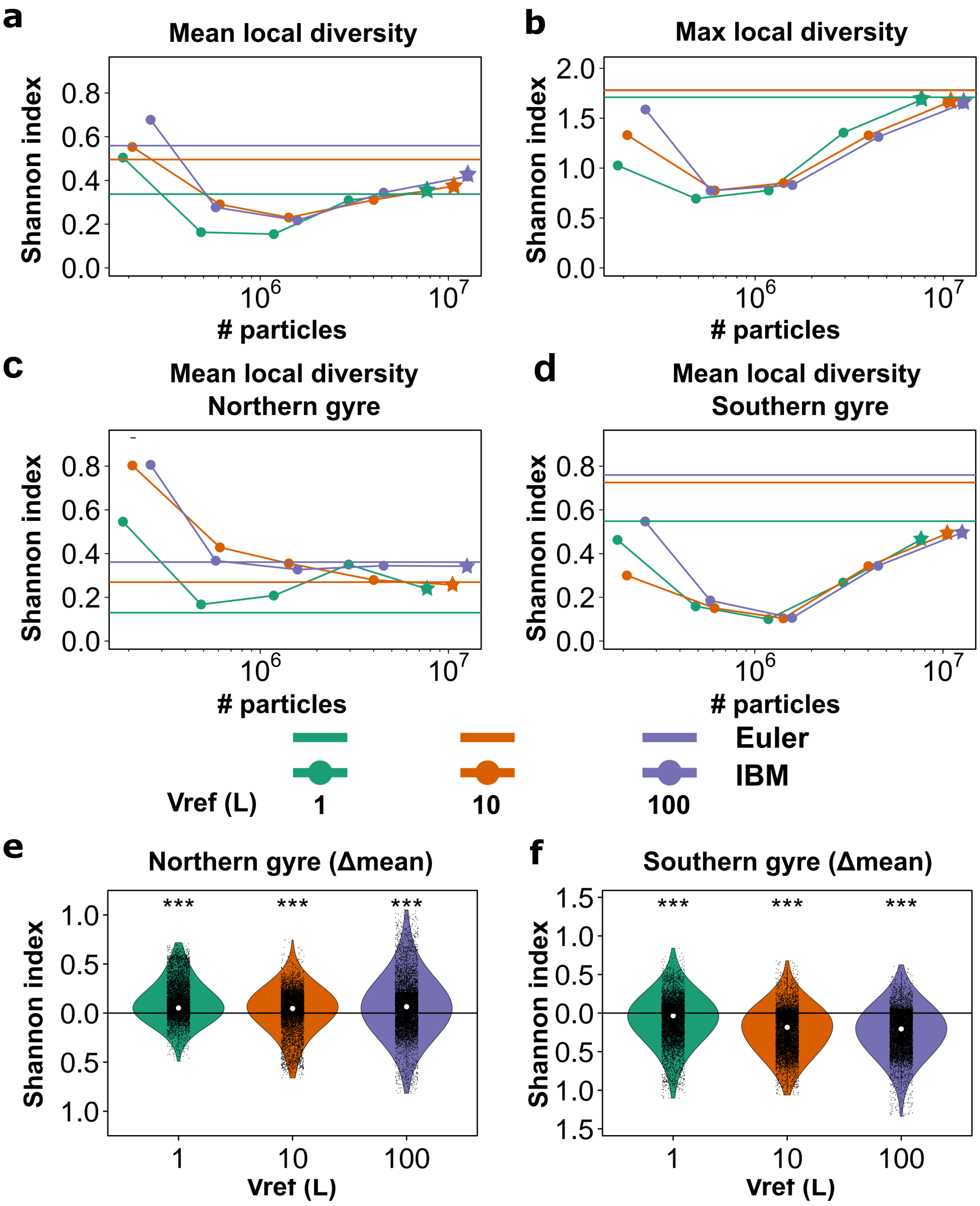


**Figure S7 | Shannon diversity index of the Eulerian and the PBM model for different mortality-threshold reference volumes and comparison of average statistics.** Time-averaged Shannon diversity: (**a**) domain mean, (**b**) maximum, (**c**) northern gyre mean, and (**d**) southern gyre mean, for the Eulerian and PBM models as a function of the average number of PBM particles and the mortality-threshold reference volume. Eulerian data is plotted as horizontal lines for the different mortality-threshold reference volume. PBM simulations presented in panels (**e**) and (**f**) are highlighted with stars for the different mortality-threshold reference volumes. (**e**) Violin plots of the difference in average Shannon diversity index at each grid point between the Eulerian and PBM model for different mortality-threshold reference volume in the northern gyre and (**f**) the southern gyre. Significantly different median delta mean are indicated by stars (Wilcoxon test, $p < 0.001$).

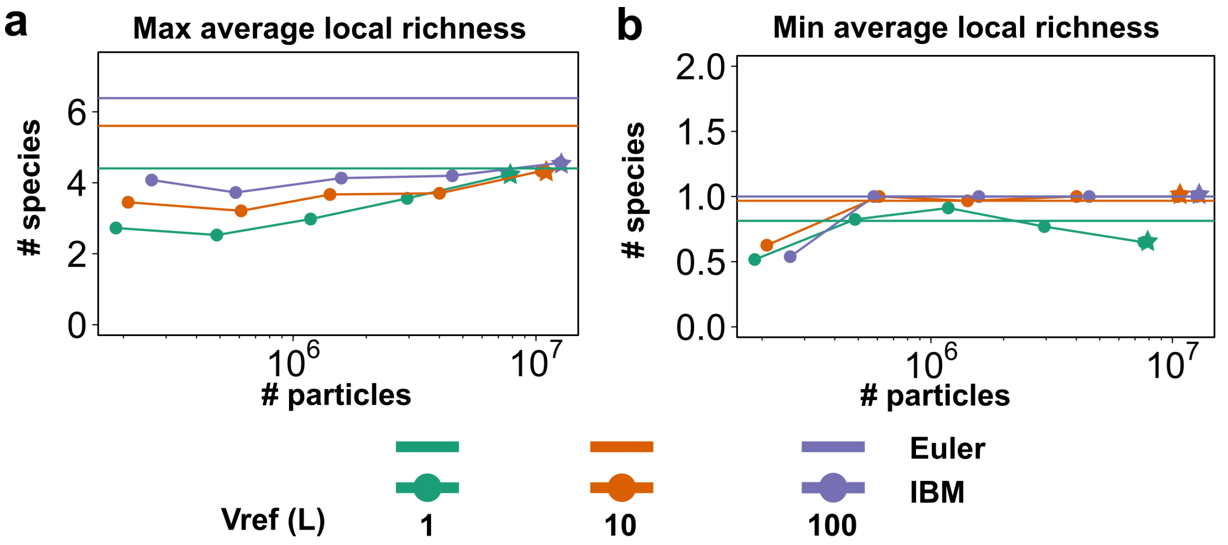


**Figure S8 | Maximum and minimum richness in the PBM model for different mortality-threshold reference volumes.** Time-averaged richness: (**a**) maximum (**b**) minimum for the Eulerian and PBM models as a function of the average number of particles in the PBM and the mortality-threshold reference volume. Eulerian data is plotted as horizontal lines for the different mortality-threshold reference volume. PBM simulations presented in Figure 2 and 3 are highlighted with stars for the different mortality-threshold reference volumes.

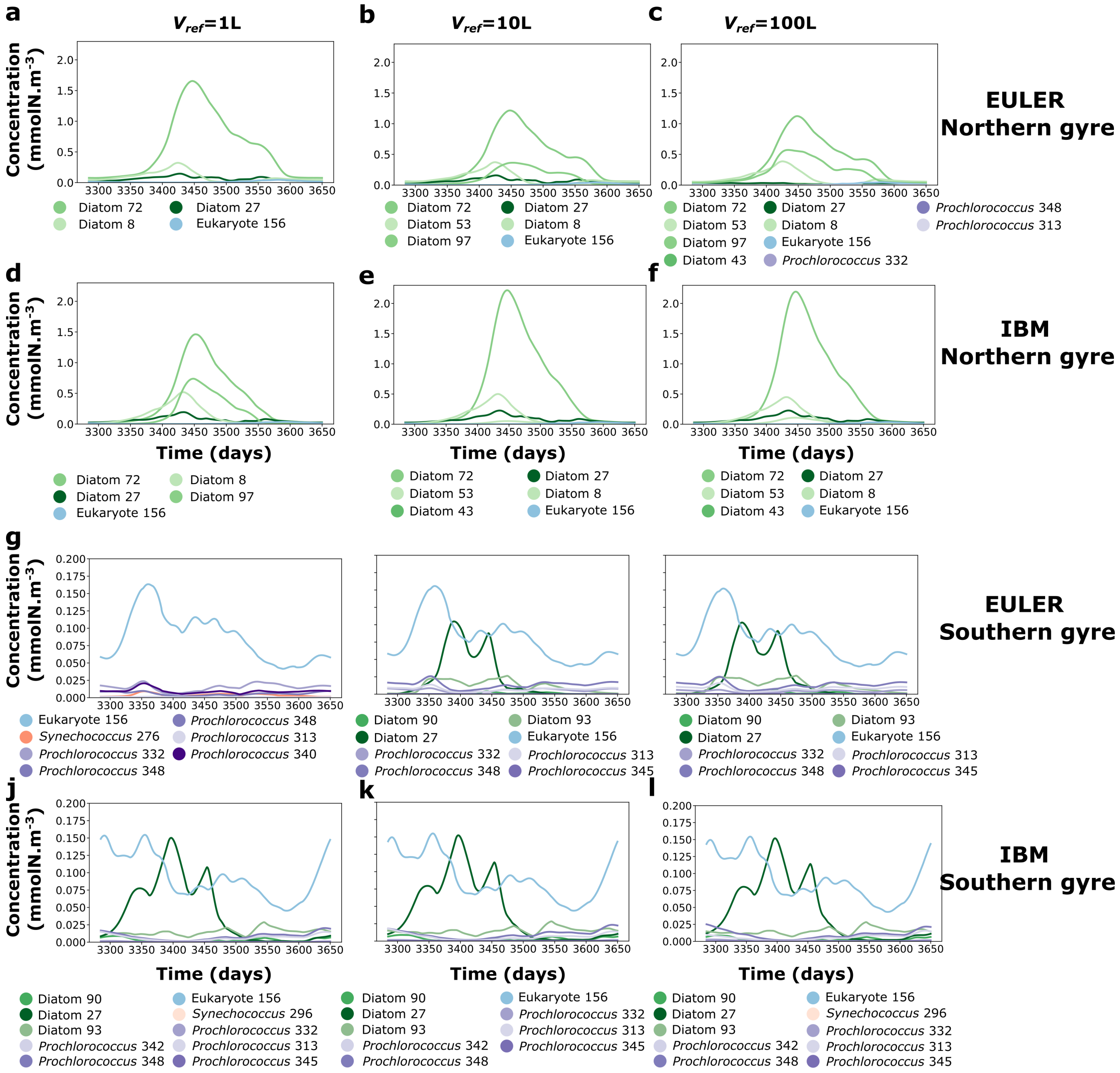


**Figure S9 | Tenth year of simulation time series of average concentration of surviving species in the last year of simulation in the northern and southern gyres for the two different models and different mortality-threshold reference volumes.** Eulerian model in the northern gyre for (**a**) $V_{ref} = 1L$, (**b**) $V_{ref} = 10L$ and (**c**) $V_{ref} = 100L$. PBM model in the northern gyre for (**d**) for $V_{ref} = 1L$, (**e**) $V_{ref} = 10L$ and (**f**) $V_{ref} = 100L$. Eulerian model in the southern gyre for (**g**) $V_{ref} = 1L$, (**h**) $V_{ref} = 10L$ and (**i**) $V_{ref} = 100L$. PBM model in the southern gyre for (**j**) $V_{ref} = 1L$, (**k**) for $V_{ref} = 10L$ and (**l**) $V_{ref} = 100L$. Diatoms are in shades of green, eukaryotes (non-diatom) in shades of blue, *Synechococcus* (excluded) in shades of red and *Prochlorococcus* in shades of purple.

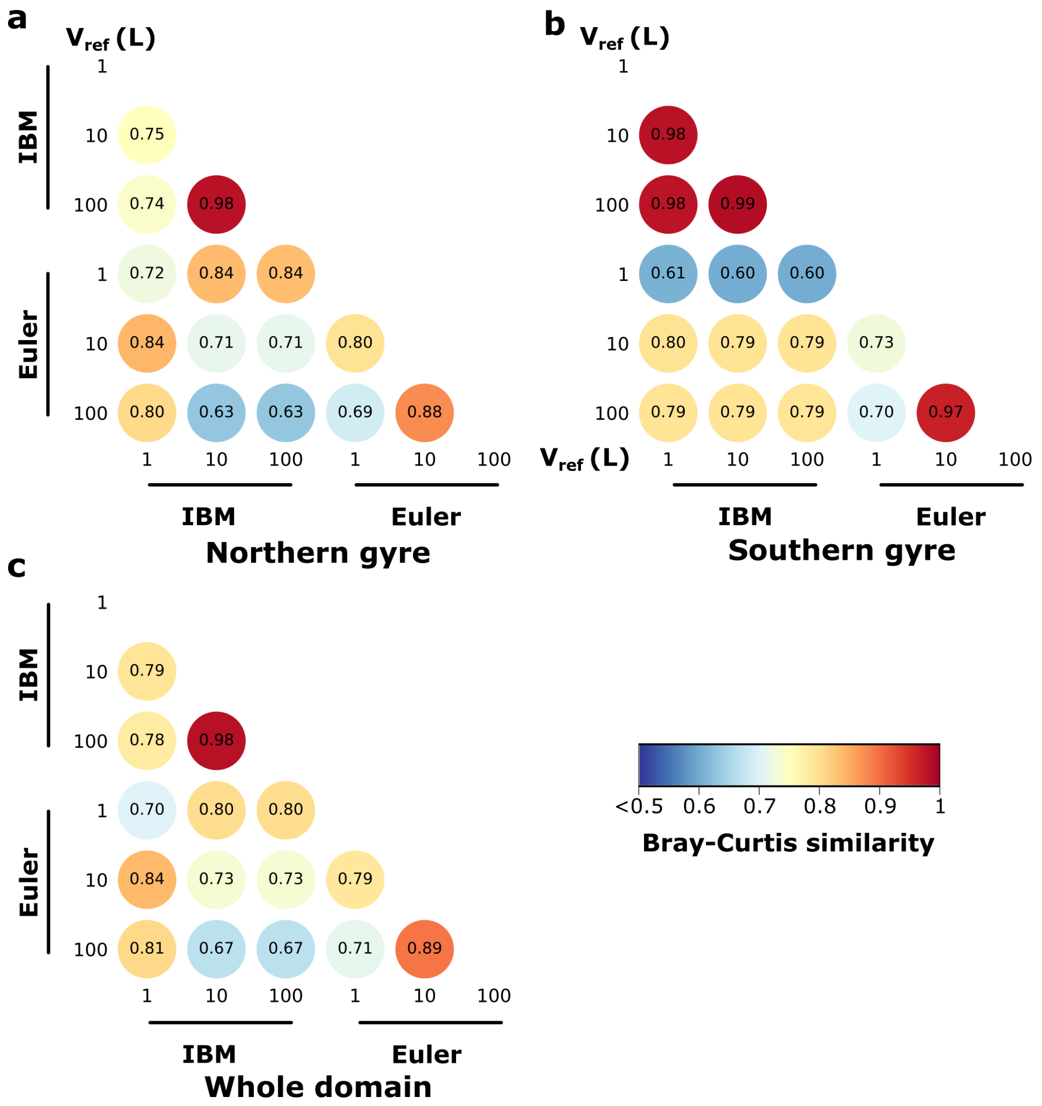


**Figure S10 | Bray-Curtis similarity indices comparing time series of the last year of simulation across mortality-threshold and reference volumes.** (**a**- **c**) Pairwise Bray-Curtis similarity indices between time series (all species taken into account) of the last year of simulation for (**a**) the northern gyre, (**b**) the southern gyre and (**c**) the whole domain.

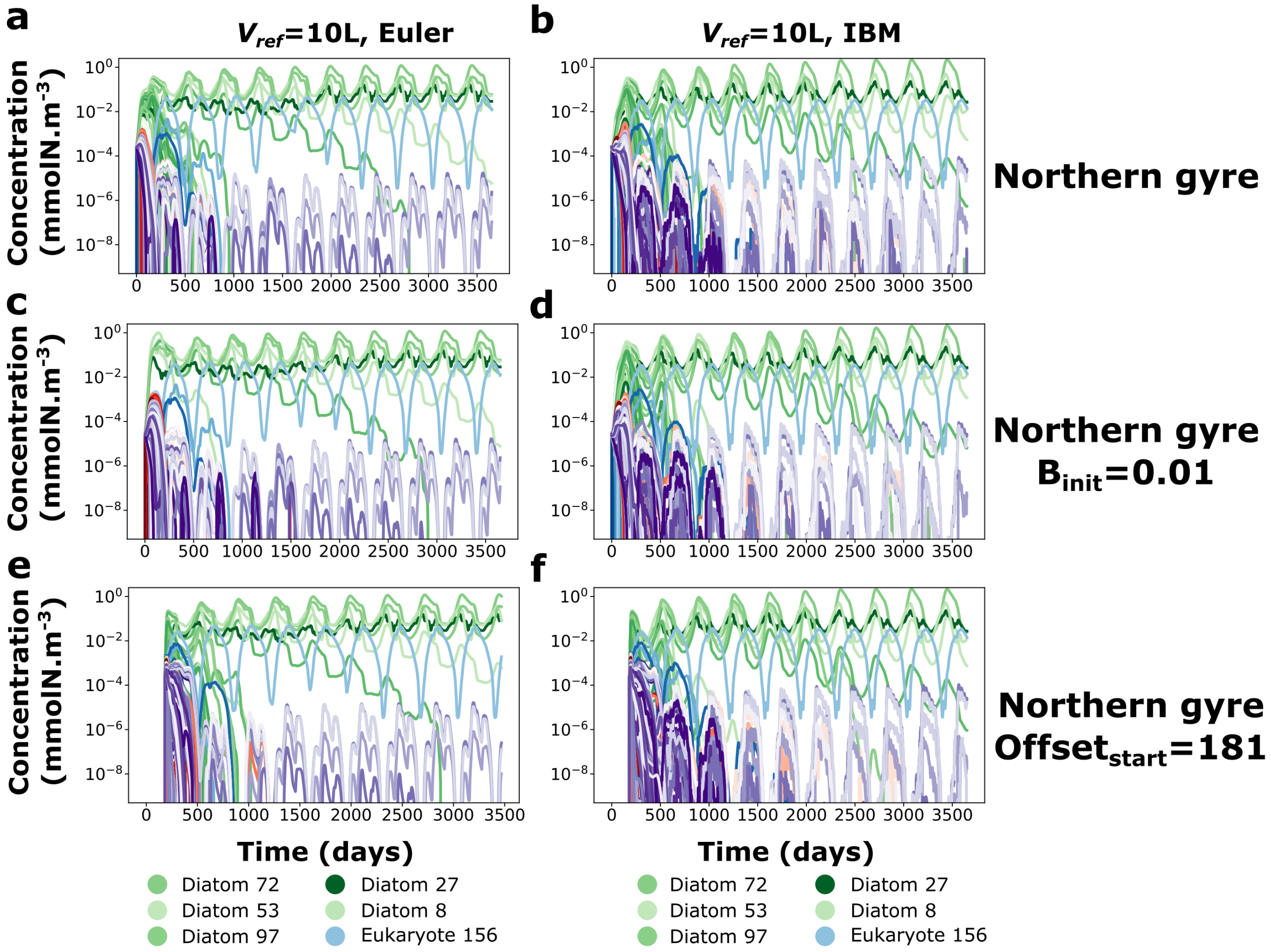


**Figure S11 | Sensitivity analysis: time series of average species dynamics for different initial conditions in the northern gyre.** (**a**, **b**) Eulerian and PBM with default initial conditions; (**c**, **d**) with lower initial biomass; and (**e**, **f**) with simulations initiated in July. All panels show the simulation with $V_{ref} = 100\ L$. Diatoms are in shades of green, eukaryotes (non-diatom) in shades of blue, *Synechococcus* (excluded) in shades of red and *Prochlorococcus* in shades of purple. For clarity, the color legend includes only species that survive in the last year of simulation throughout the whole gyre and year.

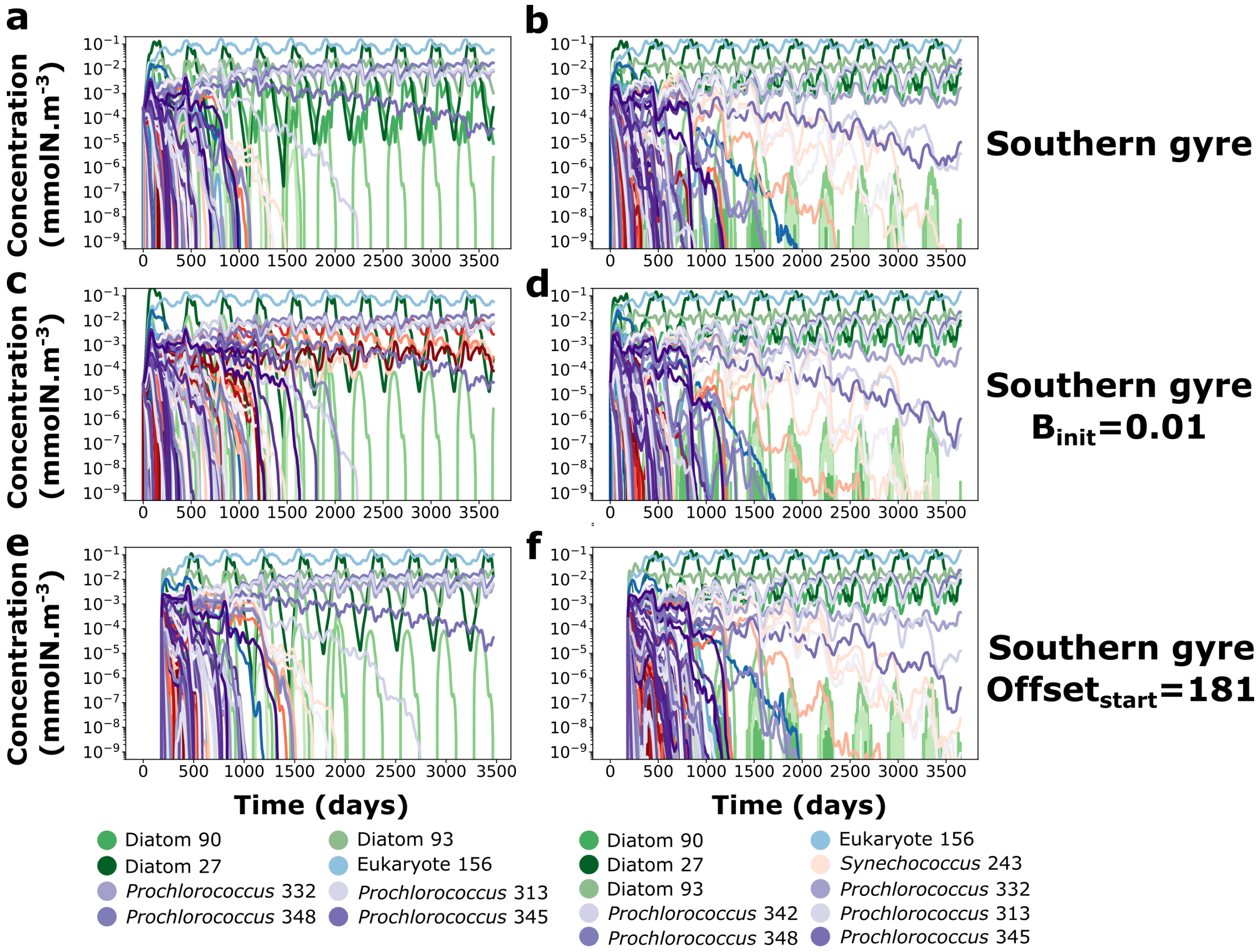


**Figure S12 | Sensitivity analysis: time series of average species dynamics for different initial conditions in the southern gyre.** (**a**, **b**) Eulerian and PBM with default initial conditions; (**c**, **d**) with lower initial biomass; and (**e**, **f**) with simulations initiated in July. All panels show the simulation with $V_{ref} = 100\ L$. Diatoms are in shades of green, eukaryotes (non-diatom) in shades of blue, *Synechococcus* in shades of red and *Prochlorococcus* in shades of purple. For clarity, the color legend includes only species that survive in the last year of simulation throughout the whole gyre, and year.

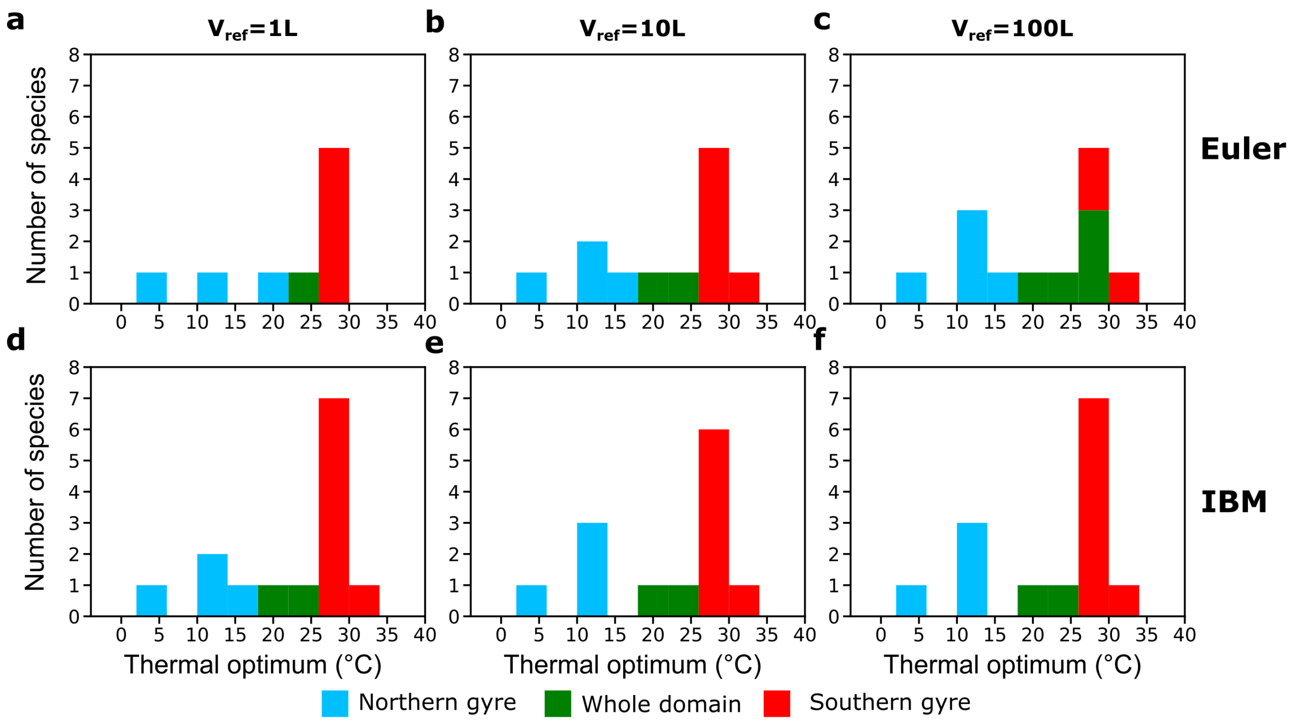


**Figure S13 | Distributions and biogeographical classification of thermal optima of surviving species across mortality-threshold and reference volumes.** (**a**- **c**) Eulerian model, (**d-e**) PBM. Species are colored according to their biogeographical range. For a species to be considered present in a gyre, it has to be present throughout the whole year.

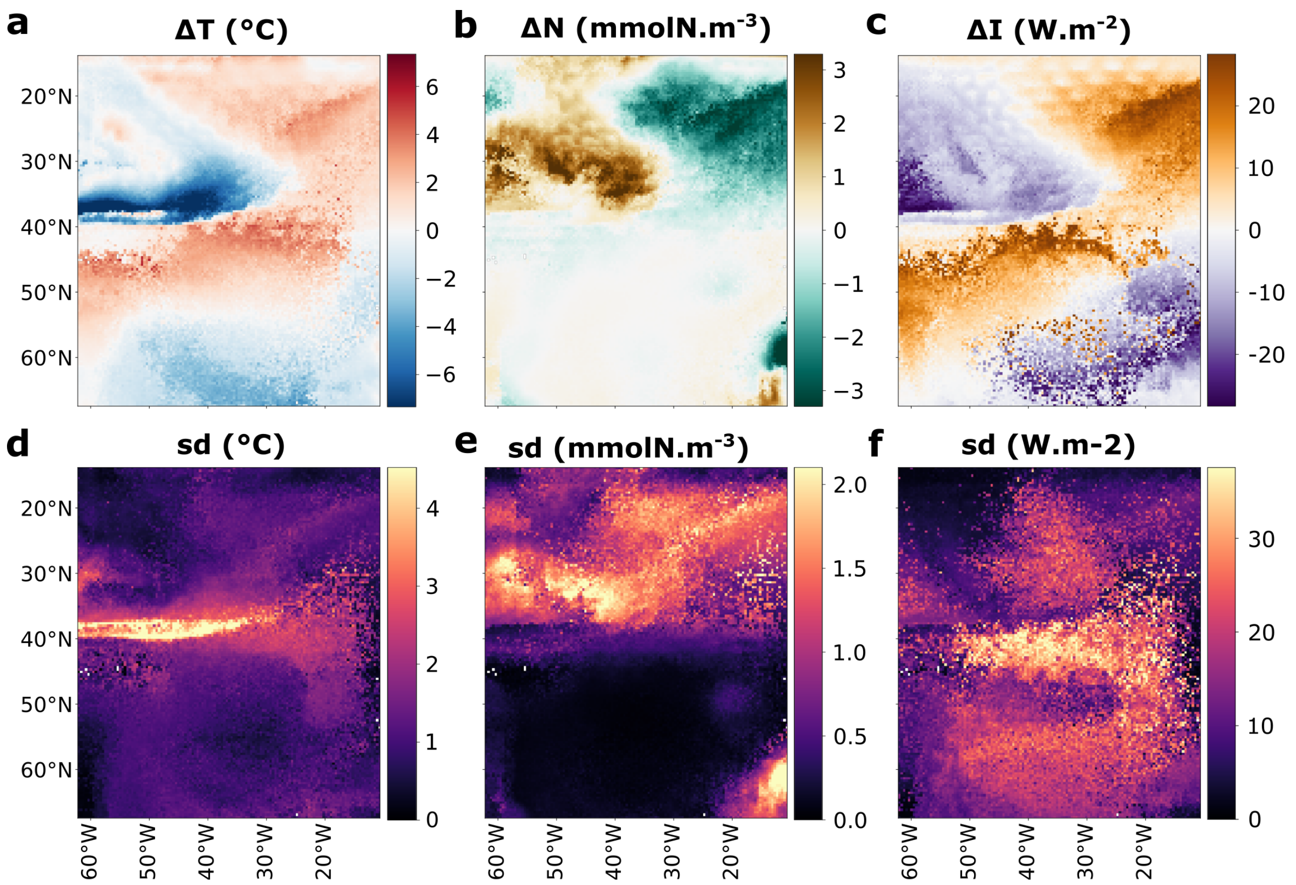


**Figure S14 | Differences between Lagrangian environmental histories and fixed-location Eulerian histories at particle death locations.** (**a**–**c**) Trajectory-length-weighted median differences between Lagrangian environmental histories and fixed-location Eulerian histories, evaluated at the grid point where each particle dies, for (**a**) temperature, (**b**) nitrate, and (**c**) light. (**d**–**f**) Trajectory-length-weighted standard deviations of the corresponding differences at each grid point.

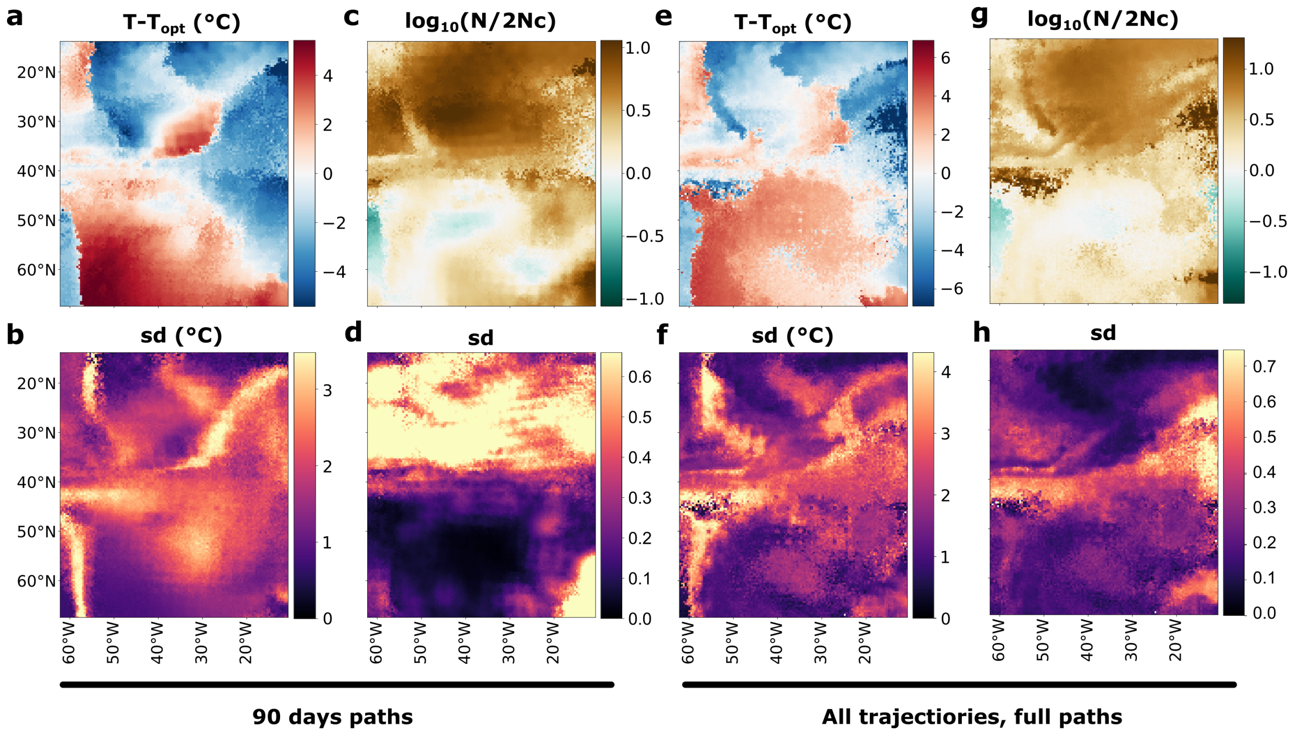


**Figure S15 | Differences between Lagrangian environmental histories and species' environmental optima.** (**a**–**d**) Differences calculated over 90-day paths and evaluated at each path's endpoint: median differences for (**a**) temperature and (**c**) nitrate, and standard deviations of the corresponding differences for (**b**) temperature and (**d**) nitrate. (**e**–**h**) Trajectory-length-weighted differences evaluated at each particle's death location: median differences for (**e**) temperature and (**g**) nitrate, and trajectory-length-weighted standard deviations for (**f**) temperature and (**h**) nitrate. All differences are calculated relative to species' respective environmental optima.

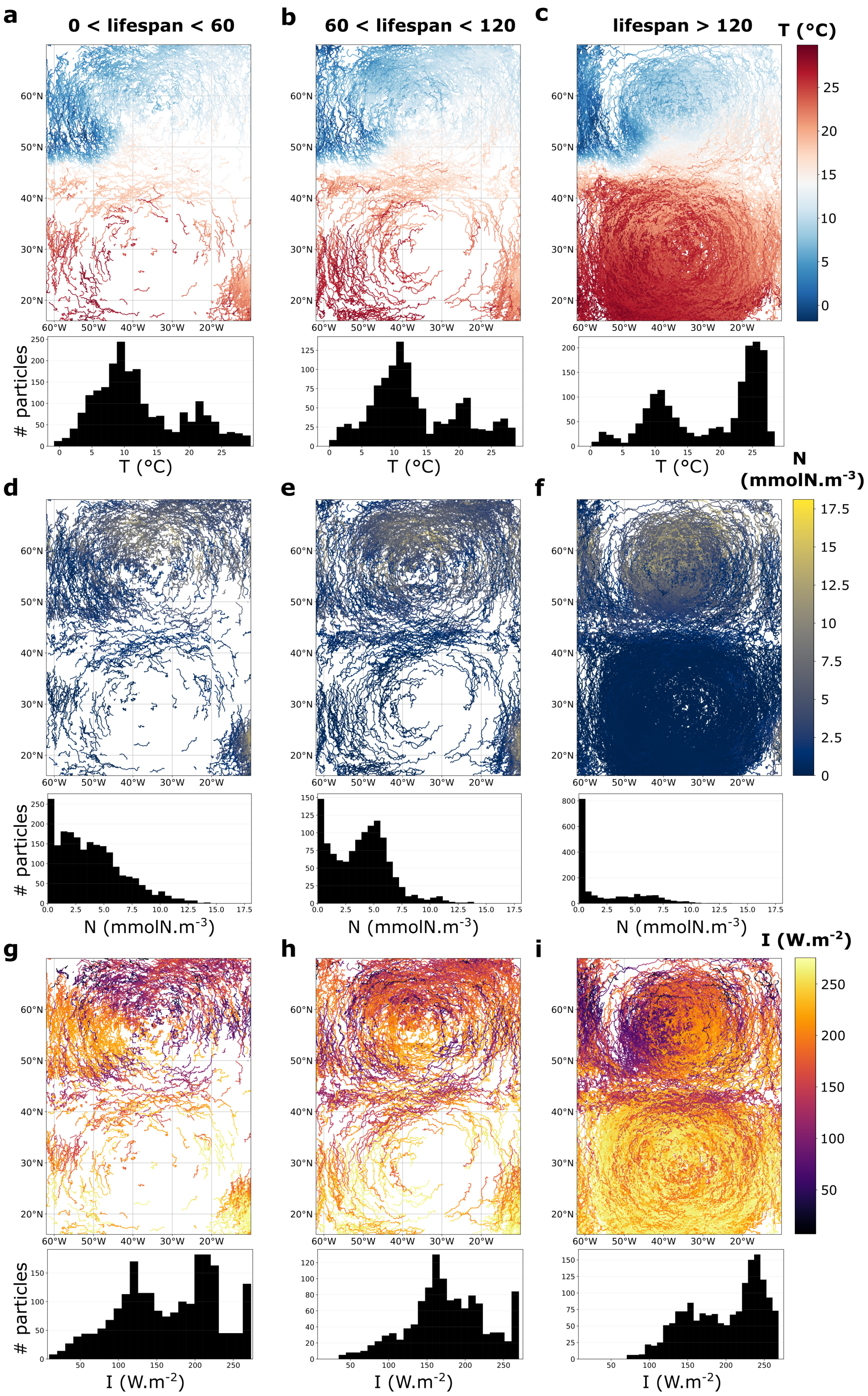


**Figure S16 | Lagrangian histories and trajectories as a function of particle lifespan.** Temperature, nitrate, and light trajectories and distributions of their mean values for particles with lifespans of (**a**, **d**, **g**) 0–60 days, (**b**, **e**, **h**) 60–120 days, and (**c**, **f**, **i**) more than 120 days. All plots shown are for the simulation with $V_{ref} = 1L$.

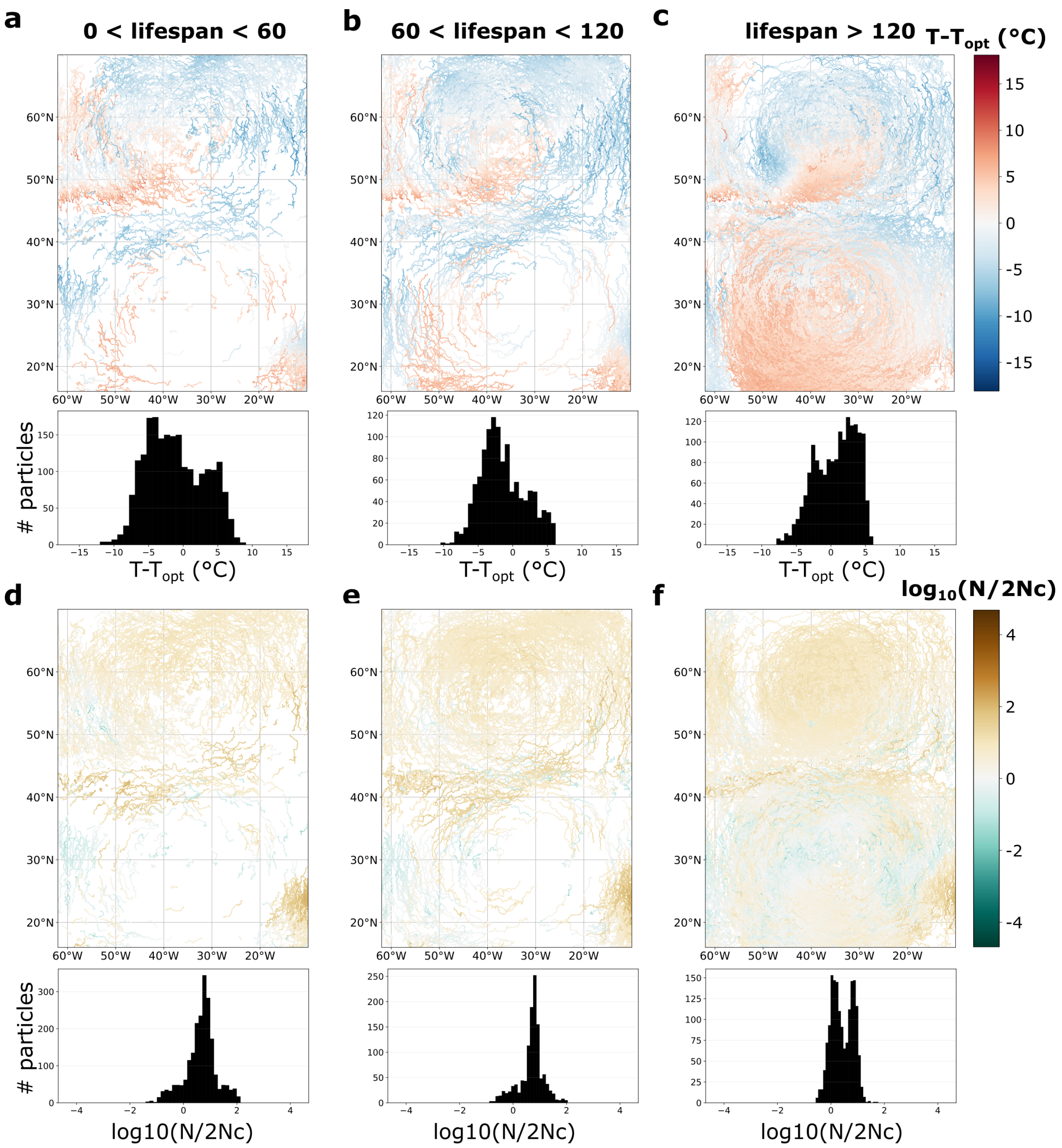


**Figure S17 | Differences between Lagrangian environmental histories and species' optima along theirs trajectories as a function of particle lifespan.** Differences between local temperature/nitrate and species optimum and distributions of their mean values for particles with lifespans of (**a**, **d**, **g**) 0–60 days, (**b**, **e**, **h**) 60–120 days, and (**c**, **f**, **i**) more than 120 days. All plots shown are for the simulation with $V_{ref} = 1L$.

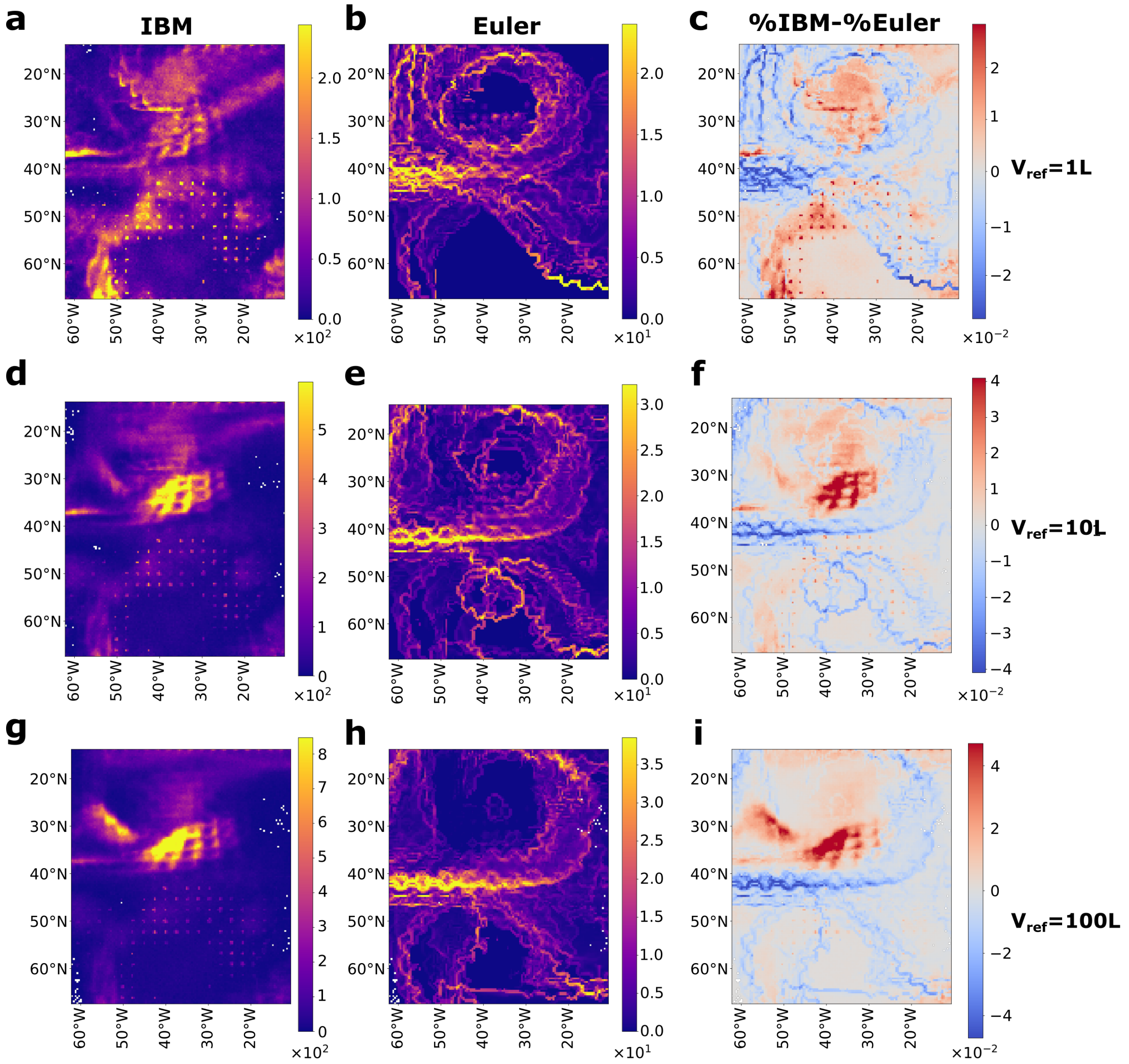


**Figure S18 | Spatial patterns of Lagrangian and Eulerian mortality.** (**a**, **d**, **g**) Number of Lagrangian particle deaths at each grid point. (**b**, **e**, **h**) Number of Eulerian deaths at each grid point. (**c**, **f**, **i**) Difference between the percentages of Lagrangian and Eulerian deaths at each grid point, calculated as Lagrangian minus Eulerian.

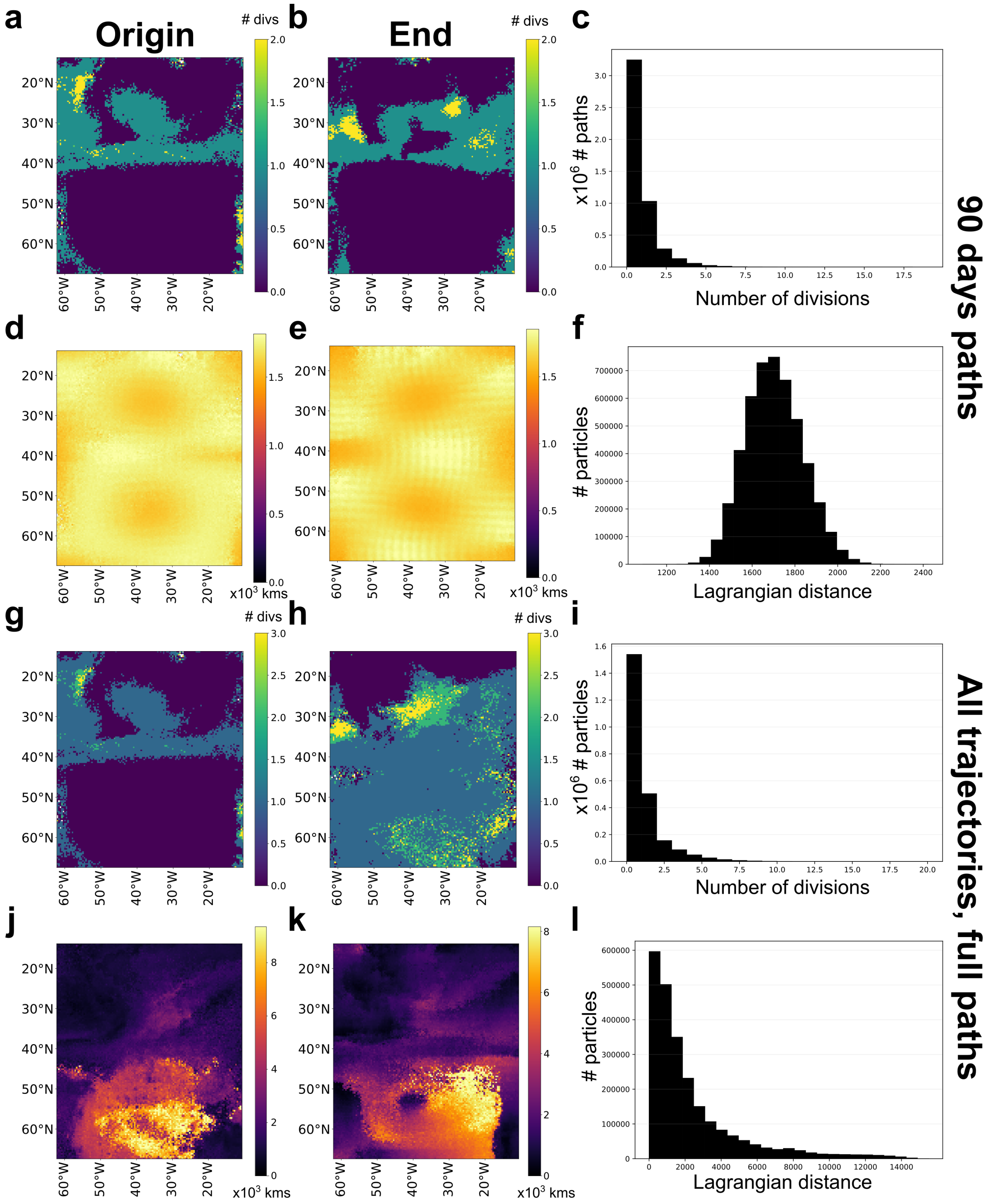


**Figure S19 | Particle divisions and Lagrangian path distances at path origins and endpoints.** (**a**–**c**) Number of particle divisions over 90-day paths: median number of divisions at (**a**) path origins and (**b**) path endpoints, and distribution of the number of divisions across paths (**c**). (**d**–**f**) Lagrangian path distances over 90-day paths: median distances from (**d**) path origins and (**e**) path endpoints, and distribution of path distances (**f**). (**g**–**l**) Corresponding results calculated over full Lagrangian trajectories.

**Videos 1 | Total phytoplankton biomass dynamics in PBM and Eulerian models across varying mortality thresholds (last year of simulation)** Eulerian model for (**a**) $V_{ref} = 1$, (**b**) 10 and, (**c**) 100 $L$. PBM for (**d**) $V_{ref} = 1$, (**e**) 10 and, (**f**) 100 $L$.

**Videos 2 | Zooplankton biomass dynamics in PBM and Eulerian models across varying mortality thresholds (last year of simulation)** Eulerian model for (**a**) $V_{ref} = 1$, (**b**) 10 and, (**c**) 100 $L$. PBM for (**d**) $V_{ref} = 1$, (**e**) 10 and, (**f**) 100 $L$.

**Videos 3 | Richness dynamics PBM and Eulerian models across varying mortality thresholds (last year of simulation)** Eulerian model for (**a**) $V_{ref} = 1$, (**b**) 10 and, (**c**) 100 $L$. PBM for (**d**) $V_{ref} = 1$, (**e**) 10 and, (**f**) 100 $L$.

**Videos 4 | Shannon diversity index dynamics in PBM and Eulerian models across varying mortality thresholds (last year of simulation)** Eulerian model for (**a**) $V_{ref} = 1$, (**b**) 10 and, (**c**) 100 $L$. PBM for (**d**) $V_{ref} = 1$, (**e**) 10 and, (**f**) 100 $L$.

**Videos 5 | Dominant species dynamics in PBM and Eulerian models across varying mortality thresholds (last year of simulation).** Eulerian model for (**a**) $V_{ref} = 1$, (**b**) 10 and, (**c**) 100 $L$. PBM for (**d**) $V_{ref} = 1$, (**e**) 10 and, (**f**) 100 $L$.
