## Supplementary figures and images for "Particle-based representation of phytoplankton modulates emergent coexistence and biogeography in a model ocean"

### Video_1a_phyto_biomass_euler_vref1.gif

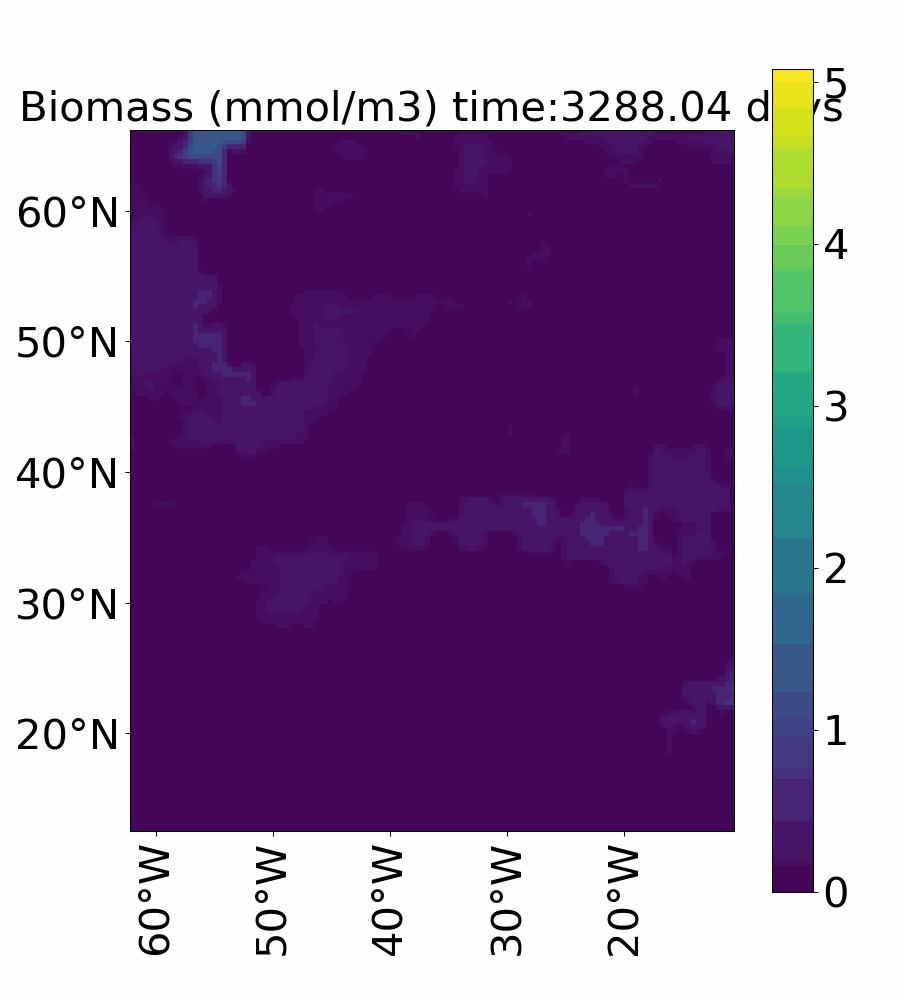

### Video_1b_phyto_biomass_euler_vref10.gif

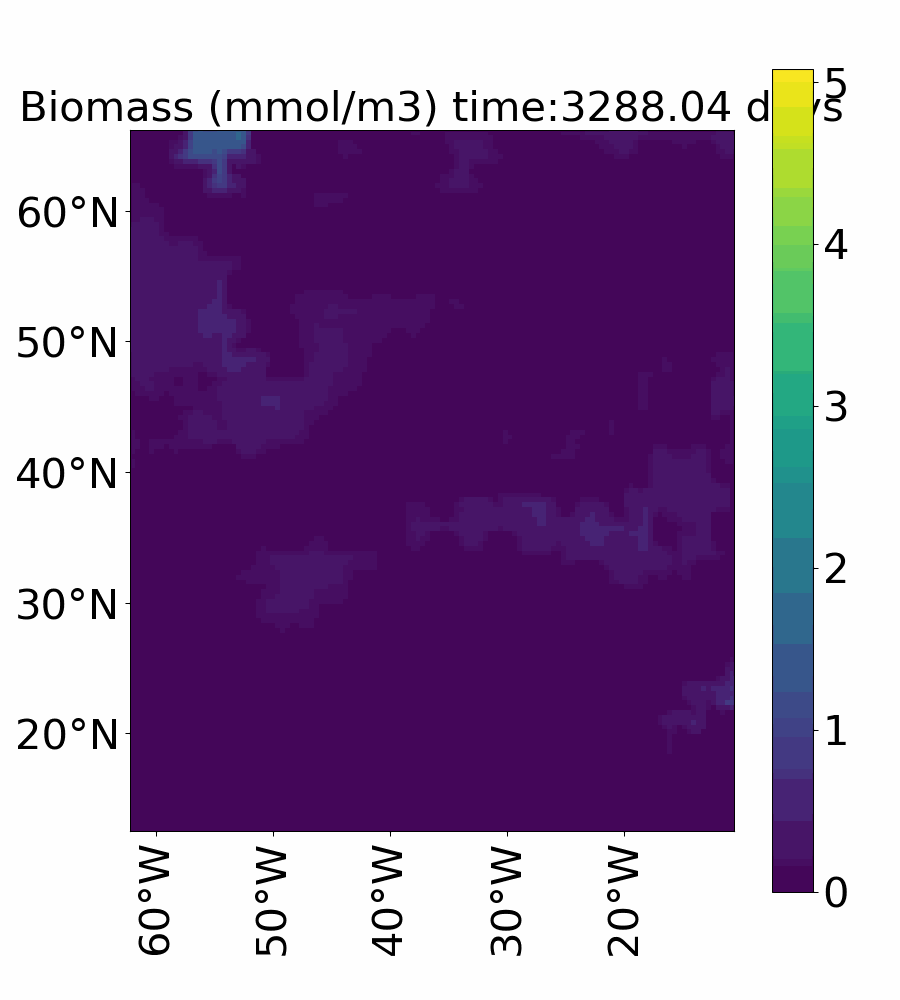

### Video_1c_phyto_biomass_euler_vref100.gif

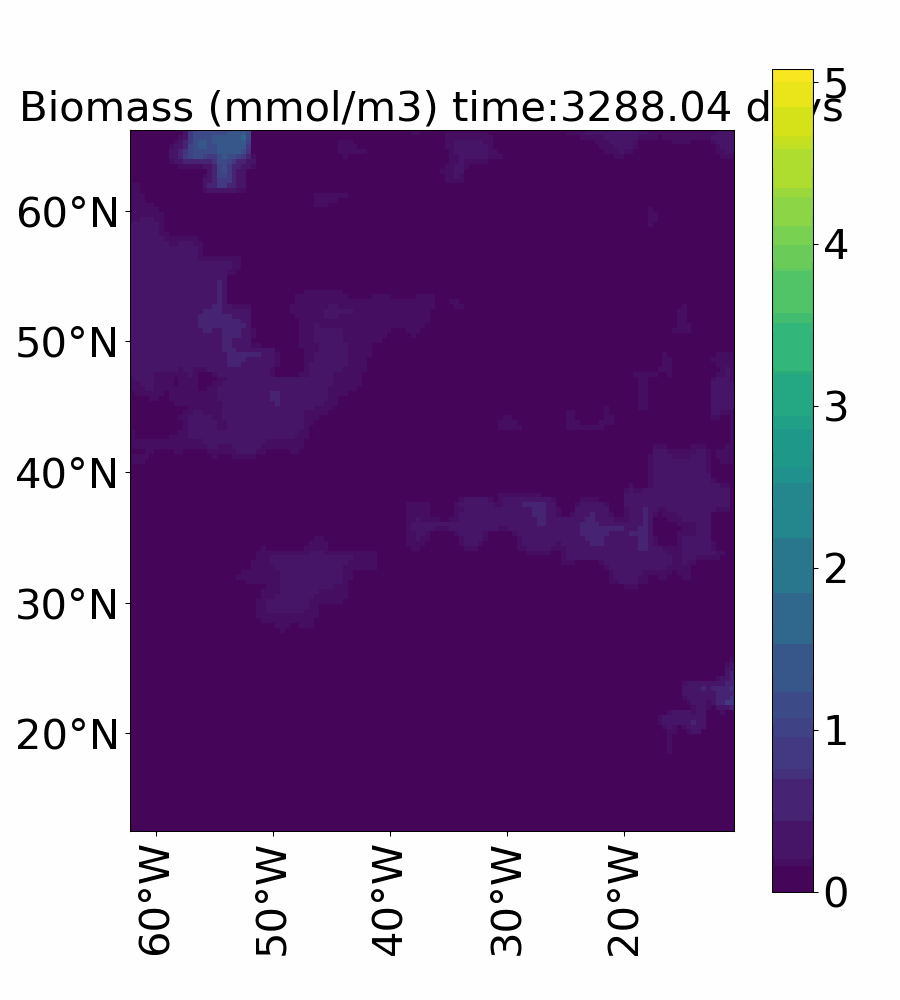

### Video_1d_phyto_biomass_ibm_vref1.gif

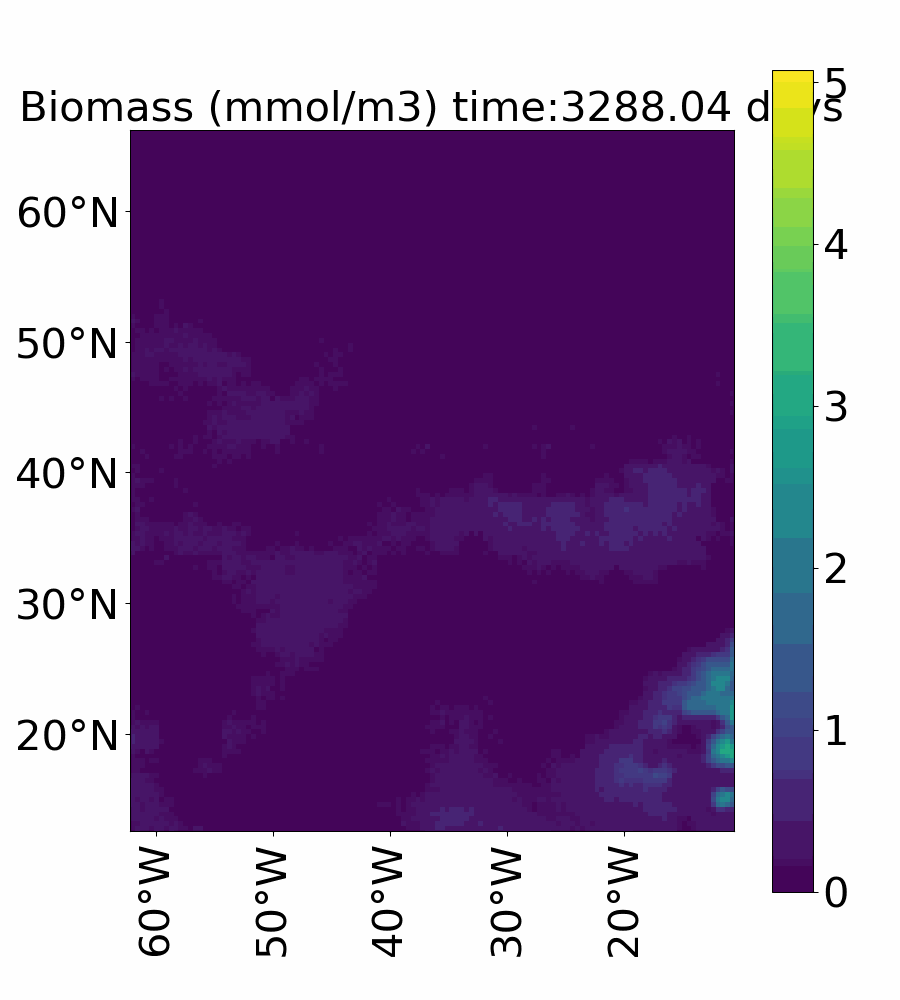

### Video_1e_phyto_biomass_ibm_vref10.gif

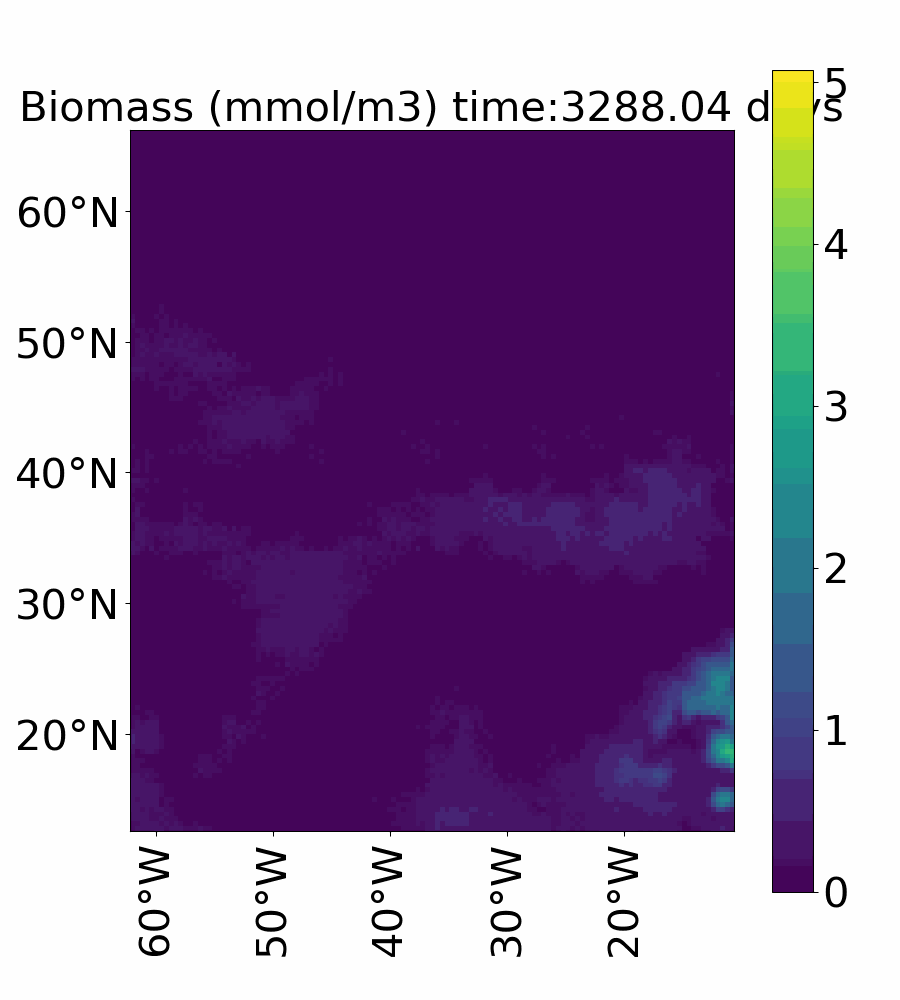

### Video_1f_phyto_biomass_ibm_vref100.gif

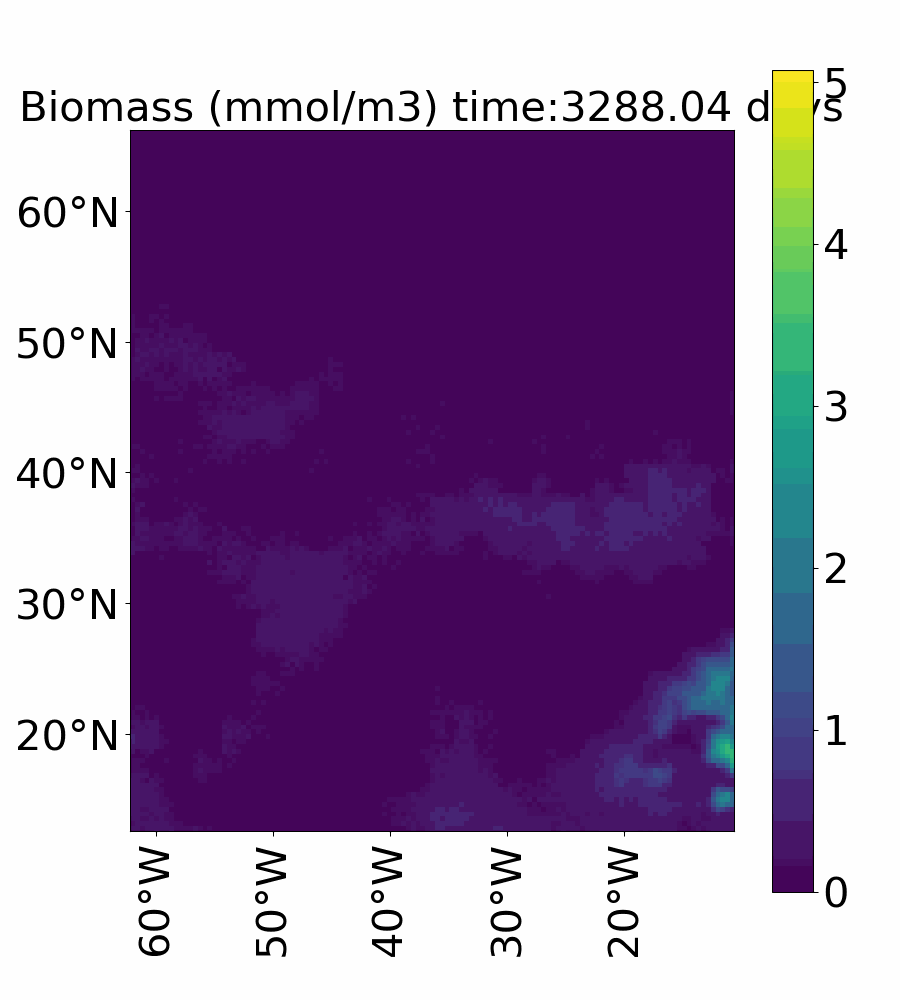

### Video_2a_zoop_biomass_euler_vref1.gif

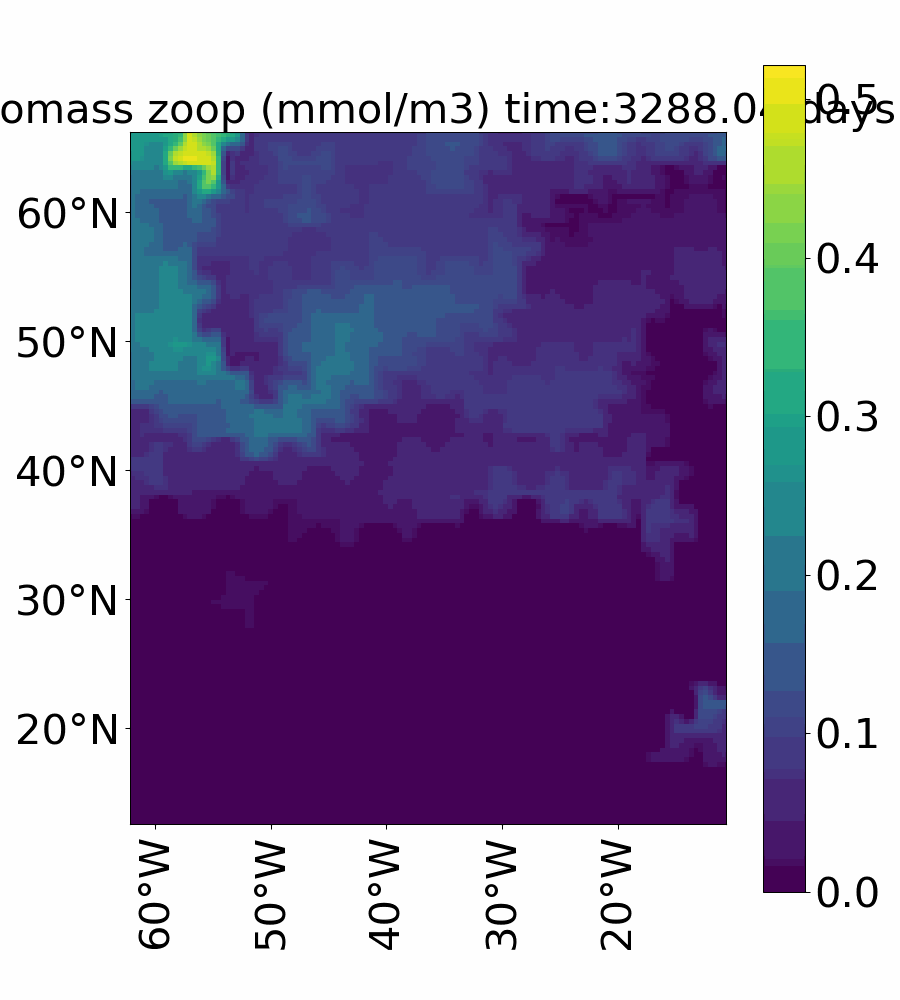

### Video_2b_zoop_biomass_euler_vref10.gif

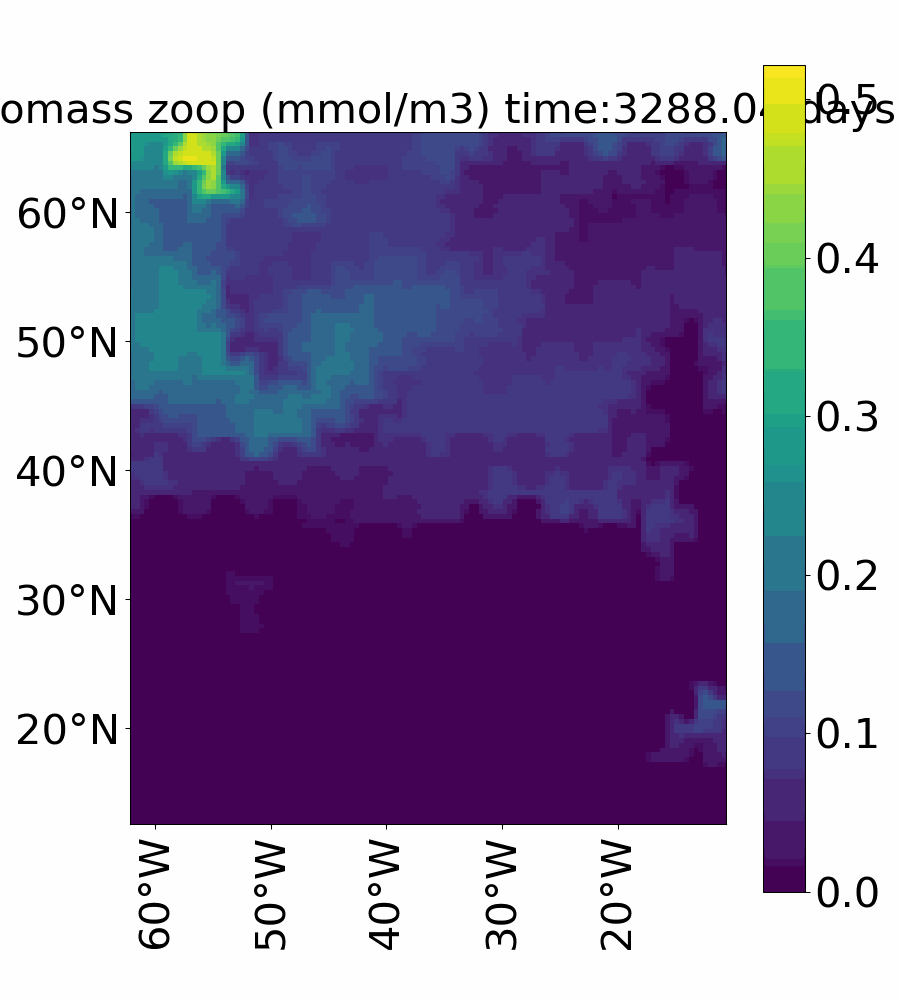

### Video_2c_zoop_biomass_euler_vref100.gif

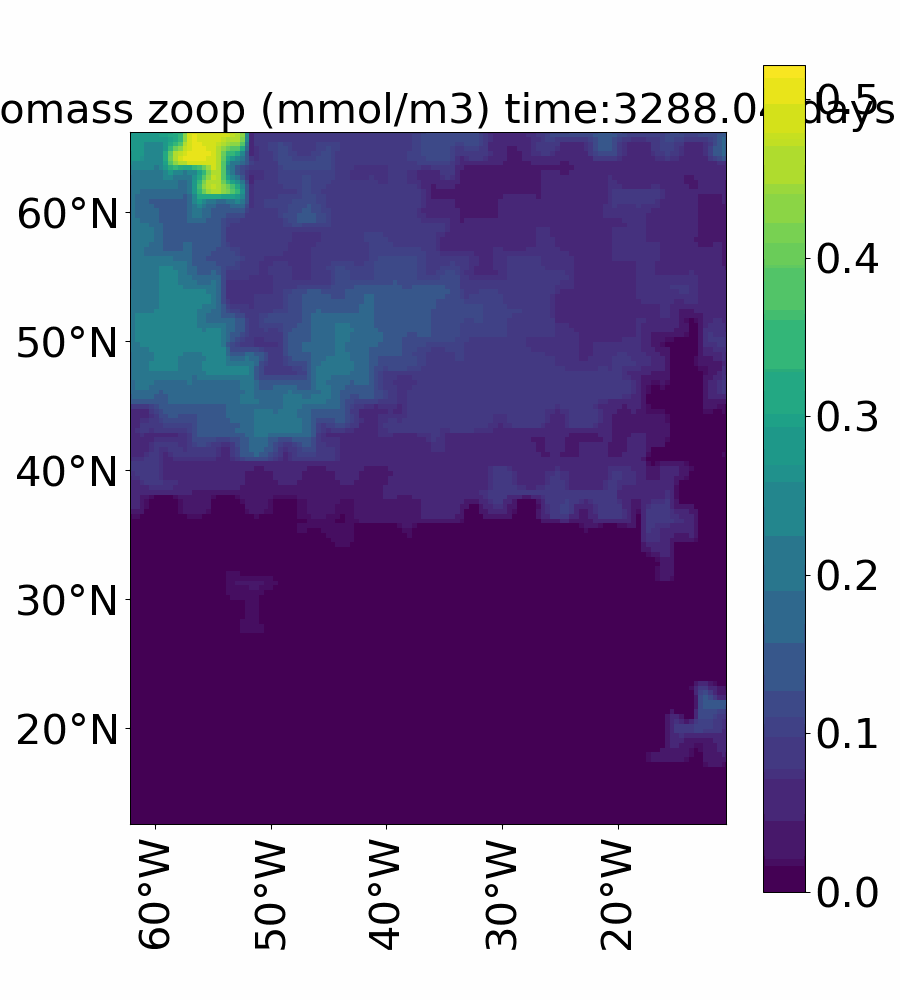

### Video_2d_zoop_biomass_ibm_vref1.gif

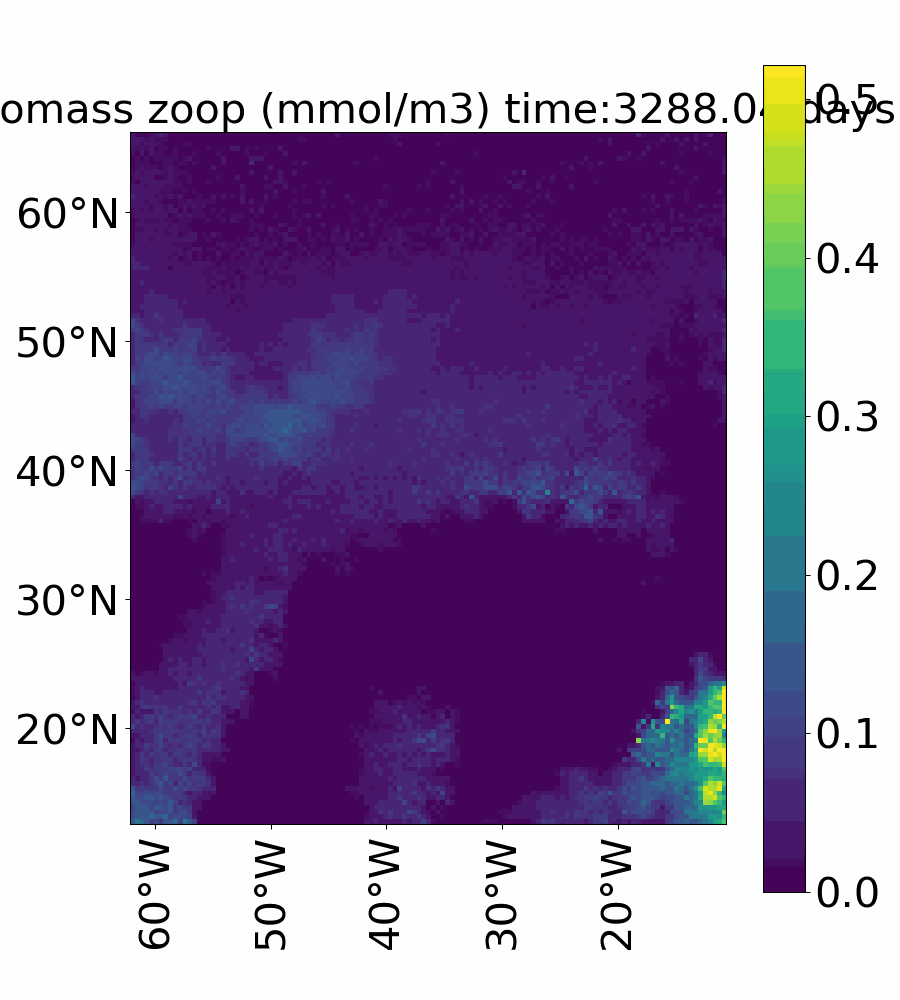

### Video_2e_zoop_biomass_ibm_vref10.gif

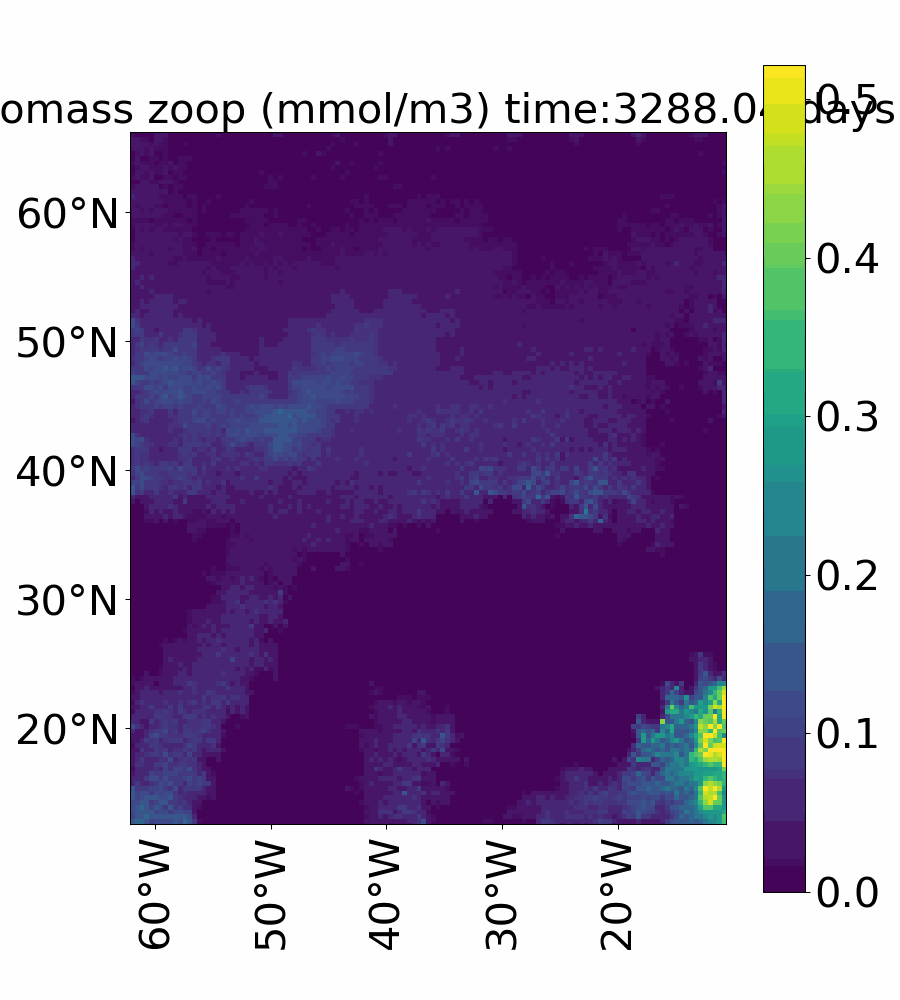

### Video_2f_zoop_biomass_ibm_vref100.gif

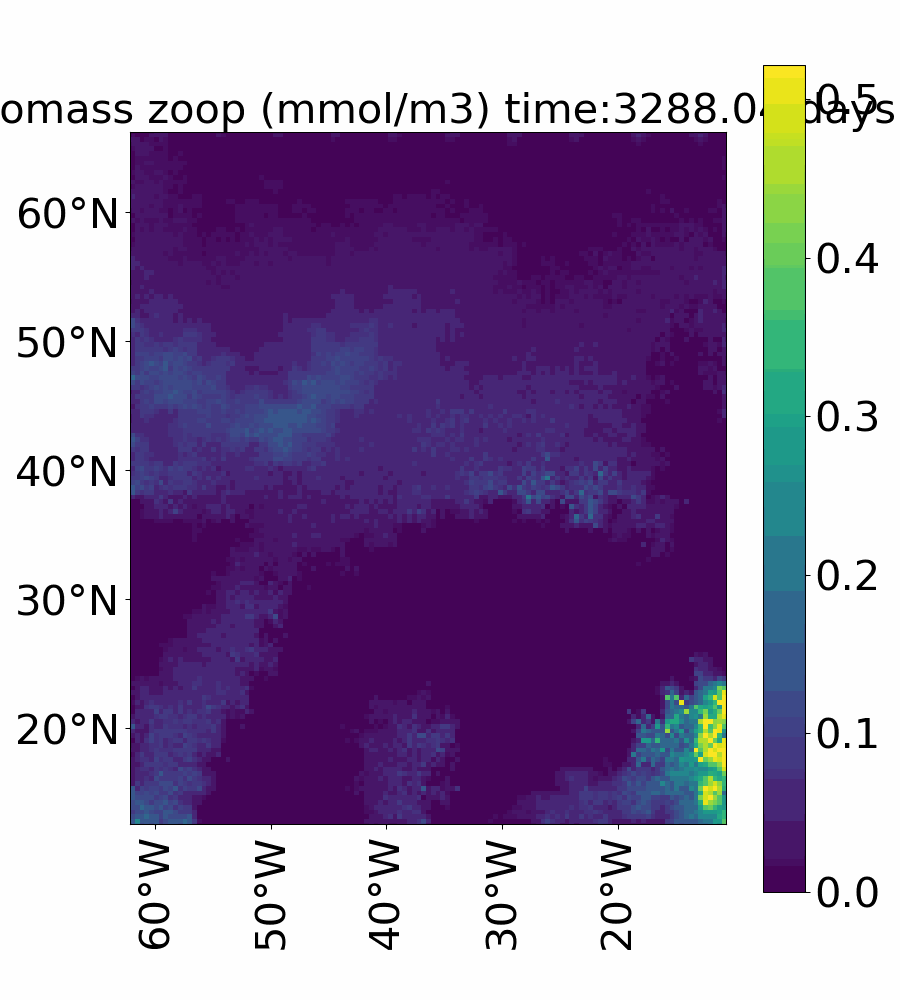

### Video_3a_richness_euler_vref1.gif

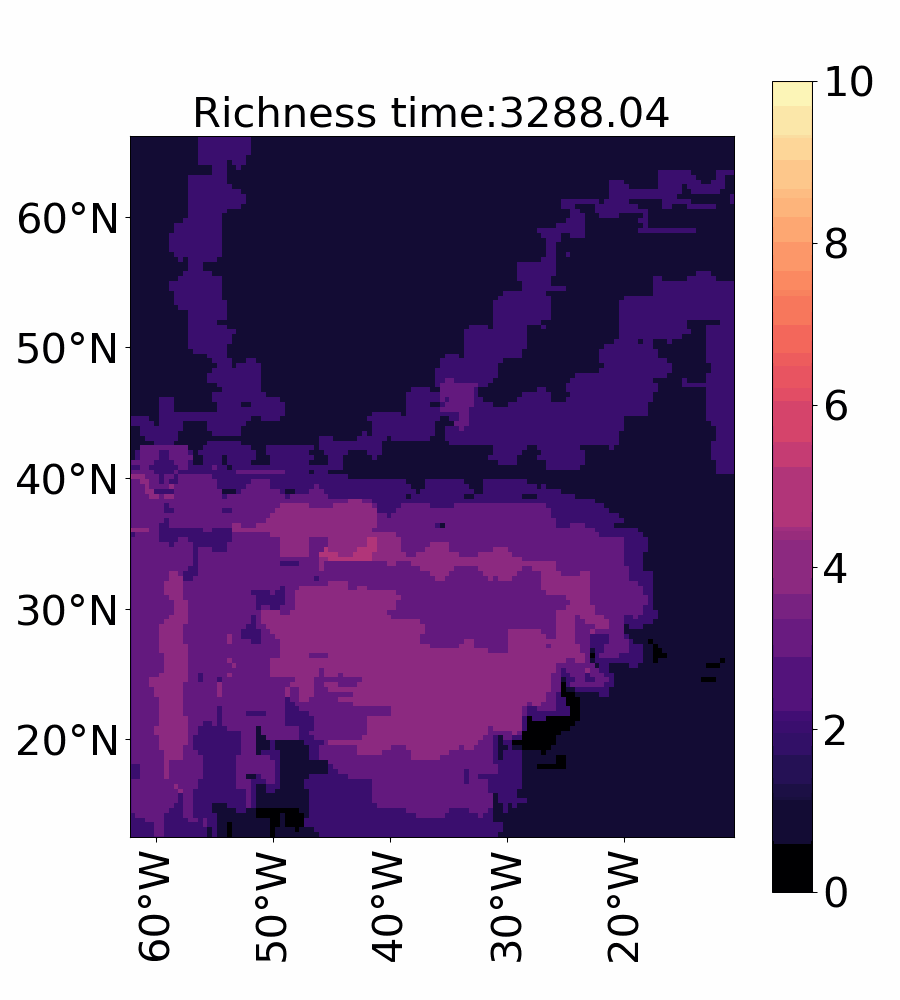

### Video_3b_richness_euler_vref10.gif

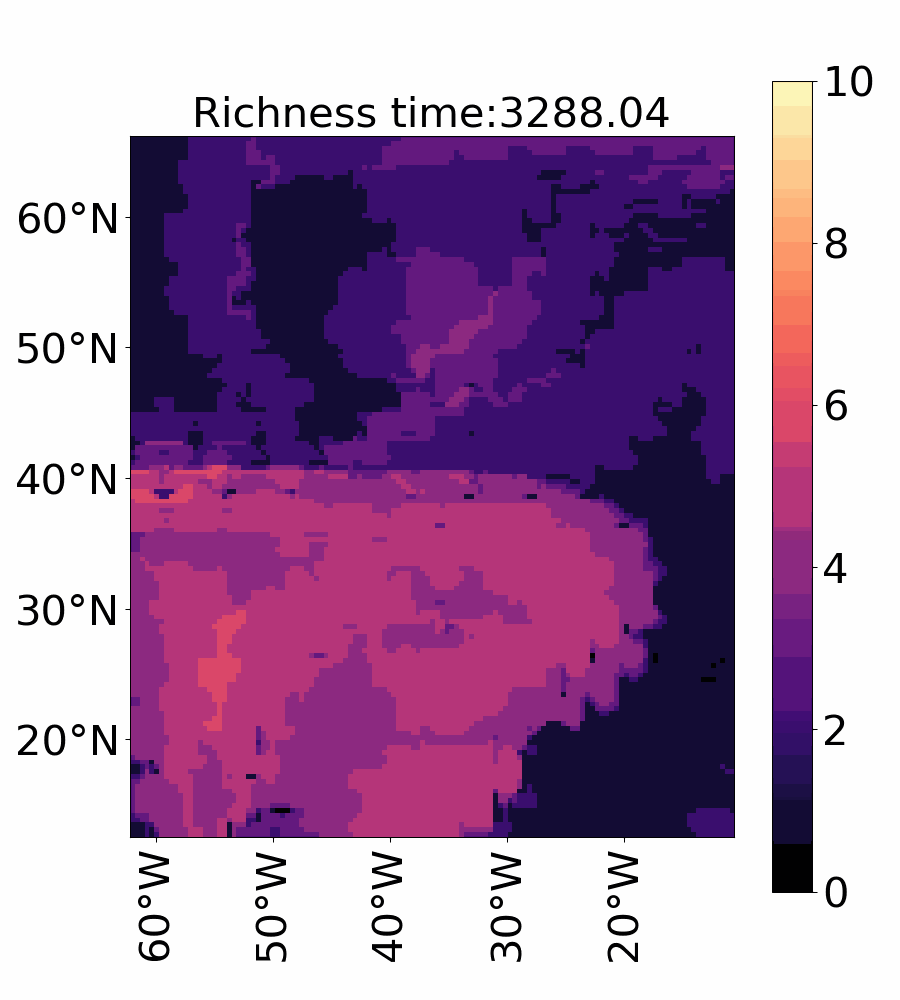

### Video_3c_richnesss_euler_vref100.gif

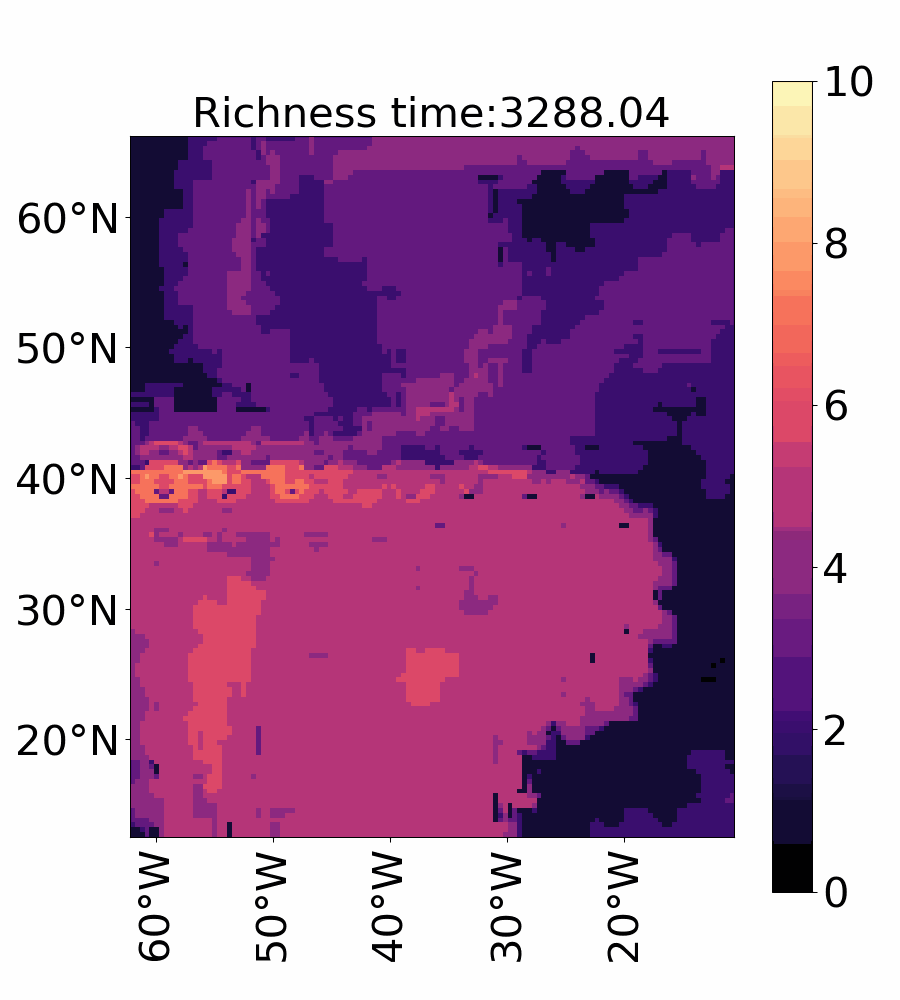

### Video_3d_richness_ibm_vref1.gif

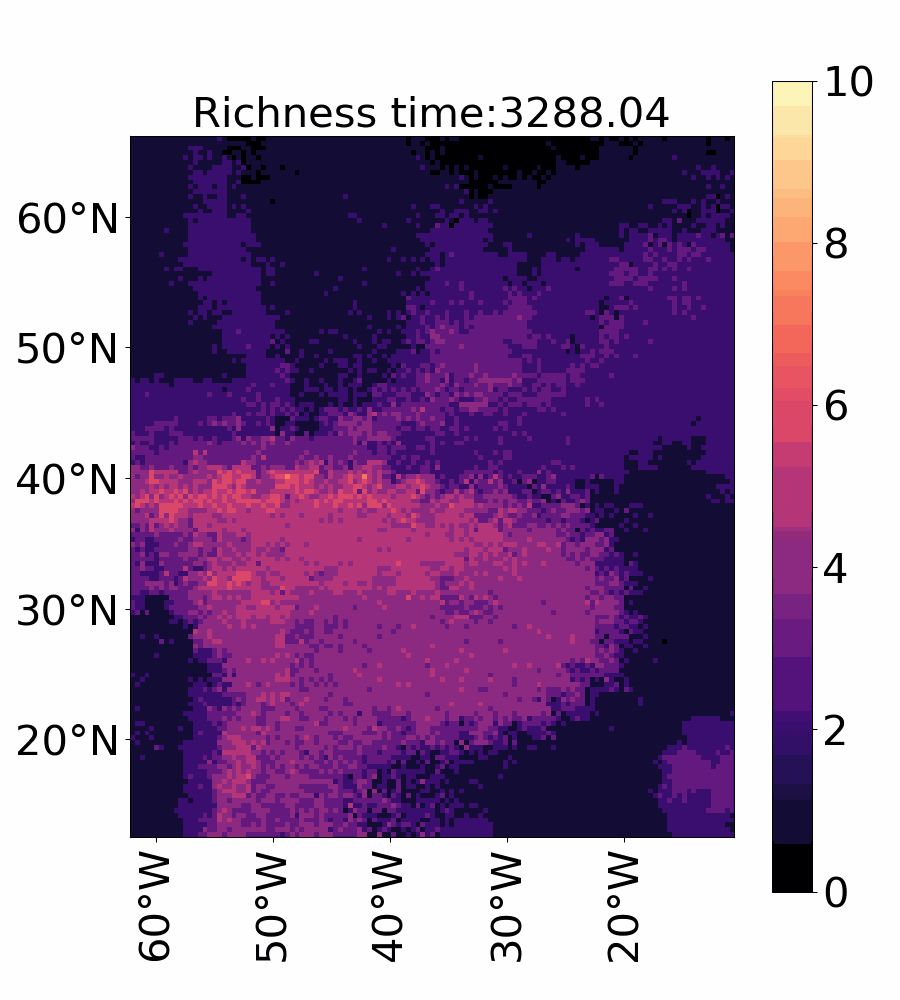

### Video_3e_richness_ibm_vref10.gif

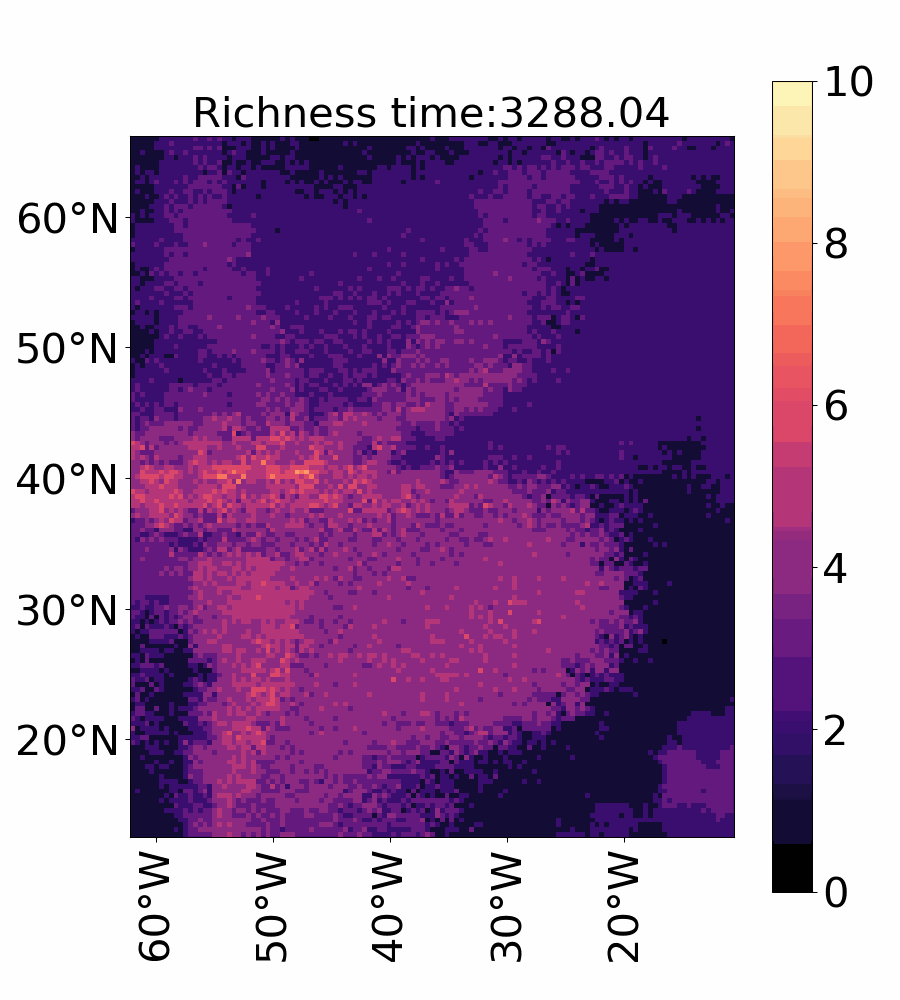

### Video_3f_richness_ibm_vref100.gif

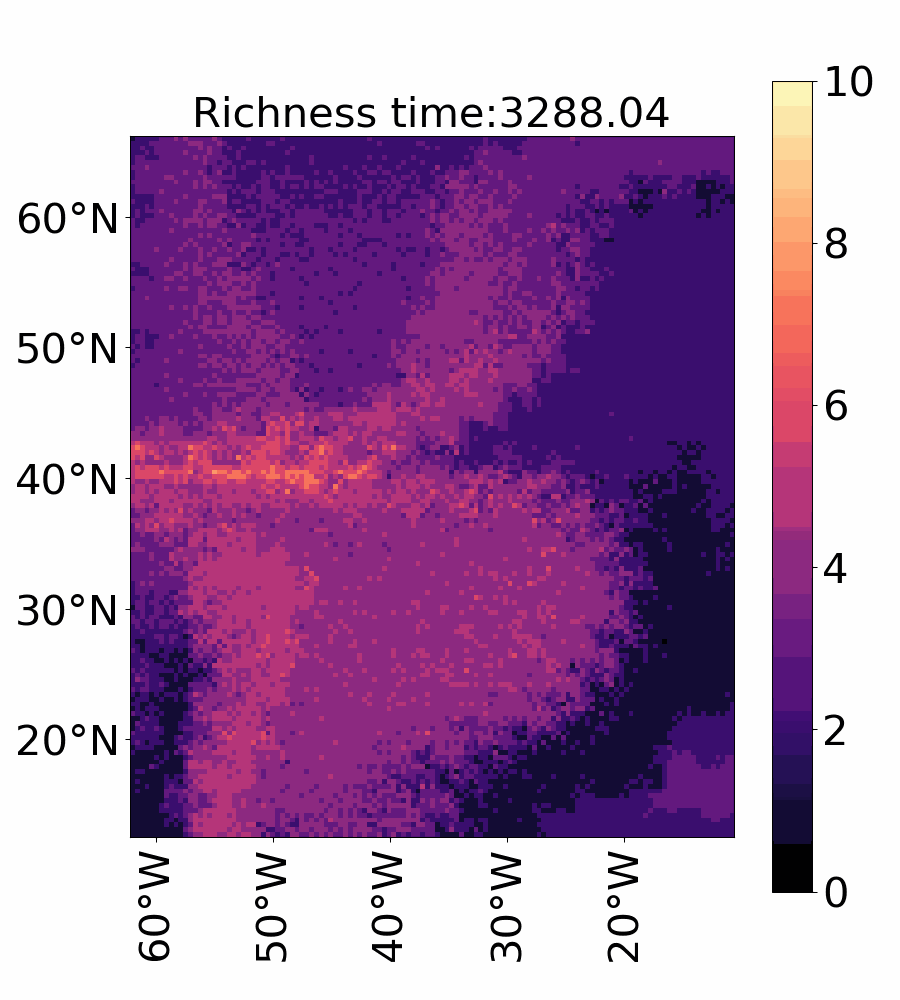

### Video_4a_shannon_euler_vref1.gif

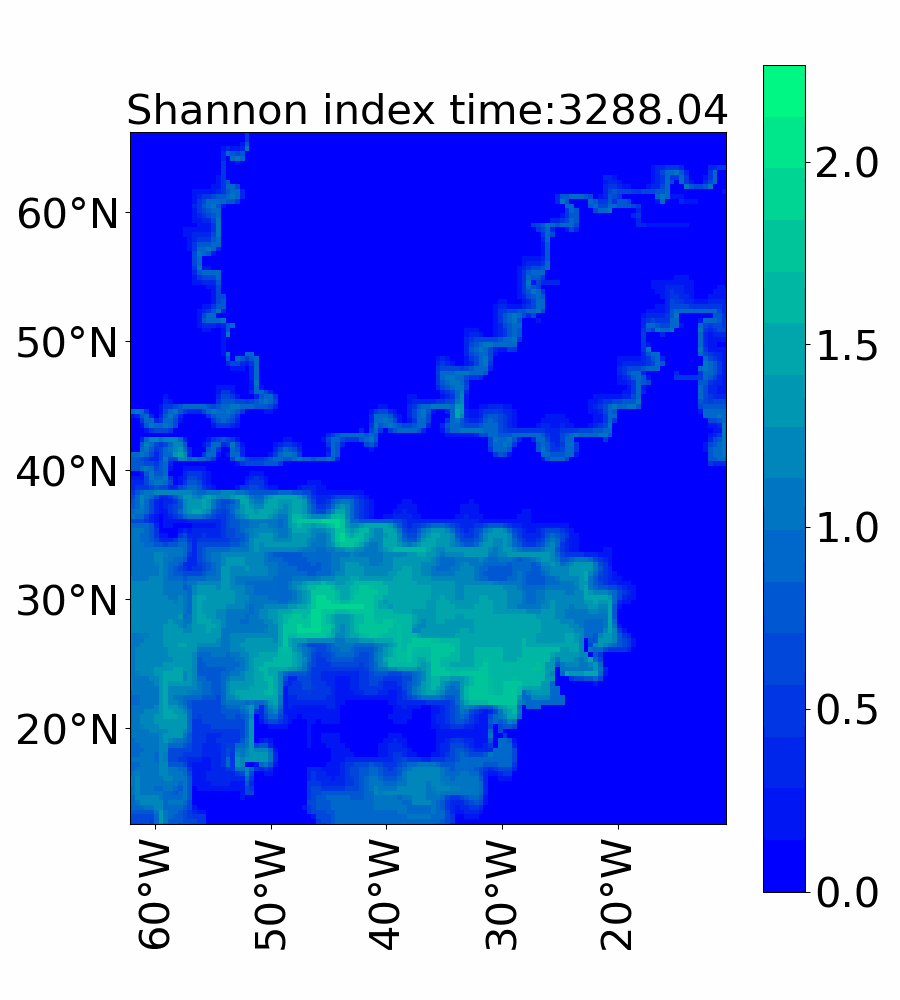

### Video_4b_shannon_euler_vref10.gif

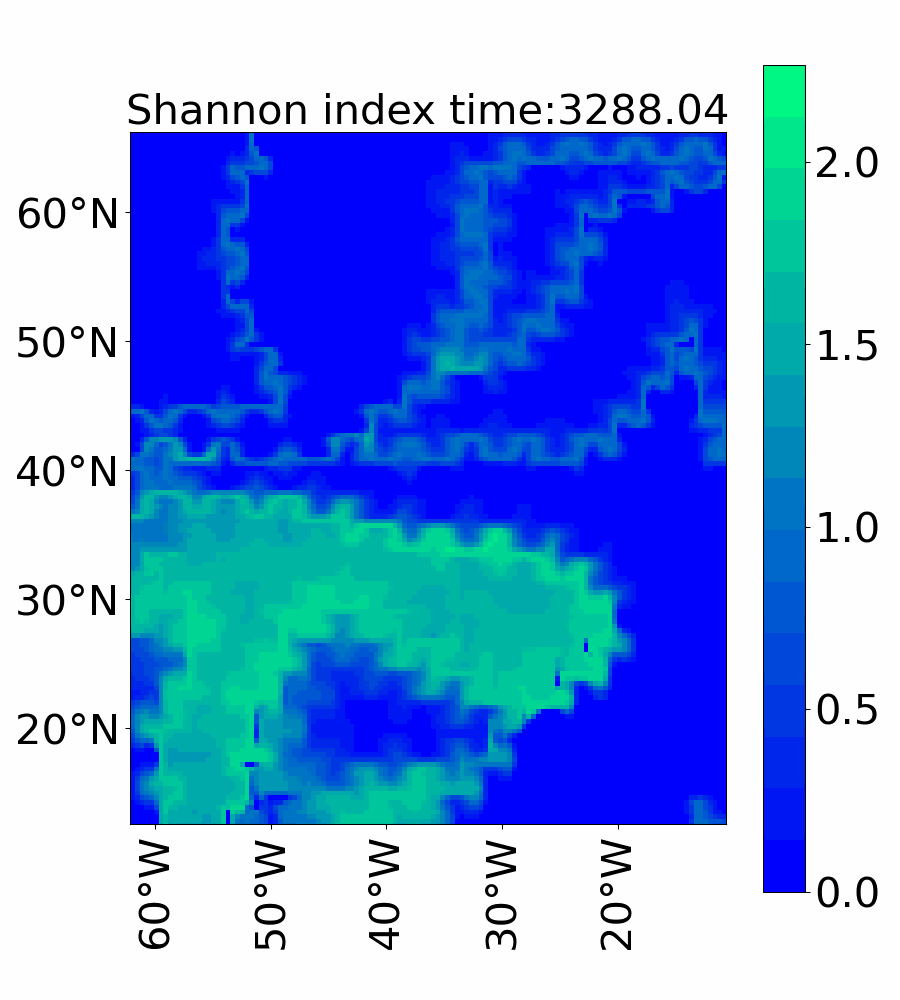

### Video_4c_shannon_euler_vref1.gif

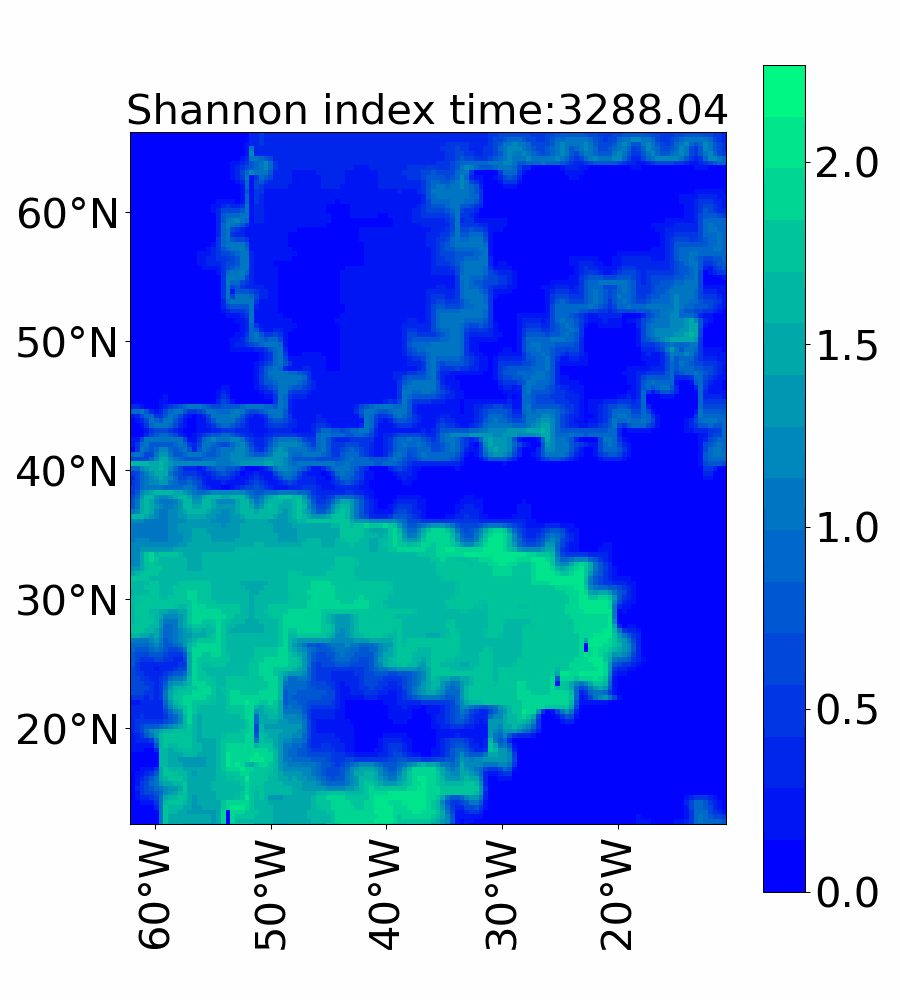

### Video_4d_shannon_ibm_vref1.gif

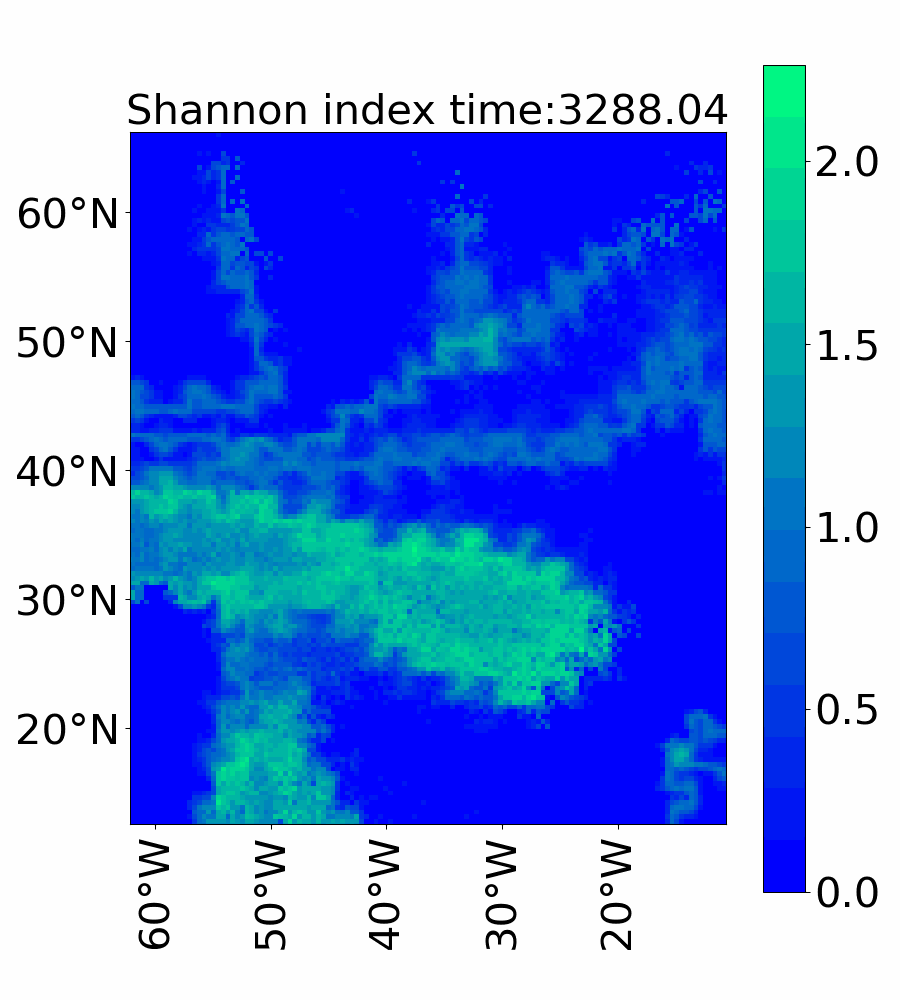

### Video_4e_shannon_ibm_vref10.gif

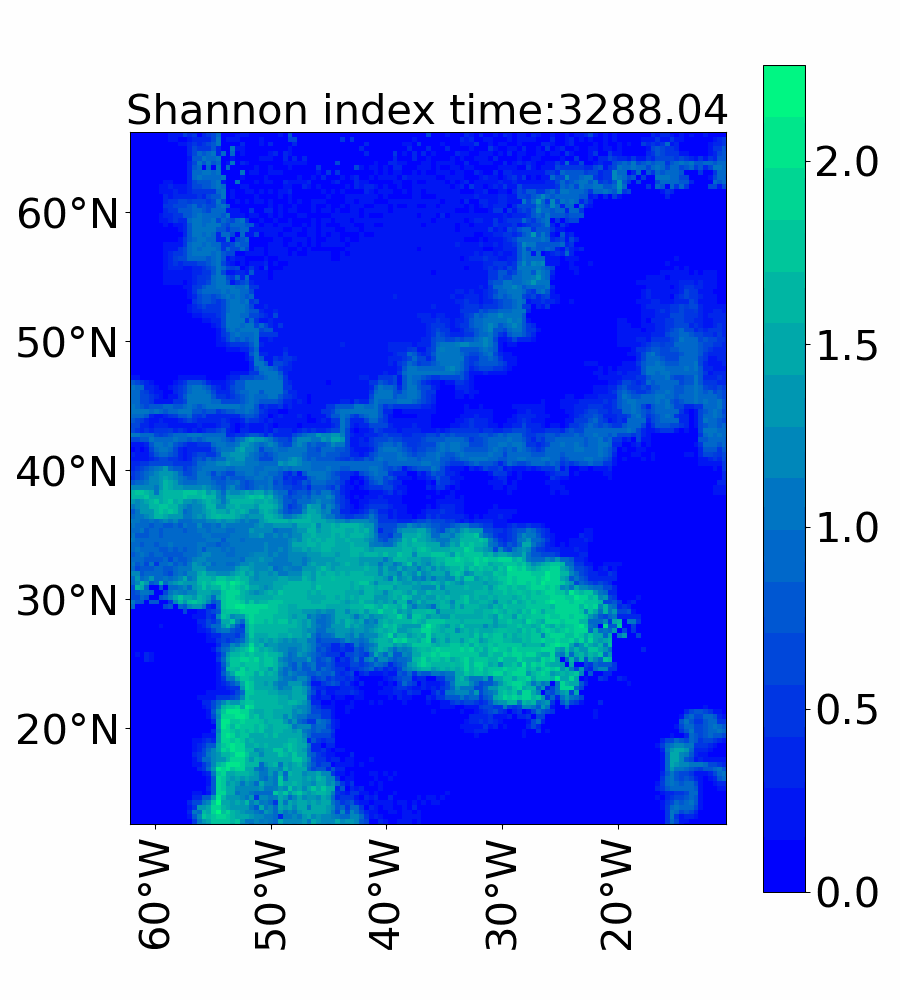

### Video_4f_shannon_ibm_vref100.gif

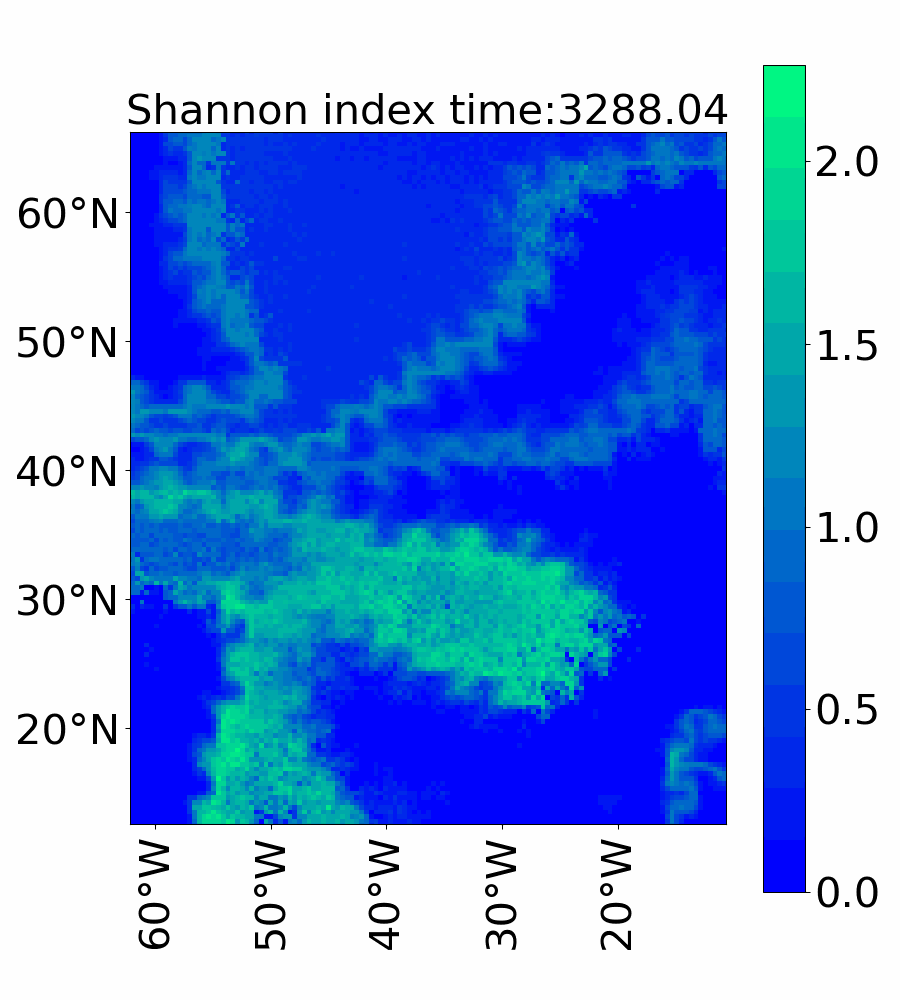

### Video_5a_dominant_species_euler_vref1.gif

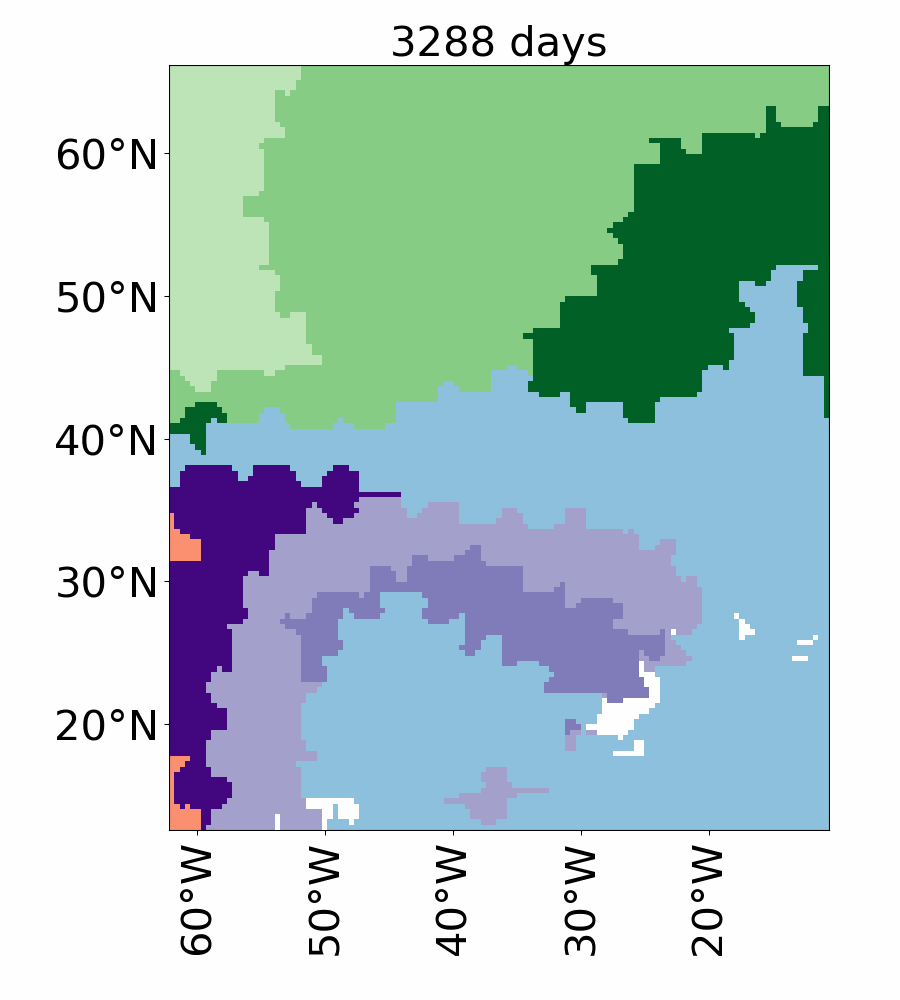

### Video_5b_dominant_species_euler_vref10.gif

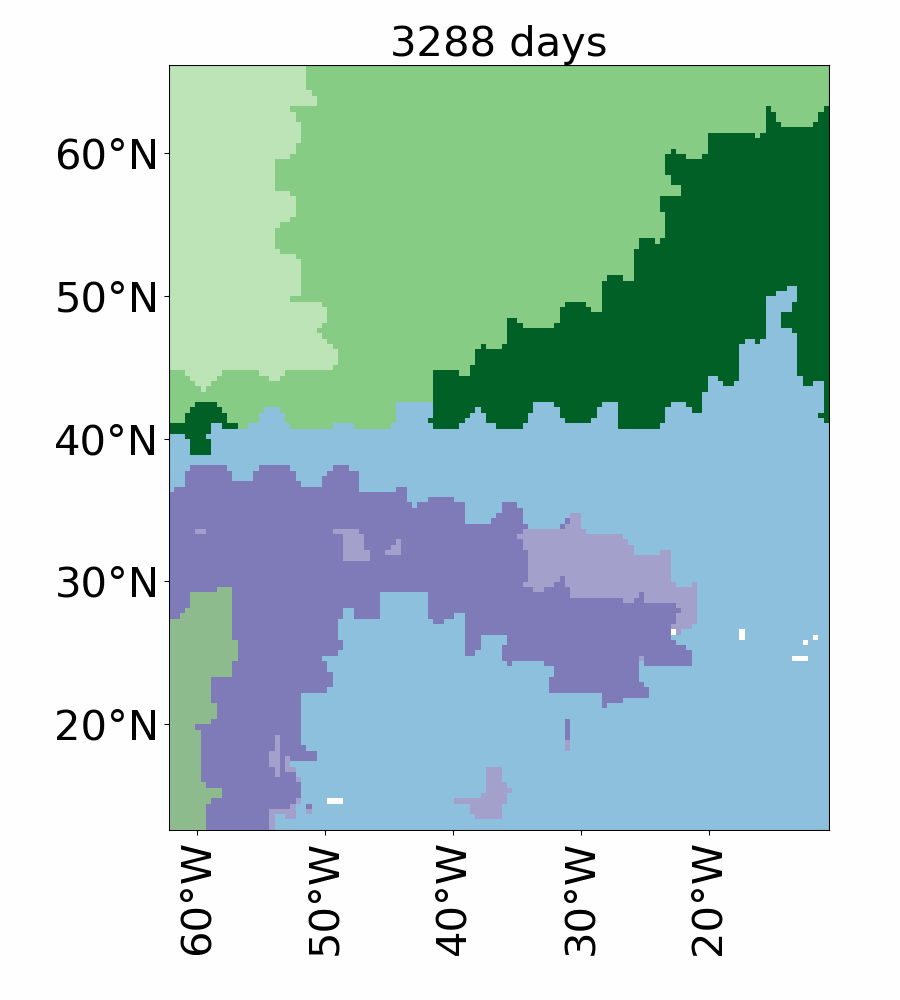

### Video_5c_dominant_species_euler_vref100.gif

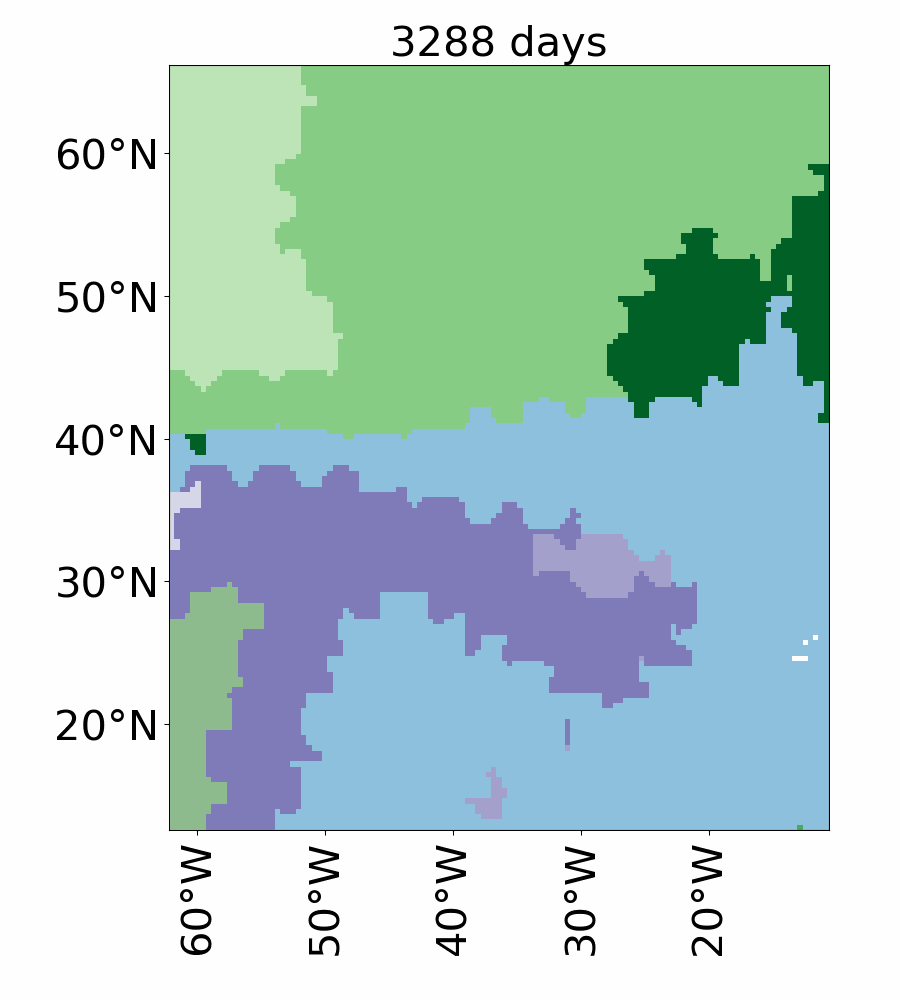

### Video_5d_dominant_species_ibm_vref1.gif

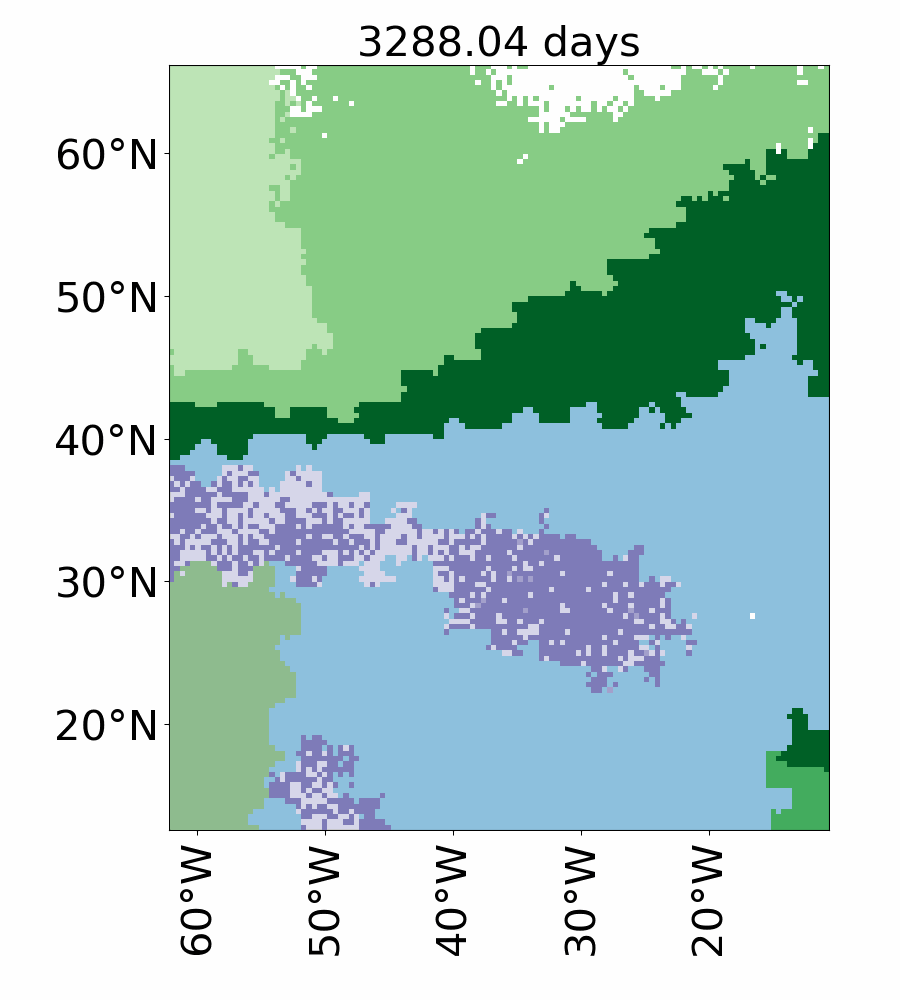

### Video_5e_dominant_species_ibm_vref10.gif

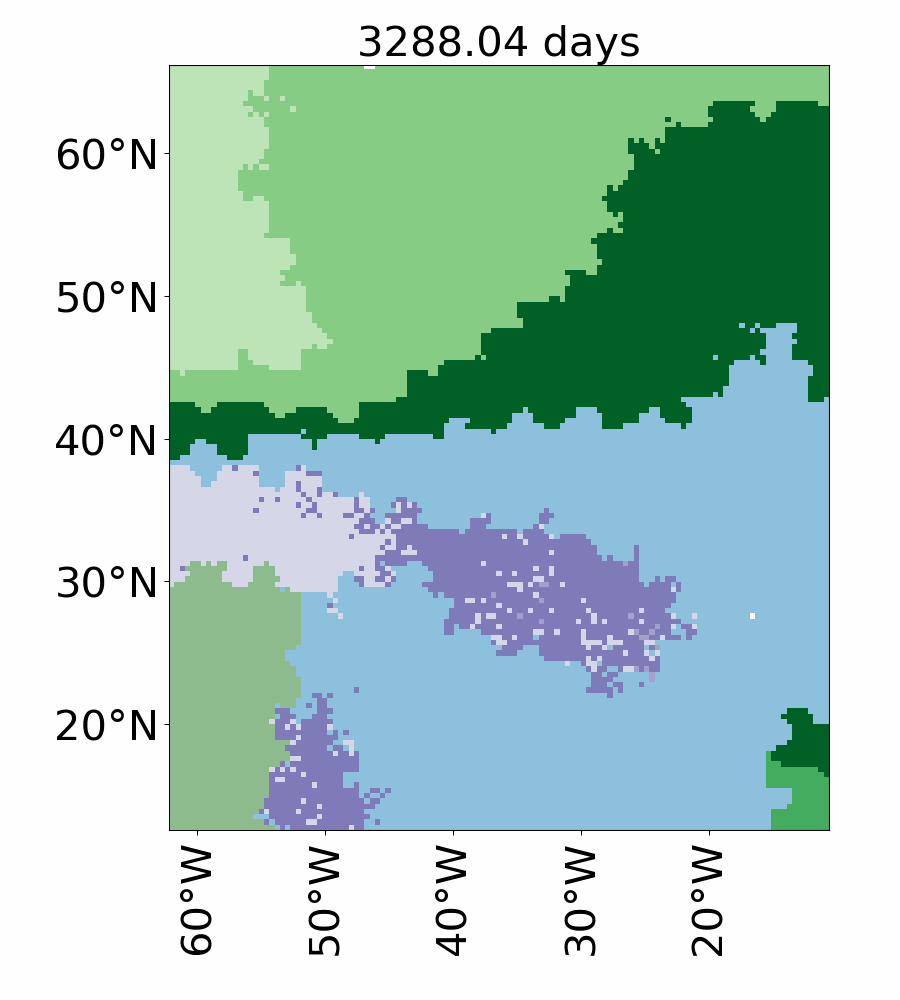

### Video_5f_dominant_species_ibm_vref100.gif

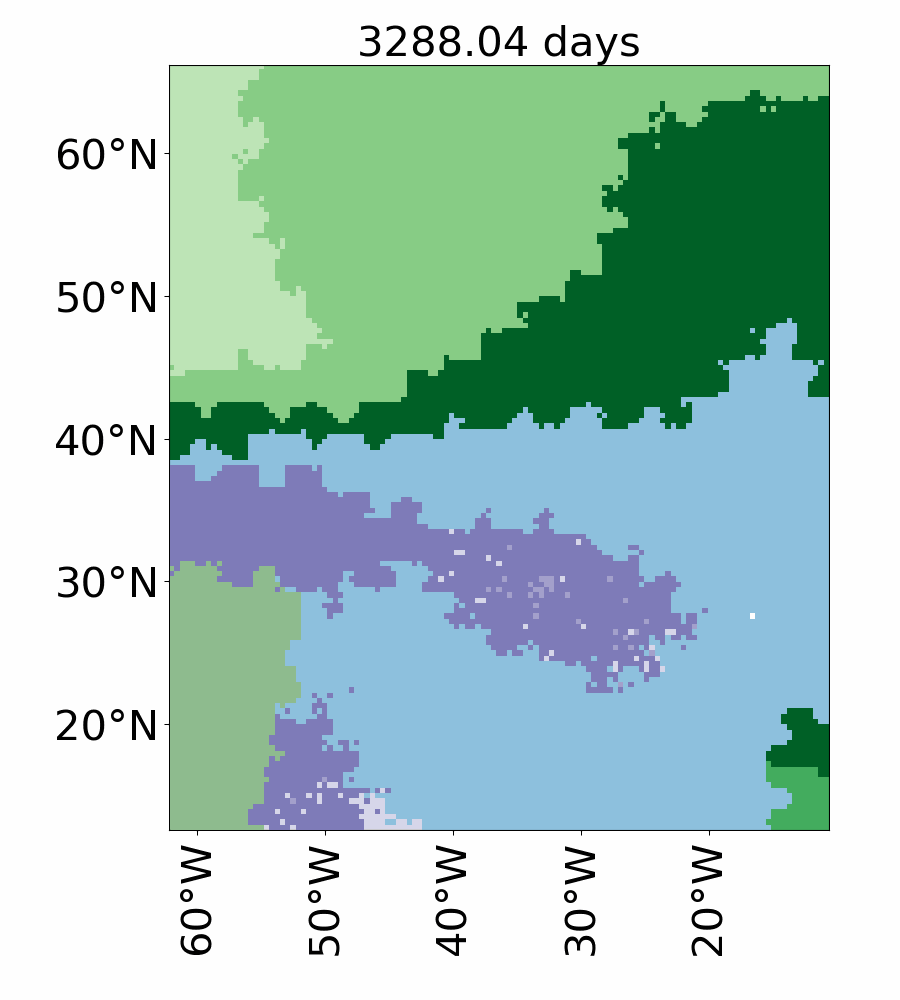
